# Deconstructing Complexity: A Computational Topology Approach to Trajectory Inference in the Human Thymus with *tviblindi*

**DOI:** 10.1101/2023.07.13.547329

**Authors:** Jan Stuchly, David Novak, Nadezda Brdickova, Petra Hadlova, Vojen Sadilek, Ahmad Iksi, Daniela Kuzilkova, Michael Svaton, George Alehandro Saad, Pablo Engel, Herve Luche, Ana E. Sousa, Afonso R. M. Almeida, Tomas Kalina

## Abstract

Understanding complex, organ-level single-cell datasets represents a formidable interdisciplinary challenge. This study aims to describe developmental trajectories of thymocytes and mature T cells. We developed *tviblindi*, a trajectory inference algorithm that integrates several autonomous modules - pseudotime inference, random walk simulations, real-time topological classification using persistent homology, and autoencoder-based 2D visualization using the *vaevictis* algorithm. This integration facilitates interactive exploration of developmental trajectories, revealing not only the canonical CD4 and CD8 development but also offering insights into checkpoints such as TCRβ selection and positive/negative selection. Furthermore, *tviblindi* allowed us to thoroughly characterize thymic regulatory T cells, tracing their development passed the negative selection stage to mature thymic regulatory T cells. At the very end of the developmental trajectory we discovered a previously undescribed subpopulation of thymic regulatory T cells. Experimentally, we confirmed its extensive proliferation history and an immunophenotype characteristic of activated and recirculating cells. *tviblindi* represents a new class of methods that is complementary to fully automated trajectory inference tools. It offers a semi-automated tool that leverages features derived from data in an unbiased and mathematically rigorous manner. These features include pseudotime, homology classes, and appropriate low-dimensional representations. These features can be integrated with expert knowledge to formulate hypotheses regarding the underlying dynamics, tailored to the specific trajectory or biological process under investigation.

## 1. Introduction

Exploring the development of mature, fully functional cells from their progenitors has been the focus of researchers for many decades. While single-cell techniques such as flow cytometry were fundamental to the description of developmental stages, recent advances in single-cell methods measuring large sets of markers (mass cytometry, single-cell RNA-seq) have enabled truly comprehensive descriptions of entire tissues and organs. In parallel, computational methods that capture and describe possible dynamics in the data, as well as topological relationships among cell populations, were developed. After the seminal Wanderlust methodological study (*1*), trajectory inference (TI) and pseudotime data analysis became an area of intensive research, producing gradually more and more sophisticated tools for a growing number of general topologies (*2–7*). Ideally, bioinformatic tools should reveal the structure of the data to an expert who can interpret it with a focus on his or her research question.

However, certain limitations prevent widespread adoption of these tools in analytical workflows of real-world data. First, the computational complexity of existing TI methods is often prohibitive when analyzing large datasets from current single-cell RNA methods (especially when the data comprise cell atlases of entire organisms, embryos, and human organs) or from flow and mass cytometry with datasets of millions of cells (*3,8–11*). Second, TI methods often use multiple steps of dimensionality reduction and/or clustering, inadvertently introducing bias. The choice of hyperparameters also fixes the a priori resolution in a way that is difficult to predict. Third, the current TI methods work well on artificial datasets but lack a straightforward approach to control the effect of noise (technical artifacts or unexpected events) in multiscale topologies of real-world data. The fourth, and perhaps the major obstacle is that existing TI tools do not offer the necessary interaction with the analytical process. We believe that such interaction, which helps the researcher understand the particularities of a sample, is crucial for exploratory analyses of unknown data.

Motivated to develop a generic TI solution useful to a biologist investigating a real-world dataset, we developed *tviblindi. tviblindi* is a modular TI method designed to tackle large datasets, significant noise, technical artifacts, and unequal distribution of cells along the time axis. As a proof of concept, we investigate αβ T-cell development in human thymus and the transition of mature T cells to peripheral blood.

Human T cells develop in the thymus, which is seeded by bone marrow derived CD45^pos^ CD44^pos^CD34^hi^CD7^neg^ and Notch primed CD45 ^pos^CD44^pos^CD34^hi^CD7^pos^ progenitors (*12–15*). Large CD34^hi^CD1a^neg^ immature thymocytes develop into small CD34^dim^CD1a^pos^ immature thymocytes (*13*), becoming gradually restricted to the T-cell lineage (*16*). After losing CD44 expression, right before gaining CD1a, they become committed to a T-cell fate (*17*).

These early human CD34^pos^ thymocytes then progress through three stages (*18*) (1) CD34^pos^CD38^neg^CD1a^neg^, (2) CD34^pos^CD38^pos^CD1a^neg^ and (3) CD34^pos^CD38^pos^CD1a^pos^. The first TCRβ D-J rearrangements are detected in the CD34^pos^CD38^pos^CD1a^neg^ population. In-frame TCRβ are mostly selected at the transition from CD34^pos^CD38^pos^CD1a^pos^ to the next stage: the CD4^pos^ immature single positive (ISP) population. Concordantly, pre-TCRα (pTα) expression peaks in the CD34^pos^CD38^pos^CD1a^pos^ and ISP stages, after which it declines. TCRα rearrangements are initiated when thymocytes progress from CD34^pos^CD38^pos^CD1a^pos^ toward the ISP stage and continue until the CD3^pos^CD4 and CD8 double positive (DP) stage (*18*).

The complete TCRαβ receptors are checked for their binding to MHC-self peptide complexes on thymic epithelial cells and dendritic cells. Failure to engage the MHC-self peptide leads to death by neglect (*19*). Efficient TCR binding leads to positive selection and further development along the CD4/CD8 axis. Positively selected DP cells express CD69 on their surface and start to downregulate the CD8 co-receptor. If they are selected via the MHC-II peptide complex, the TCR signal persists. When the duration of this signal exceeds a certain time limit, the transcription factor ThPOK is induced, and the cells become CD4 cells (*20*). If selected on the MHC-I peptide complex, the signal diminishes, the transcription factor RUNX3 is induced, expression of the CD8 co-receptor is reactivated and CD4 eliminated, leading to the CD8 cell fate (*20*). On the other hand, strong TCR binding leads to negative selection and apoptosis or to the so-called agonist selection and development of regulatory T cells (Tregs) (*21*). During these processes the cells acquire the chemokine receptor CCR7 and move from the thymic cortex to the medulla, where tissue-specific self-antigens are ectopically expressed on the surface of medullary epithelial cells (*22,23*). The current models suggest that both the strength/duration of TCR signal (*24–27*) and the integration of TCR signals from consecutive T cell – antigen-presenting cell encounters (*28,29*) help to determine the autoreactive T-cell fate. Costimulatory signals (*30–32*) as well as cytokines (*33–39*) contribute to the process. The upregulation of the high-affinity IL-2 receptor CD25 and its signaling are known to stabilize FOXP3 expression (*36,37*). Importantly, JAK/STAT signaling triggered by γ chain cytokines promote FOXP3^pos^ Treg proliferation (*38*) and induction of BCL-2 and its anti-apoptotic effect (*39*). While the TCR repertoire of conventional T cells and Tregs are largely distinct and non-overlapping (*40*), the exact branching point of these two main lineages is still a matter of intensive research (*41,42*) also employing human data based on single-cell transcriptomics (*43–45*).

When investigated using single-cell methods (*15,43–48*), the process of T-cell development translates into high-dimensional single-cell data with complex topology which need to be interrogated computationally.

We developed *tviblindi,* a new tool which breaks new ground in several aspects:

(1) It offers a highly scalable, linear complexity framework that works at a single-cell level.
(2) Analysis is performed in the original high-dimensional space, avoiding artifacts of dimensionality reduction.
(3) The framework is adapted to discover features taking into consideration their varying scales (along the time axis or in different cell types).
(4) Crucially, our method allows the user to interact with the analytical process and to set the appropriate level of resolution in real time.

In this paper, we first describe our new computational approach to exploration of developmental trajectories. Then we present an analysis of real-world datasets of T-cell development to illustrate the power of our analytical method and to describe the development of human T regulatory cells in detail. Last, we describe novel discrete stages in the development of Tregs and cell surface markers used in their study.

## 2. Results

### 2.1. Computational method for the TI and interrogation – *tviblindi*

The objective of *tviblindi* is to offer a flexible framework for TI and topological data analysis (TDA) interrogation of single-cell data. *tviblindi* facilitates efficient interpretation of data as well as sensitive discovery of minor trajectories. These two competing goals are resolved by interactive grouping of random walks, which are used as probes in the high-dimensional space, into trajectories, allowing for real-time adjustment of trajectory-specific resolution. This interactive approach provides an exhaustive description of the data, while allowing the researcher to introduce expert knowledge into the analytical process and to gradually gain insight into a particular dataset.

*tviblindi* was designed as an integrated framework implementing several independent modules to facilitate the use of diverse approaches to pseudotime estimation and data visualization, without the need to rerun the analysis. *tviblindi* allows to compute pseudotime which is resilient to unequal distributions of cells (e.g., accumulation of double positive T cells during T-cell development or in case of the presence of a developmental block). It also keeps the resolution on the single-cell level, while processing data in a reasonable time frame. Random walks, directed by the pseudotime, are then simulated, thus creating a set of probes in the high-dimensional space. To assemble random walks into meaningful trajectories, *tviblindi* employs witness complex triangulation (*49*) to capture the topology of the original data without the need for dimensionality reduction. It implements a representation of homology classes, which accounts for incomplete coverage of the high-dimensional points by the triangulation, navigating around the potential pitfall of this computational approach.

Contrary to other approaches, *tviblindi* considers each random walk a separate entity, which allows for probing major as well as minor trajectories while keeping track of the feature dispersion along the time axis. This approach enables sensitive detection of irregularities, such as hubs and discontinuities, in the topology of the underlying graph. While pseudotime captures local geometry of the data, persistent homology (*50*) aims to capture significant non-local features such as sparse regions (holes), which persist over a range of scales. In simplified terms, the persistence of a sparse region may be understood as the difference between the density of points at its boundary and in its interior.

Most persistent features can be extracted and used to capture the differences between random walks, inducing a multiscale clustering (*51*). We generalized and extended this idea to high-dimensional data. We took advantage of the natural hierarchical structure of persistent homology classes to achieve real-time, on-demand aggregation of random walks into trajectories respecting both local geometry and global topology of the point cloud.

#### 2.1.1. Introducing tviblindi on an artificial data set

We showcase our methodology using an artificial dataset created with the *dyntoy* package (for further details see Supplementary note, subsection 1.1 *dyntoy* dataset). Figure 1A shows a schematic representation of the data, capturing the basic topology with sparse regions labeled α and β.

**Figure 1:**
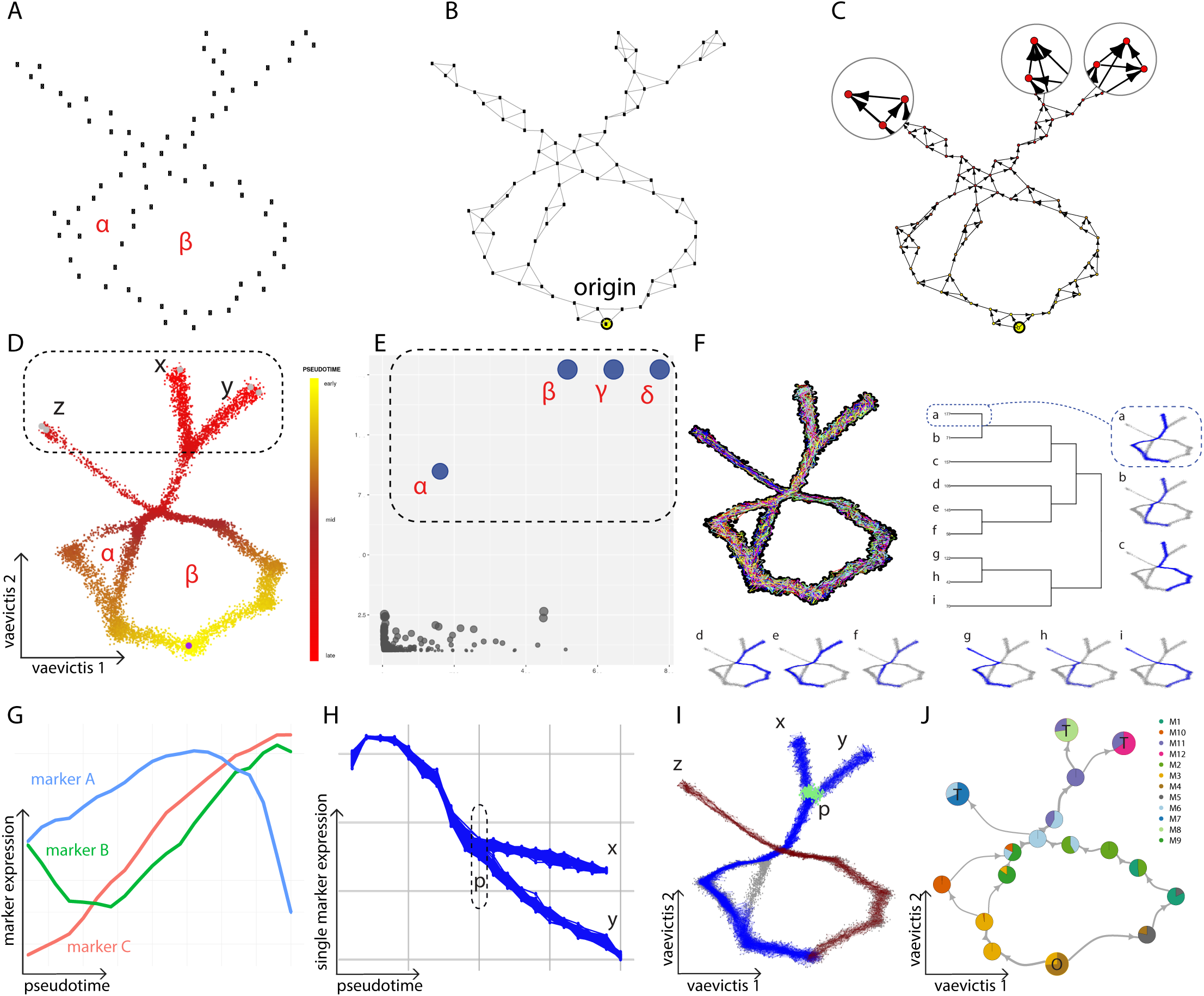
Analytical process on the artificial *dyntoy* dataset. Dashed lines mark selections made by the user, panels A-C show a scheme of the whole dataset, panels D-I correspond to the interactive GUI for this dataset: (A) The scheme of a point cloud representing single-cell data, where sparse regions α and β are present. (B) The dataset is represented as a k-NNG and the cell of origin is selected. (C) Pseudotime is calculated as expected hitting distance and the edges of the k-NNG are oriented according to the pseudotime. Candidate endpoints are automatically suggested as vertices without outgoing edges (circled). (D) Estimated pseudotime is represented by a yellow-to-red color gradient, the origin is shown as the purple dot. Potential developmental fates (x, y, z) are shown as gray points on the *vaevictis* plot. Endpoints can be selected for further investigation (in this example, all endpoints are selected, dashed line). (E) Sparse regions are represented on a persistence diagram, which enables the selection of the significant sparse regions (dashed line). The sparse regions γ and δ correspond to essential classes created by the selection of disparate endpoints. This way, random walks are classified using the selected ends and sparse regions. (F) All random walks are shown on the *vaevictis* plot and their classification into trajectories is depicted in the hierarchical clustering dendrogram. (G) By selecting branch (a) leading to the endpoint (x) of this dendrogram, the given trajectory can be investigated in detail, including viewing the average expression of multiple markers along pseudotime. (H) By selecting two endpoints (x and y) but only a single relevant marker, its dispersion and possible branching points can be examined. Trajectories (a and e) from panel F are visualized and the branching region (p) is selected. (I) Multiple trajectories (trajectory i in red and a and e in blue) can be visualized on the 2D plot. Points in the selected branching region (p) are highlighted in green. (J) Connectome representing a basic structure of the data and the simulated random walks. The pie charts indicate the distributions of cell populations in each vertex (cluster). The arrows show the direction of pseudotime. Vertex containing the cell of origin (O) and vertices containing endpoints (T).

First, the data are represented as an undirected k-nearest neighbor graph (k-NNG; Figure 1B). The cell of origin is defined by the user (either directly, or a population of origin is specified and the point closest to the centroid of this population is then taken) and used to estimate pseudotemporal ordering (pseudotime) directed away from the origin (Figure 1B). Edges of the k-NNG are oriented with respect to the pseudotime forming a directed acyclic graph (DAG).

Next, a large number (typically thousands) of random walks is simulated on the DAG. These walks are finite and their final vertices are candidates to become developmental endpoints (Figure 1C). The user can select one or more potential endpoints for further investigation (Figure 1D labeled x, y, z).

A persistence diagram, which captures the prominence of sparse regions within the point cloud, then allows the user to select significant sparse regions (Figure 1E), and random walks are organized into trajectories by means of hierarchical clustering. This clustering respects the global geometry and classifies trajectories based on how they navigate around the sparse regions (Figure 1F). In the presented artificial dataset, *tviblindi* correctly identified three putative endpoints (x, y, z) and organized simulated random walks into three distinct trajectories for each endpoint (a, b and c, for end x; d, e and f for end y and g, h and i for end z).

Our interactive framework allows the user to inspect the evolution of specific markers (Figure 1G), track key points in their development (e.g., branching point p) and focus on cells at such key points (Figure 1H, I). For a quick overview, a “connectome” summarizing the basic structure of the point cloud and of simulated random walks can be plotted (Figure 1J). We describe each *tviblindi* module below. Detailed algorithmic descriptions of particular modules can be found in the Supplementary note, sections 2-4.

#### 2.1.2. Visualization

Visualization of high-dimensional data is essential for the initial overview and for the informed interactions with the data during downstream analysis. However, any dimensionality reduction introduces simplification, which can lead to misinterpretation. In *tviblindi* framework, the dimensionality reduction step is independent of all other modules (which work directly in the original high-dimensional space) and is used solely for a visual representation of the dataset.

While dimensionality reduction has been extensively and successfully used for single-cell data (e.g., UMAP (*52,53*) and viSNE (*54,55*)), interpreting developmental trajectories requires not only the local, but also the global relationships of cells to be captured (often distorted in the lower-dimensional embedding by current dimensionality reduction algorithms). For this purpose, we created *vaevictis*, a variational autoencoder model based on ideas in *scvis* (*56*) and *ivis* (*57*) fitted with naive importance sampling. This technique creates a continuous representation of the data rather than isolated islands of populations (Supplementary note, section 2 Dimensionality reduction, Figure 1B, C). Due to the naive importance sampling step, numerically large compact populations are not overrepresented. This way, we obtain a clearer picture of the developmental dynamics in the data (Supplementary note, section 2 Dimensionality reduction, Figure 1D, E). Moreover, once calculated, it can be applied to new datasets with the same measured parameters (protein markers or detected transcripts), facilitating integration of new results.

#### 2.1.3. Pseudotime and random walks simulation

Correct pseudotemporal ordering is an essential and computationally intensive part of TI algorithms. Here we use a particular formulation (and modification) of the idea of *expected hitting time* (*58*). We calculate the expected number of random steps necessary to reach any cell in a dataset from the cell of origin. To leverage the sparsity of the k-NNG and to calculate the hitting times on a single-cell level, we use the following formulation: given an undirected k-NNG *G = (V, E, p*), with vertices *V*, edges *E*, weights *p*: *E* → [0,1] representing the probability of transition of each edge respectively, and an origin *v*_0_, the expected hitting time of a cell is a weighted average of hitting times of its neighbors increased by 1 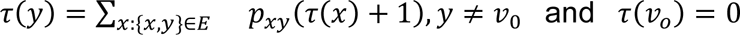. This formula translates into a sparse linear system, which can be solved efficiently by numerical methods (such as conjugate gradients) to any precision even for graphs with millions of vertices. Consequently, we are able to calculate the pseudotime directly in the original space at a single-cell level. Furthermore, we obtain a measure of success: the relative error reported by the numerical solver.

A key application of *tviblindi* is the comparison of trajectories between normally developing tissues and those with developmental abnormalities, manifesting a block and/or pathological proliferation resulting in an overabundant intermediate population. In such cases, the hitting time calculation would assign the highest pseudotime to this intermediate population, which would hamper the topology of simulated random walks (Supplementary note, section 3, Pseudotime estimation & random walks simulation). To mitigate this effect, the formula above can be modified in the following way: suppose that apart from *G* = (*V*, *E*, *p*), we have a sparse matrix *D*, recording mutual distances for all vertices in the graph connected by an edge. Then the definition of the pseudotime as the *expected hitting distance is* 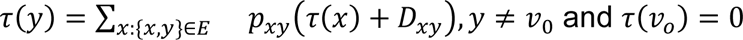.

In other words, the expected distance to travel before reaching vertex y. Both methods are also very robust with respect to the choice of the number of neighbors in the k-NNG (Supplementary note, section 8 Performance evaluation).

Once pseudotime is computed, the edges in the graph *G* = (*V*, *E*, *p*) get oriented (Figure 1B) and a large number of random walks is simulated on the DAG (Figure 1F). Importantly, all random walks on the graph are finite. The set of terminal vertices of these walks represents estimates of terminal biological fates (for further details see Supplementary note, section 3 Pseudotime estimation & random walks stimulation).

#### 2.1.4. Real-time interactive topological classification

The most distinctive feature of *tviblindi* is the interactive aggregation of simulated random walks into trajectories with respect to the global topology of the point cloud. This procedure is based on the idea of multi-scale topological classification using persistent homology (*59*). Random walks are considered distinct entities, and they can be aggregated into trajectories respecting the structure of the point cloud.

Walks are clustered based on how they circumnavigate significant sparse regions identified using persistent homology. The appropriate level of significance can be adjusted by the user by the selection of significant sparse regions on a persistence diagram (Figure 1E). Classification by the means of persistent homology has a natural hierarchical structure induced by filtration (*59,60*) (see Supplementary note, section 4 Topological clustering of random walks for details), which translates into a dendrogram of random walks (Figure 1F).

Applying classification by persistent homology to a large high-dimensional dataset with considerable technical noise requires several improvements of previously published work (*59*). First, as noted above, the selection of significant sparse regions is interactive, which allows the user to choose an appropriate level of detail for each dataset or trajectory. Second, we use witness complex triangulation (*61*) to adapt for high dimensionality of single-cell data (for which the original methodology was intractable). Third, since the witness complex triangulation does not guarantee filling the space completely, we introduce the detection and representation of sparse regions with infinite persistence. This last modification also allows us to classify random walks into trajectories based on their different endpoints and consequently to study developmental branching. (This is impossible with the original formulation.) See Supplementary Videos and Supplementary note for a presentation of the *tviblindi* graphical user interface (GUI).

#### 2.1.5. Connectome – a fully automated pipeline

For a quick overview of a given dataset, we have implemented a “connectome” functionality inspired by PAGA (*5*) (Figure 1J). This is a fully automated pipeline to create a directed graph, in which the data is clustered (by default using Louvain community detection (*62*)). Random walks are contracted to the identified clusters, providing a lower-resolution estimate of developmental trajectories. The clusters are plotted as pie charts of the represented cell populations. The graph layout employs the same 2D representation(s) as *tviblindi* GUI to facilitate the interpretation. The clusters containing the cell of origin and endpoints are automatically detected and denoted O and T respectively. At the same time the orientations and widths of edges reflect the direction and number of underlying random walks. Connectome suffers from some of the limitations of other, fully automated, end-to-end methods. The resolution is predetermined at the cell-population level by the choice of clustering. In addition, there is no direct way to interactively choose a resolution at the level of random walks, nor to detect and exclude clear artifacts. Therefore, we recommend using this functionality only to get an overview of a given single-cell dataset with suggested trajectories before performing the full interactive *tviblindi* analysis.

#### 2.1.6 Performance evaluation

We evaluated *tviblindi* against several state-of-the-art trajectory inference tools (Monocle 3(*63*), Stream(*64*), Palantir(*65*), Via(*66*), PAGA(*67*), CellRank 2(*68*) and StaVia(*69*)) and found it to be superior in terms of speed and resolution with respect to typical challenges. Specifically, it was better at dealing with locally abundant clusters, developmental loops, and non-dominant trajectories (shown and discussed in detail in Supplementary Note, Section 8). To further evaluate the performance of *tviblindi* on complex single-cell RNA sequencing data, we analyzed in detail the development of immune cells in the bone marrow of a published dataset(*70*). We also tested *tviblindi* on mouse gastrulation data(*71*) to assess its ability to interrogate large cell atlas datasets. We compared the results of *tviblindi analysis* of these datasets with the two best-performing methods (StaVia, CellRank2) from the performance evaluation. This comparison has shown that *tviblindi* is superior when dealing with distinct trajectories converging to a common fate (shown and discussed in detail in Supplementary Note, Section 10).

Sensitivity to hyperparameters is discussed in Supplementary Note, Section 8.1.

### 2.2. Using tviblindi on mass cytometry and single-cell RNA-seq data sets

#### 2.2.1. tviblindi exhaustively dissects human T-cell development

We applied *tviblindi* to the 34-parameter mass cytometry data obtained in our human T-cell development study (Supplementary Table Mass panel 1). The data contained barcoded human thymocytes and human peripheral blood mononuclear cells (PBMC). Dead cells and cells not belonging to the αβ T cell developmental lineage were excluded, yielding a total of 1,182,802 stained cells for analysis (Supplementary Figure 1). When the cells were visualized using a CD4xCD8 dot plot, we observed the expected patterns of double negative (DN), double positive (DP), CD4 single positive (SP) and CD8 SP populations for a cryopreserved thymus (Figure 2A). We then used the default *vaevictis* dimensionality reduction to reveal the basic structure of the T-cell compartment. Thymic and peripheral blood T cells were projected into distinct areas of the *vaevictis* embedding with a minor overlap (Figure 2B). Localization of the CD8, CD4 and Annexin V positive cells can be viewed on the plot using a heatmap color scale for expression levels of these markers. This provides an interpretable image of the T-cell compartment (Figure 2C).

**Figure 2:**
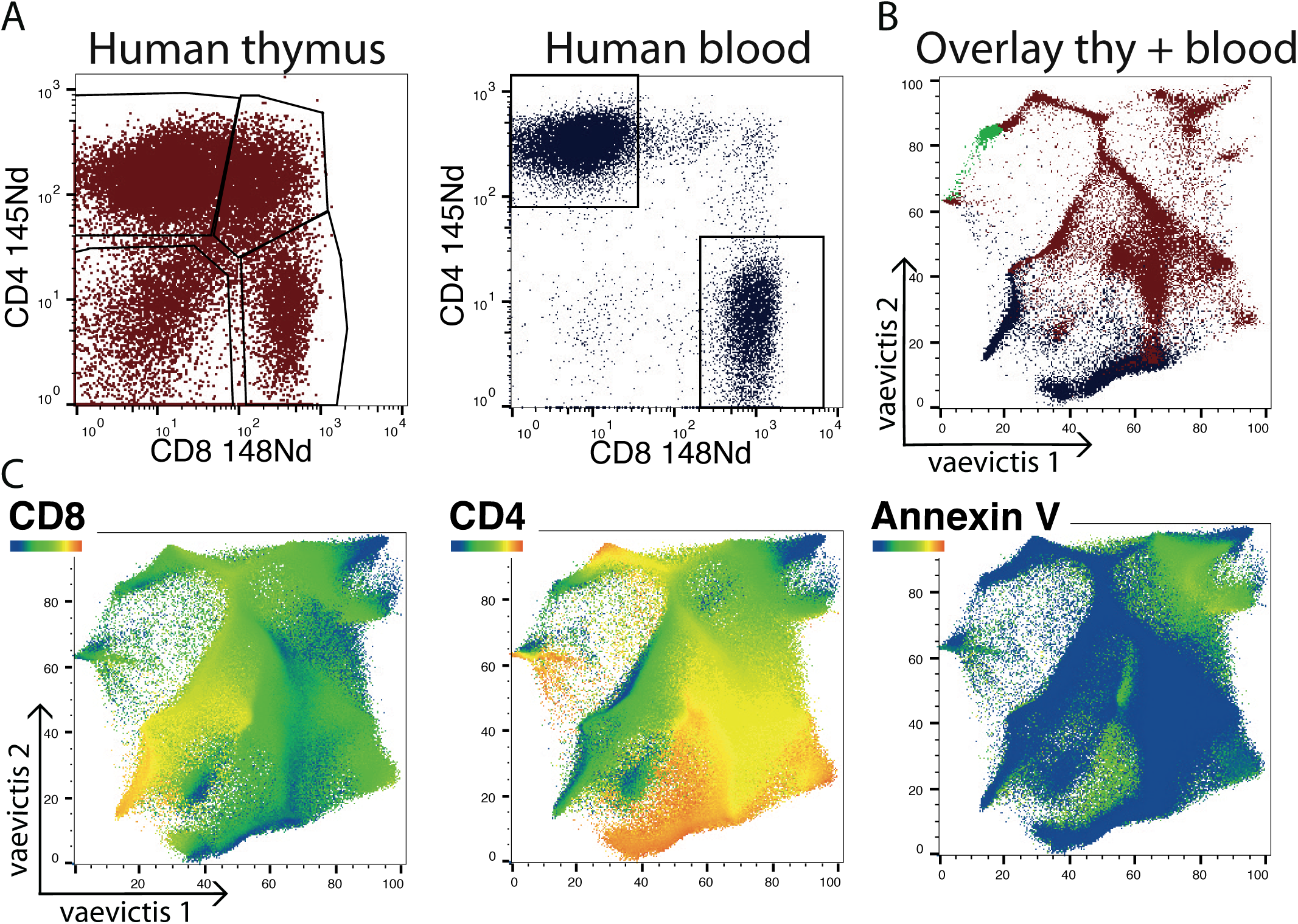
Visualization of the basic structure of the T-cell compartment using *vaevictis.* (A) Bivariate CD4 x CD8 dot plot of cells from human thymus (brown) and human peripheral blood (blue) acquired using mass cytometry. The cells were gated as DN, DP, CD4 SP and CD8 SP. (B) *vaevictis* plot of 37-parameter (including barcode) mass cytometry panel measurement showing the positions of human thymus (brown) and human peripheral blood (blue), with the CD34^pos^ progenitors shown in green. (C) Expression of CD8, CD4 and Annexin V shown using a blue-green-yellow-red color gradient on the *vaevictis* plot. Blue color indicates the lowest expression and red color indicates the highest.

Next, we gated the population of origin as CD34^hi^ CD1a^neg^ DN cells and analyzed the data with *tviblindi*. We simulated 7500 random walks and discovered eleven groups of endpoints labeled #1 - #11 (Figure 3A). Five of them (#1 to #5) were located in the region of peripheral CD4 T cells (where #5 corresponded to mature naive CD4 T cells transiting to peripheral blood). Endpoint #6 found among the thymic CD4 SP T cells is further described in the section *Non-conventional end interpretation.* Three groups of endpoints were located in the peripheral CD8 T cell compartment (where #7 corresponded to cells of CD8 TEMRA phenotype, #8 to CD8 central memory T cells and #9 to mature naive CD8 T cells transiting to the periphery). Two remaining endpoints (#10 and #11) belonged to the apoptotic thymocytes (compare to Figure 2B and 2C).

**Figure 3:**
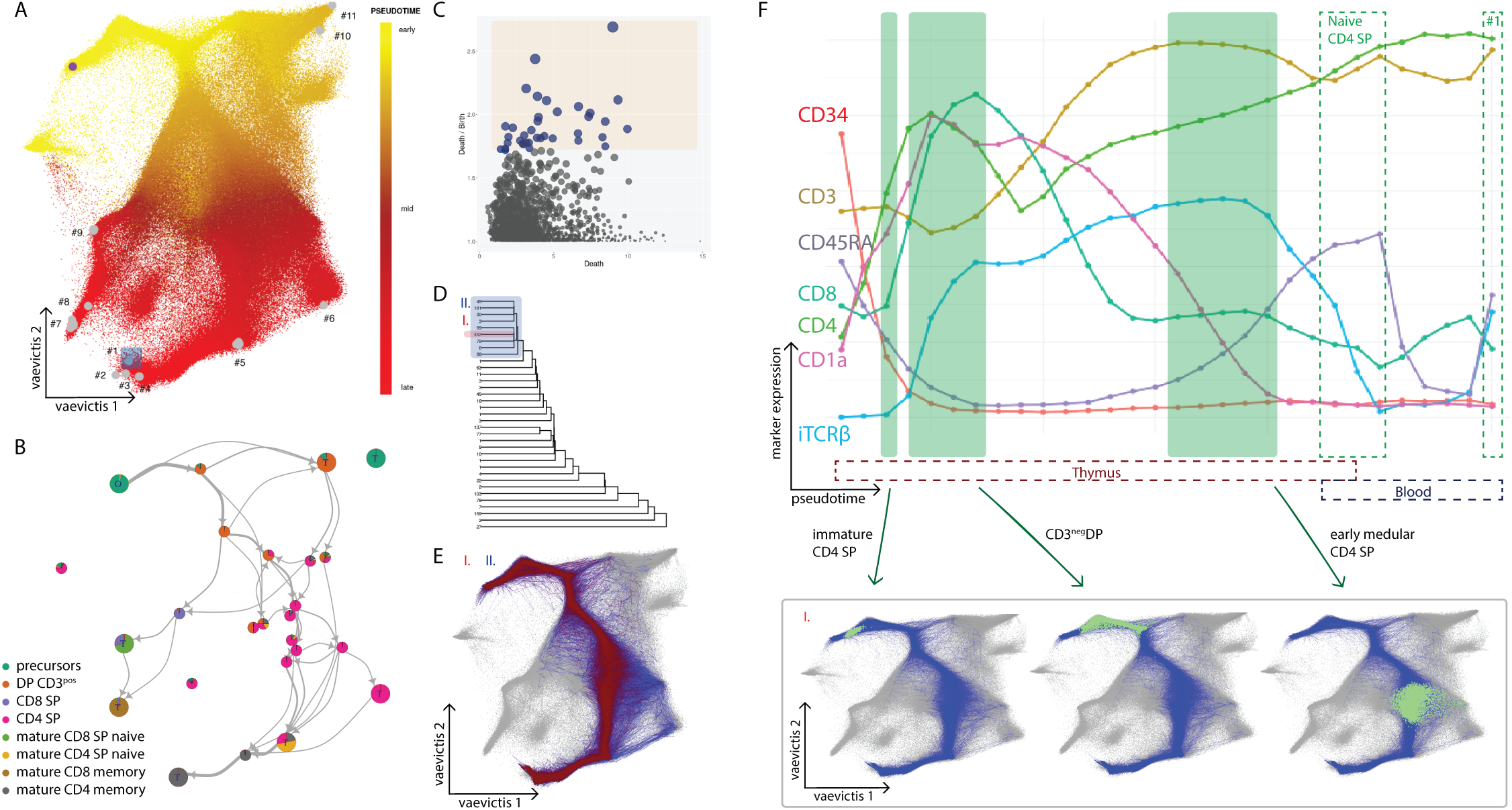
Developmental endpoints and detailed analysis of major trajectory leading to endpoint #1 by *tviblindi.* (A) *vaevictis* plot of T-cell development in thymus and peripheral blood. Estimated pseudotime is represented by a yellow-to-red color gradient with CD34^pos^ progenitors as the population of origin (purple dot). Gray dots indicate the discovered developmental endpoints. Endpoint #1 highlighted by a blue rectangle represents mature CD4^pos^ effector memory T cells selected for further exploration. (B) Connectome shows low resolution structure of the data and of simulated random walks. (C) Persistence diagram representing sparse regions detected within the point cloud of measured cells. The orange rectangle marks a user-defined selection of sparse regions. (D) Dendrogram of clustered trajectories shows a subcluster of 452 walks (red rectangle, labeled I) within a larger group of random walks (blue rectangle, labeled II). The number to the left of each leaf indicates the number of random walks in the leaf. (E) *vaevictis* plot displaying the above selected trajectories in corresponding colors. (F) Pseudotime line plot showing the average expression of selected individual markers along the trajectory to endpoint #1 (top). The selected areas of interest corresponding to T-cell developmental stages (green rectangles) are shown in green (as indicated by the arrows) on the *vaevictis* plots (below).

The connectome shows a basic overview of dynamics in the data on a cluster level (Figure 3B). The subsequent interactive analysis of simulated random walks gives biological meaning to the observed connections and allows for the detailed interpretation of data on a single-cell level.

First, we chose endpoint #1 representing peripheral CD4 effector memory T cells for further analysis. We applied persistent homology to select the significant sparse regions in the data and used these to cluster individual simulated random walks (Figure 3C). The topology of the point cloud representing human T-cell development is more complex than that of the illustrative artificial dataset shown in Figure 1 and does not offer a clear cutoff for the choice of significant sparse regions. Interactive selection allows the user to vary the resolution and to investigate specific sparse regions in the data iteratively.

Clustering the random walks resulted in a dendrogram, where leaves with abundant walks represent the dominant developmental trajectories. Nonetheless even the less abundant groups of walks can be selected for in-depth investigation (Figure 3D).

Leaf I. contains the largest number (452) of random walks (Figure 3D and 3E). Of note, leaf I. faithfully represents a larger supercluster of random walks II. (Figure 3D and 3E). For the leaf I. trajectory, we displayed the intensity of expression of key markers along the calculated pseudotime, the pseudotime lineplot (Figure 3F top). We then mapped the key developmental stages onto the *vaevictis* plot (Figure 3F bottom).

The CD4 ISP stage was found where CD4 and CD1a expression begins, but prior to CD8, intracellular TCRβ and CD3 expression. This is followed by a CD3^neg^ DP stage, where intracellular TCRβ increases. Eventually, the DP cells become CD3^pos^, reflecting the successful expression of the TCRαβ receptor. Next, they lose CD8 and later they lose the cortical thymocyte marker CD1a. Finally, the CD4 SP thymocytes gain CD45RA and are ready to egress from the thymus as naive CD4 SP cells. In the peripheral blood they lose the CD45RA during maturation only to gain it at their terminal stage (Figure 3F). Similarly, if endpoint #7 is chosen, the canonical trajectory leading to peripheral CD8 cells can be visualized and described (Supplementary Figure 2).

This knowledge of the topology of key points of canonical development helped us to interpret additional developmental endpoints and the trajectories leading to them.

#### 2.2.2. Variants of trajectories including selection processes

We then focused on alternative trajectories leading to peripheral CD4 cells corresponding to endpoint #1. Leaf IV., comprising 103 walks (Figure 4A) exhibits a clear artifact connecting highly dividing progenitor cells with doublets, which could invalidate a fully automated analysis (Supplementary Figure 3).

**Figure 4:**
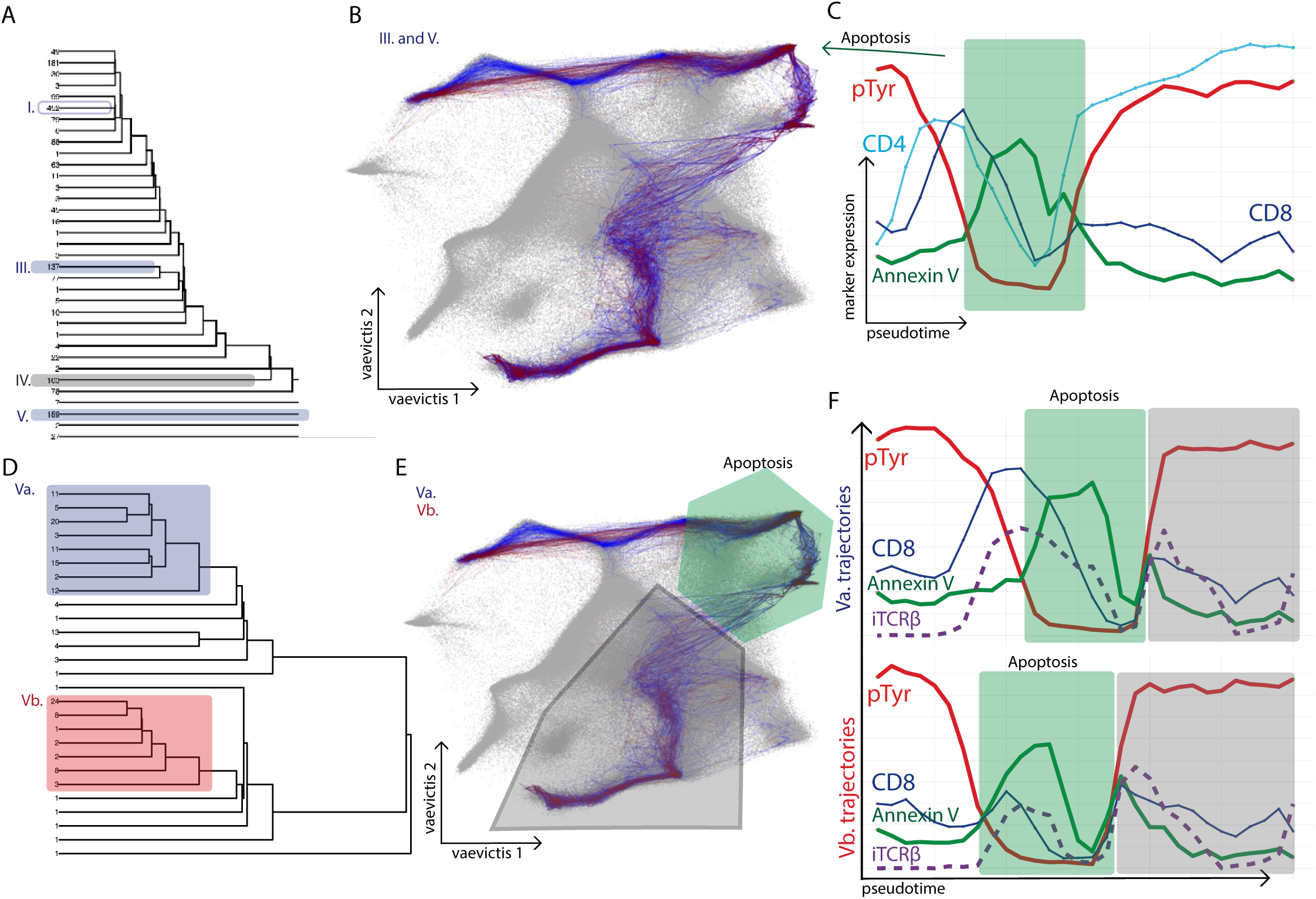
*tviblindi* analysis of trajectories leading to apoptosis. (A) Dendrogram identical to Figure 3D with additional trajectories selected for closer investigation (blue rectangles) labeled III (137 random walks) and V (159 random walks). (B) *vaevictis* plot showing the topology of trajectories in leaves III and V. Apoptotic cells are shown in green. (C) Pseudotime line plot, which shows the average expression of selected markers along calculated pseudotime. The green rectangle highlights the region with increased expression of apoptotic marker Annexin V and decreased expression of phosphotyrosine. Events in the selected region are displayed in panel 4B in green (see the arrow). (D) A detailed dendrogram of leaf V from panel 4A. Two distinct trajectories were selected for further analysis (blue rectangle, Va and red rectangle, Vb). (E) *vaevictis* plot showing the topology of trajectories in leaves Va and Vb. The green polygon marks the region of apoptosis, the gray polygon marks the region of more advanced stages of thymic and peripheral CD4 T-cell development. (F) Pseudotime line plot, which depicts the average expression of selected markers along the trajectories Va and Vb. The region of apoptosis and the region of more advanced stages of CD4 T-cell development are highlighted (green and gray rectangle respectively).

For further in-depth investigation we selected the two remaining larger groups of random walks represented in the dendrogram by leaves highlighted as III. and V. (Figure 4A). After the CD3^neg^ DP stage, these trajectories diverted from the canonical course described above. The cells passed through a region with an overall decrease in phosphorylated tyrosine expression and an increase in Annexin V expression, indicating apoptosis (Figure 4B and 4C). Of note, endpoints #10 and #11 were situated in this apoptotic region (see Figure 3A for their precise location). We therefore interpreted the first part of these trajectories as leading to apoptosis upon failure of DP thymocytes to pass either positive or negative selection checkpoints in the thymic cortex (Figure 4B and 4C).

Trajectories grouped in the leaves III. and V. continued through the apoptotic stage (indicated by a green hexagon in Figure 4E) and reconnected to the canonical CD4 T-cell maturation trajectory (indicated by a gray hexagon in Figure 4E). While this is an accurate description of the topology of the single cell data points, expert interpretation finds it counterintuitive since apoptotic death of thymocytes is the biological end-point for the unselected thymocytes. t*viblindi* interface offers interaction with the data in the form of pseudotime lineplot showing the evolution of each marker versus pseudotime (Figure 4C), where in the stage following the apoptotic region, we observe a gradual gain of CD4 (but not CD8). Thus, the biological explanation for this counterintuitive observation is that this stage is actually described in the reverse direction. It corresponds to the negative selection of CD4SP thymocytes in the thymic medulla where cells lose CD4 en route to apoptotic thymocytes. Topologically, trajectories form a triangular shape with corners in the DP thymocytes, CD4SP thymocytes and apoptotic thymocytes. Since the pseudotime distance from DP to apoptotic thymocytes is shorter than the real negative selection (DP to CD4SP to apoptotic cells), the calculation of the pseudotime inference cannot properly interpret the biological direction of the CD4SP cells to apoptotic cells. This highlights the value of the data-driven expert interpretation approach of *tviblindi*, which can solve even such complex topologies.

One of the most important questions in exploratory data analysis is whether the chosen level of detail is sufficient to exploit the information present in the data, or whether resolution needs to be increased. The interactive design of *tviblindi* allowed us to change the level of resolution on the persistence diagram and to investigate the random walks heading to apoptosis in more detail. After increasing the resolution, we observed that leaf V random walks split into two clear trajectories (Va and Vb, Figure 4D). The Va trajectories corresponded to the aforementioned failure to pass the positive/negative selection checkpoint. The Vb trajectories diverted from the canonical trajectory even earlier (Figure 4E), before the DP stage, and headed directly toward apoptosis.

Contrary to Va where the cells expressed intracellular TCRβ prior to Annexin V, the Vb cells failed to express TCRβ before entering the apoptotic trajectory (Figure 4F). We interpret this as failure to pass the β selection checkpoint due to unsuccessful TCRβ chain rearrangement. The later appearance of TCR β, after expression of Annexin V (Figure 4F), is caused by mixing with cells entering apoptosis at later developmental stages.

Similarly, if we focus on CD8 T cells, the same checkpoints can be discovered and interrogated (Supplementary Figure 4).

In summary, we show that *tviblindi* organized individual cells expressing various levels of 34 markers into trajectories leading to the expected ends of αβ T-cell development and discovered the corresponding checkpoints. Biological interpretation was enabled by plotting the expression of key markers along the developmental pseudotime. We were able to describe the canonical development of conventional CD4 T cells and CD8 T cells from their progenitors in the thymus to effectors in peripheral blood. We also found trajectories leading to apoptosis, when thymocytes failed to pass TCRβ selection or positive selection or when they were negatively selected (See Supplementary Video 2).

#### 2.2.3. Non-conventional end interpretation

The last analyzed trajectory remaining to be interpreted was the one leading to endpoint #6 (Figure 3A, Figure 5A). The corresponding population is hereafter referred to as End#6. End#6 was found in the thymic region (compare Figure 2B and Figure 3A for precise location in the *vaevictis* plot) and contained CD4 SP cells as well as a smaller fraction of cells which fall within the DP gate (Supplementary Figure 5A). The pseudotime line plot of CD8 expression shows that the End#6 cells gain dim expression of CD8 at the very end of the trajectory (Supplementary Figure 5B, Figure 5B). The possibility that CD4^pos^CD8^dim^ cells from the End#6 population are doublets consisting of a developing DP cell and a more mature CD4 SP T cell, as previously suggested (*72*), was ruled out by comparing the DNA content of End#6 SP and DP populations with the DNA content of dividing CD34^pos^ precursor cells (Supplementary Figure 5C).

**Figure 5:**
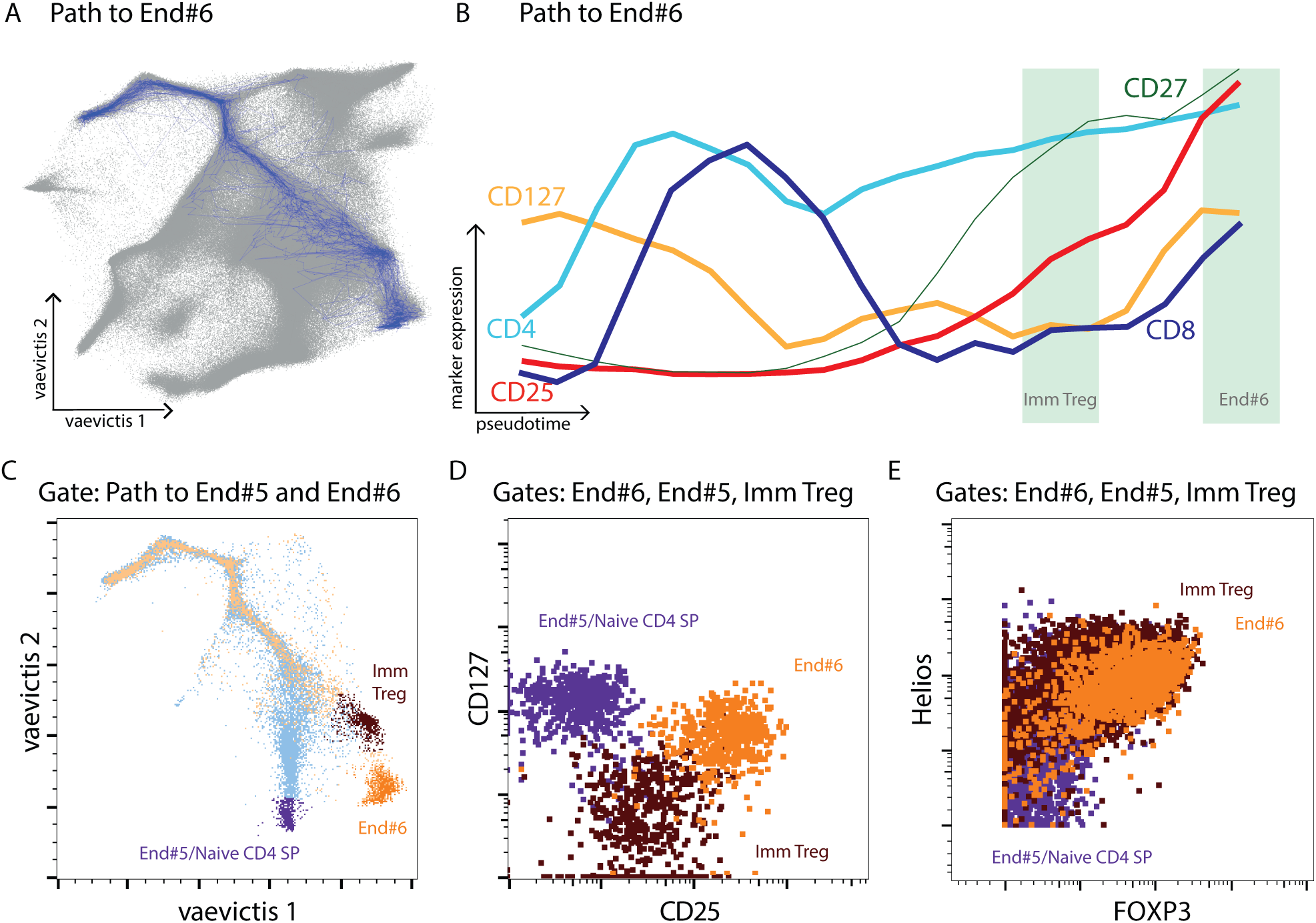
*tviblindi* analysis of the trajectory leading to End#6. (A) A trajectory leading to End#6 located in the thymic portion of the *vaevictis* plot (compare to Figure 2B). (B) Pseudotime line plot showing average expression of selected individual markers along developmental pseudotime. (C) *vaevictis* plot showing the trajectory to the conventional naive CD4 SP T cells (blue) and the trajectory to End#6 (orange). Conventional naive CD4 SP T cells are shown in purple, the immature Treg stage in brown and End#6 in orange. (D) Bivariate plot of CD25 and CD127 expression on gated conventional naive CD4 SP T cell (purple), immature Treg stage (brown) and End#6 (orange). (E) Bivariate plot of Helios and FOXP3 expression on conventional naive CD4 SP T cell (purple), immature Treg stage (brown) and End#6 (orange) in a validation experiment.

The trajectory leading to End#6 diverged from the conventional CD4 trajectory at the branching point of negative selection (Figure 5C, compare with Figure 4E) and continued through a distinct intermediate subset before reaching End#6. The End#6 cells have a CD25^hi^, CD127^pos^ phenotype, which distinguishes them from the preceding stage reminiscent of T regulatory (Treg) cells (CD25^pos^CD127^neg^), which could represent the immature Treg stage (Figure 5B and 5D). This observation led us to investigate the expression of Treg-specific markers in End#6 cells using a modified antibody panel (Supplementary Table Mass panel 2). After we gated the relevant populations and visualized them in the *vaevictis* plot (Supplementary Figure 6), we observed that FOXP3 (transcription factor of Treg cells) and Helios (transcription factor shown to be upregulated upon negative selection in mice (*21,73,74*)) is expressed by the CD25^pos^CD127^neg^ immature Treg cells as well as by the CD25^hi^CD127^pos^ End#6 cells (Figure 5E). Therefore, we hypothesized that End#6 cells constitute a specific Treg population (Figure 5D and 5E).

Some published reports describe the divergence of Treg cell development at the DP stage, with FOXP3 expression already detectable at the DP stage (*38,39,72,75–78*). Therefore, we wanted to ensure that positioning of the CD8^dim^ cells at the end of the trajectory is not an artifact of the TI. We decided to sort the populations of interest by fluorescence-activated cell sorting (FACS) and estimate their developmental order based on the analysis of the presence of T-cell receptor excision circles (TRECs).

We used a specific panel (Supplementary Table Mass panel 3) to select surface markers distinguishing between End#6 cells, immature Tregs, CD4 SP and DP cells (Supplementary Figure 7) and to design a gating strategy for sorting these populations (Supplementary Figure 8). To confirm that the sorting strategy led to identification of the correct populations, we overlaid the conventionally gated populations over the *vaevictis* plot (Supplementary Figure 9). Based on this analysis, we designed an 11-color panel for FACS sorting (Supplementary Table FACS panel) and sorted the key populations from thymus and from pediatric and adult peripheral blood (Supplementary Figure 10). We measured the presence of TRECs and T cell receptor alpha constant gene region (TCRAC) in the sorted populations to estimate the number of cell divisions undergone since the recombination of the TCRα chain (Figure 6A). The number of cell divisions, as estimated by the calculated TREC/TCRAC, ratio was lower than two in immature CD3^pos^ DP cells and immature Treg (DP as well as CD4 SP) cells. Thymic conventional mature naive CD4 SP did not feature a significant number of divisions. The TREC/TCRAC ratio in conventional naive CD4 SP cells in pediatric peripheral blood corresponded to 1-2 cell divisions, while in the adult peripheral blood it corresponded to 3-4 cell divisions. For CD45RA^neg^ Treg cells sorted from pediatric as well as adult peripheral blood, this was more than 5 cell divisions. For the End#6 CD4^pos^CD8^dim^ and End#6 CD4 SP cells, the TREC/TCRAC ratio corresponded to more than 5 divisions. These results confirm that CD4^pos^CD8^dim^ cells at the End#6 of the trajectory leading through immature Treg CD4 SP cells are not the true thymic DP cells. In agreement with our TI algorithm, we confirmed that End#6 cells represent a more developmentally advanced stage.

**Figure 6:**
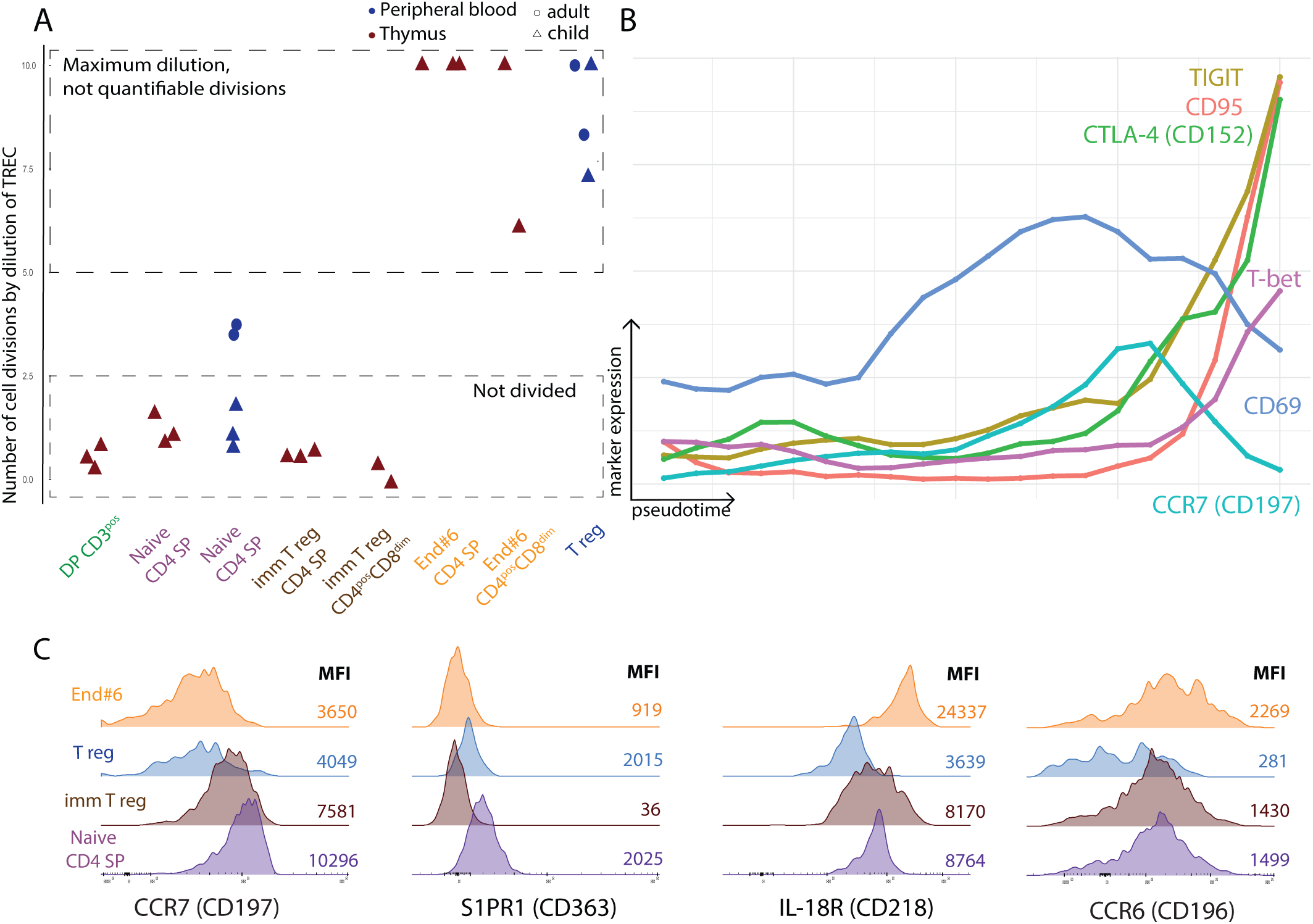
Detailed analysis of End#6. (A) Visualization of the TREC/TCRAC ratio, depicted as the calculated number of divisions for each population. Symbol code: cells from peripheral blood (blue), cells from thymus (brown), cells from adult donor (circle), cells from pediatric donor (triangle). (B) Pseudotime line plot depicting the expression of TIGIT, CD95, CD152, T-bet, CD69 and CD197 markers by cells from trajectory to End#6. (C) Overlaid histograms showing the expression of chemokine and cytokine receptors CD197, CD363, CD218 and CD196 in the respective populations. Symbol code: End#6 cells from thymus (orange), Tregs from peripheral blood (blue), immature Tregs from thymus (brown) and naive CD4 SP T cell from peripheral blood (purple).

Further investigation revealed an increase in the expression of several other markers at the end of the trajectory: apart from CD127 and CD8 the cells gained expression of T-bet, CD95, TIGIT and CD152 (CTLA-4) (Figure 6B). In fact, End#6 cells, irrespective of CD8 expression, phenotypically resembled the previously described long-lived thymic regulatory cells, which have been reported to recirculate from the periphery back to the thymus (*44,79–83*).

To test the hypothesis that End#6 cells are recirculating Tregs homing into thymus, we measured the expression of key chemokine receptors (*80,81,84–86*) in the populations of interest (Figure 6C). In contrast with the thymic mature naive CD4 SP (*84*), End#6 cells as well as immature Treg cells lack the S1PR1 (CD363) necessary to egress from the thymus. On the other hand, End#6 cells express IL-18R (CD218), which has been reported to cause upregulation of CCR6 (CD196) upon stimulation with IL-18 (*81*). CCR6 can then mediate the entry of Treg cells into the thymus (*81,85*). End#6 cells expressed more CCR6 compared to CD4 SP naive T cells and immature Treg cells. The last measured chemokine receptor, CCR7 (CD197) has been previously reported to distinguish the newly formed Treg cells, which do express CCR7, from the recirculating Treg cells which do not express CCR7(*80*). Consistent with this, while our immature Treg cells express CCR7, the End#6 cells do not (Figure 6C).

Since the thymic Treg population identified in the study of Park et al. (*45*), interpreted by the authors as long-residing in the thymus (here referred to as Treg-atlas), was placed in a terminal developmental position similar to End#6 (Figure 7A), we reanalyzed their single-cell RNA-seq data using *tviblindi* (See Supplementary note, section 9, Analysis of human thymus single-cell RNA-seq data). See Figure 7B for the localization of Treg-atlas on the *vaevictis* plot and Figure 7C for the trajectory leading to Treg-atlas. The RNA expression profile of Treg-atlas corresponds to the protein profile of End#6 suggesting that we are analyzing the same population (Figure 7D-F, compare to Figure 5B and Figure 6B, C). Apart from markers already presented in our mass cytometry panels, Treg-atlas show a sharp increase in *IL-1R2* RNA (Figure 7E) further reinforcing the hypothesis that these cells are recirculating from the periphery (*83,87–91*). Furthermore, Figure 7G shows the expression of additional chemokine receptors known to play a role in guiding Treg cells to peripheral sites (tumors and other sites of inflammation). Thus, *tviblindi* detected the trajectory leading to developmentally advanced Treg cells in single-cell RNA-seq data as well as in mass cytometry protein data.

**Figure 7:**
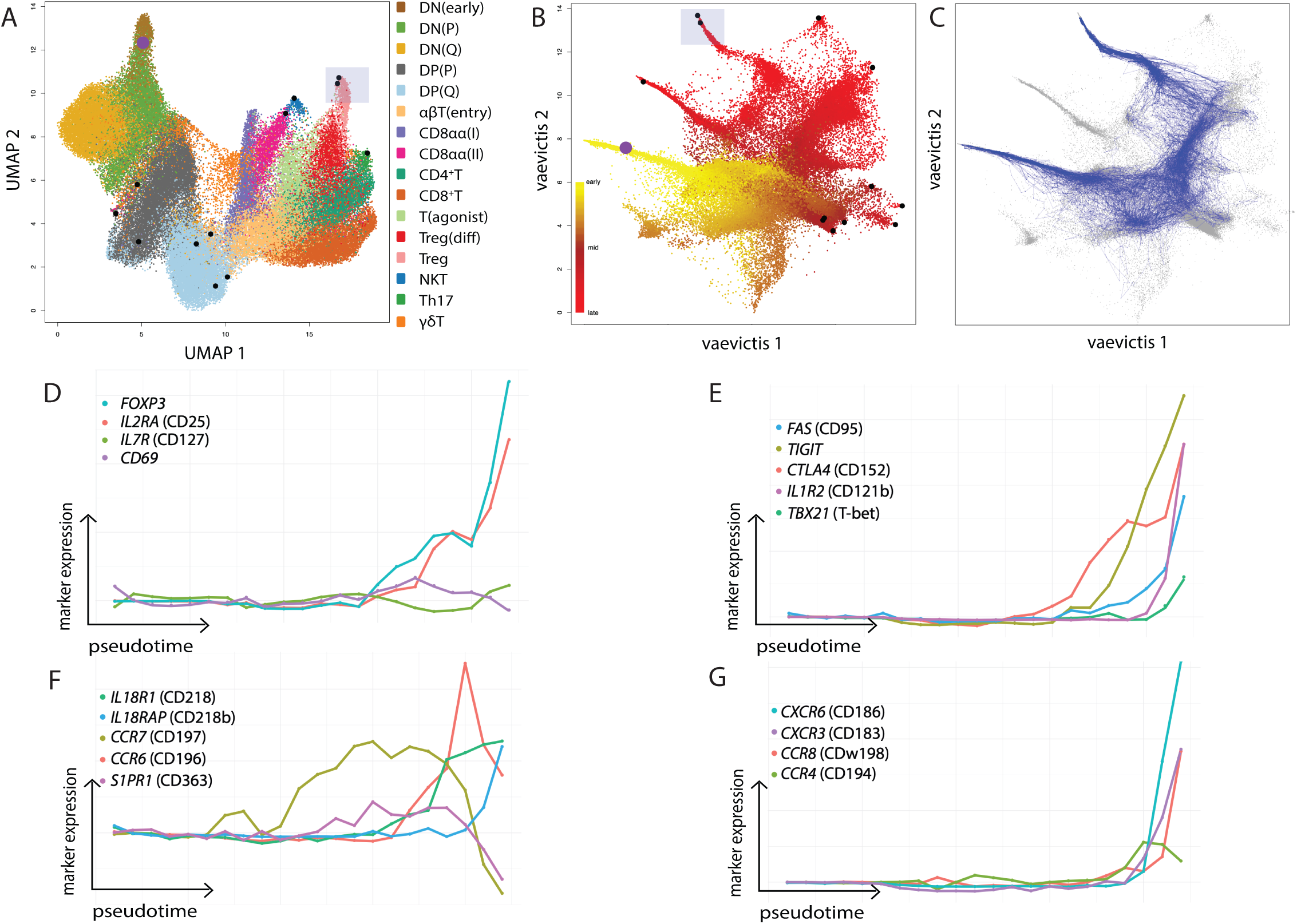
*tviblindi* analysis of single-cell RNA data from the study of Park et al. (45) showing the trajectory to Treg population corresponding to End#6. For A and B, the origin (centroid of DN (early) population) is marked by a purple dot, the candidate endpoints with more than 1% of simulated random walks are indicated by black dots and the endpoints corresponding to the Treg-atlas are highlighted by a blue rectangle. (A) UMAP plot identical to the one shown in Figure 3A of published research by Park et al. (45) showing the populations using the annotations from the authors, legend at the bottom left. (B) *vaevictis* plot of the same data as in A with shown pseudotime represented by yellow-to-red color gradient. (C) *vaevictis* plot showing the trajectory leading to Treg-atlas. For (D-G), the individual markers are labeled in accordance with the original research by their gene names and the CD designated markers are shown in parentheses. (D) Pseudotime line plot of markers canonical to the Treg population. (E) Pseudotime line plot of markers associated with Treg activation overlapping with our mass cytometry data from End#6 cells shown in Figure 6B. (F) Pseudotime line plot of cytokine and chemokine receptors overlapping with our cytometry data from End#6 cells shown in Figure 6C. (G) Pseudotime line plot showing the expression of additional chemokine markers measured on Treg-atlas.

## 3. Discussion

We have developed *tviblindi*, a modular and interactive TI tool which allows a biologist working with real-world high-dimensional datasets to explore the development of cells from progenitors toward differentiated stages. *tviblindi* simulates possible random walks from a user-defined point of origin and highlights their endpoints in the dataset. When used in an automated mode, it creates a “connectome” graph that visually represents the dataset with all ends and the trajectories connecting them to the point of origin. While most current TI tools stop here, we prefer to follow up with expert interaction with the data, which allows us to focus in depth on selected discovered ends and trajectories. *tviblindi* allows selection and grouping of similar random walks into trajectories based on visual interaction with the data. For each trajectory, marker intensity changes can be shown as the trajectory pseudotime progresses. Also, the key points (e.g., branching) can be projected back onto the two-dimensional visual representation of the dataset. *tviblindi* does not anticipate a single endpoint nor a single branching point with two endpoints. Importantly, it can be used for both mass cytometry (millions of cells but only dozens of parameters) as well as single-cell RNA-seq (thousands of parameters) data.

By integrating several modules (random walks simulation, pseudotime inference, real-time interactive topological classification using persistent homology and two-dimensional visualization using the *vaevictis* algorithm), *tviblindi* presents a feasible algorithmic solution for TI, which significantly extends upon existing TI algorithms. Its versatility and ability to interact with the user are its main contributions, creating a unique tool for a biologist to intuitively understand the dataset, visualize the data, and to focus the analysis on meaningful TI endpoints. An efficient discovery of the underlying dynamics is made possible by using persistent homology to aggregate random walks into trajectories, project them immediately onto a low-dimensional embedding (created using a custom dimensionality reduction algorithm: *vaevictis*) and visualize marker-vs-pseudotime line plots.

Importantly, the complete workflow has linear time complexity (See Supplementary note, subsection 8.3 Running times) and thus is manageable on a desktop computer in a reasonable computation time. We extensively compared its performance (See Supplementary Note, Sections 8 and 10 for details) to current TI algorithms and we found it to be superior in most criteria, as well as versatile in dealing with different datasets with widely varying numbers of cells and parameters investigated (single-cell RNA-seq or mass cytometry). It has been applied to publicly available datasets with success as well as to new data on B-cell development in health and immunodeficiency(*92*).

The dimensionality reduction is only used to visualize the data, and is not involved in any computation within the other modules. *vaevictis* reduction is designed to show gradual changes of single-cell phenotypes, suppressing the effect of abundant populations and showing local as well as global relationships of cells. This is in contrast to UMAP (*52,53*) and t-SNE (*54,55*), which prioritize clustering of locally similar cells at the expense of global relationships.

The *vaevictis* plot of mass cytometry data from thymus and peripheral blood faithfully depicted the basic scheme of T-cell development in this study and was also used to depict gradually changing data of T-ALL signaling (*93*)and B-cell development (*92*)(*94*).

When CD34^pos^ progenitors were selected as the population of origin in our dataset (thymocytes and PBMC), the endpoints of trajectories were found within peripheral CD4 and CD8 cells, the apoptotic cells, and the population of CD4^pos^CD8^dim^ thymic cells which we further explored in this paper and which we refer to as End#6 cells.

The endpoints at the thymus/peripheral blood transition (#5 for CD4 SP and #9 for CD8 SP) reflect the accumulation of cells at the stage of mature naive T cells. From this stage activated T cell may progress to become memory T cells. Of note, certain biological endpoints might be skipped if the final population is too small (the endpoint cells are not sufficiently represented) or if their phenotyping markers are missing (e.g., Th1-type, Th2-type markers are not included in any of our mass cytometry panels).

Furthermore, in the absence of markers resolving similar but developmentally unrelated cells, a random walk can create a connection between these unrelated cells and skip from one trajectory to another. When these immunophenotypically undistinguishable cells themselves represent developmental fate of two distinct trajectories, the two trajectories may combine into a single trajectory, skipping the endpoint in the middle. This phenomenon has been observed in the apoptotic region where some random walks leading to cells dying due to the failure of DP cells to pass the positive/negative selection checkpoint continued past the apoptotic stage in a reverse direction towards SP cells undergoing negative selection. Here the algorithm correctly found the trajectories, but expert input (understanding that apoptosis is a one-way process) was essential for the interpretation of the direction.

While the *vaevictis* embedding plots the cells on a 2D plot in an intuitive manner, some inherent limitations of 2D embeddings to comprehensively capture complex topologies were manifested. An example is the failure to visualize positive/negative selection of CD8 SP T cells, which is clearly detected by *tviblindi.* Note that when focusing on the trajectories of cells undergoing positive selection (V. Trajectories in Figure 4) in more detail, we have identified two groups of trajectories, one of them corresponding to positive selection (Va. trajectory) and one of them corresponding to an earlier β selection (cells entered the apoptotic pathway before expressing TCR β chain, Vb. trajectory). This highlights the benefit of expert-guided adjustment of the level of detail.

Investigation of the canonical CD4 and CD8 T-cell development reassured us that we can reconstruct the known sequence of maturation steps extensively described by CDMaps (*95*), connecting the thymic and peripheral T-cell compartment. We were able to dismiss artifacts caused by cell doublets. Putting expression levels of markers on a pseudotime axis showed us the proper sequence and intensity of expression changes. Note that, for the trajectory leading to CD4 SP cells, both CD4 and CD8 reach an expression peak at the DP stage, both decrease their intensity afterwards and then CD4 steadily increases towards the CD4 SP stage. CD4 expression level in peripheral SP cells is twice as high as in thymic SP cells. Key points of marker intensity changes can be traced back to the *vaevictis* 2D plots and groups of trajectories can be selected for further investigation (expression levels and 2D plot location).

A good example of the power of the presented approach was found when interpreting thymic Treg populations. The trajectory leading towards thymic End#6 cells diverges from the conventional CD4 trajectory at the location of negative selection. It goes through a CD45RA^neg^ immature Treg stage, characterized by the expression of CD25 in the absence of CD127 (conventional Treg immunophenotype), and continues towards the End#6 cells highly expressing CD25 and also expressing CD127. Both the immature Treg cells and the End#6 cells express Treg markers FOXP3 and Helios, confirming their Treg identity. The End#6 cells gain an expression of T-bet, CTLA-4, TIGIT, while they lose CD69, CD38 and CCR7. Surprisingly, they also gain a dim expression of CD8.

It has been shown that the ThPOK expression can be reduced in CD4 T cells, leading to reacquisition of CD8, and thus mimicking the DP immunophenotype. This was shown for CD8αα CD4 intraepithelial lymphocytes and CD8αβ CD4 cells in human blood as well as DP cells found in various tissues and under various disease settings. These have been reported to have enhanced cytotoxic functions or suppressive regulatory functions (*96,97*).

Although some of End#6 cells can fall within the conventional DP gate, the CD8 expression level is lower than that of the “bona fide” DP thymocytes and *tviblindi* positioned these CD4^pos^CD8^dim^ thymocytes at the end of the developmental trajectory. We wanted to make sure that their terminal end position is correct and that they are truly developmentally more advanced than the conventional immature DP cells and the immature CD4 SP Treg cells. Indeed, the proliferation history of End#6 cells (both CD8^neg^ and CD8^dim^) showed many more cell divisions than any other thymic subset, more than naive T cells in the periphery (by a measured dilution of TREC DNA in the sorted subsets). Our End#6 CD8^dim^ population may have been included in the FOXP3^pos^ DP cells placed at the beginning of Treg development in previous studies (*39,78*). Our approach provides a tool which can overcome the limitations of conventional gating and help to distinguish these cells.

However, we cannot claim that End#6 cells follow an uninterrupted developmental path from the precursors to End#6. In fact, the basic assumption of any TI method is that all developmental stages are present in the analyzed sample. If part of the development occurs in a peripheral organ which is not analyzed then an artificial shortcut can be mistaken for the whole path.

Indeed, the surface immunophenotype of End#6 cells agrees with the previously described population of peripheral Treg cells recirculating back to the thymus (*42,82*). These cells were reported to have the ability to regulate the development of their precursors either negatively, by competing for the limiting amount of IL-2 (*82*) or positively by blocking potentially harmful effects of the inflammatory cytokine IL-1 via their IL-1R2 decoy receptor (*83*). Furthermore, End#6 cells correspond to the Cluster 3 Tregs from the previously described thymus (*44*), where scRNA and surface protein expression is suggestive of their recirculating mature effector Treg phenotype. This is in line with the low TREC DNA content in End#6 cells. Future experimental studies specifically designed to investigate this Treg population should address whether these cells result from recirculation or represent an additional path of intrathymic Treg effector maturation. The future studies could benefit from a *tviblindi* analysis of their immunophenotyping datasets.

In conclusion, we have built and validated a TI method that has generic use for mass cytometry, flow cytometry and single-cell RNA-seq. *tviblindi* is modular and interactive, thus combining the information strength of rich single-cell datasets with the power of expert knowledge for the interpretation of developmental trajectories.

## Supporting information

Supplementary Table Mass panel 1

## Acknowledgement

T.K. and J.S. were financially supported by project NU23-07-00170 of the Czech Republic Ministry of Health. Institutional support was provided by the project National Institute for Cancer Research (Project No. LX22NPO5102), funded by the European Union–Next Generation EU. D.N. is a fellow of FWO (Fonds Wetenschappelijk Onderzoek – Vlaanderen), supported by Strategic Basic research grant 1S40423N.

## 4. Methods

### 4.1. Human cells

Thymocytes were obtained from healthy donor thymi of pediatric patients undergoing heart surgery at the Motol University Hospital, after informed consent was given by the patient guardians, in accordance with the Declaration of Helsinki. Peripheral blood mononuclear cells (PBMC) were obtained by density gradient centrifugation from buffy coats of healthy adult donors, at Institute of Hematology and Blood Transfusion in Prague. The pediatric PBMC were obtained from healthy pediatric patients undergoing pre-surgical examination for non-malignant and otherwise unrelated medical issue after consent was given by patients’ guardians. Cells were stored in liquid nitrogen and thawed when needed for the experiment. After thawing, the cells were quickly transferred into 10 ml of pre-warmed RPMI supplemented with 10% FBS and 100µg of Pulmozyme (dornase alpha, Roche, Basel, Switzerland) for 30 min at 37°C. In order to remove dead cells and avoid clumping of the thymocytes, the cells were centrifuged at low speed 108g for 15 minutes. Subsequent steps were performed at 4°C.

### 4.2. Mass cytometry

#### 4.2.1. Reagents

Unless otherwise stated, buffers and isotope labeled monoclonal antibodies (moAbs) were purchased from Standard BioTools Inc. (South San Francisco, USA). In-house antibody conjugations were performed using MaxPar labeling kits (Standard BioTools Inc.) according to the manufacturer’s instructions. Carrier-protein-free moAbs were purchased from Exbio (Prague, Czech Republic), BioLegend (San Diego, USA), R&D Systems (Minneapolis, USA), eBioscience (San Diego, USA, part of Thermo Fisher Scientific Waltham, USA), Miltenyi Biotec (Bergisch Gladbach, Germany), Cell Signalling Technology (Danvers, USA) (for details see Supplementary Table Mass panels).

##### Antibody cocktails

The amount of labeled moAbs needed for one test was determined by titration. Two sets of antibody cocktails were prepared and frozen as described previously (*98*): 1) the antibody cocktail containing moAbs against surface antigens, which was applied prior to fixation and permeabilization steps, and 2) antibody cocktail containing moAbs against intracellular antigens, which was applied after the permeabilization.

#### 4.2.2. Staining

Cells were washed in Annexin binding buffer, ABB (Exbio), and counted, approximately 5 million thymocytes and 1 million PBMC were used for one experiment.

To minimize technical variability, the individual tissues were barcoded with CD45 Abs conjugated with different isotopes (30 min, 4 °C). Biotin labeled Annexin V was added together with the barcoding Abs, to distinguish apoptotic cells with exposed phosphatidylserine residues. After this incubation, the sample volume was risen to 1 ml and cisplatin 198Pt was added to distinguish dead cells (5 min, 4 °C). Cells were then washed twice in ABB (92 g, 15 min, 4 °C) and the samples were pooled together, spun (92 g, 15 min, 4 °C) and processed further as one sample.

The surface Ab cocktail, including the isotope labeled anti-biotin mAb to visualize the annexin, was added to the cell pellet (total volume of surface Abs was approx. 100 µl, 30 min, 4 °C).

Cells were washed twice in ABB (92 g, 15 min, 4 °C), prefixed with 1 ml freshly diluted 1,6% PFA (10 min, RT) and washed once more in Maxpar® Cell Staining Buffer, CSB (1050 g, 5 min, 4 °C). Then, the cells were fixed and permeabilized using Maxpar® Nuclear Antigen Staining buffer (30 min, 4 °C) followed by two washings in Maxpar® Nuclear Antigen Staining Perm (1050 g, 5 min, 4 °C).

In cases when the FOXP3 antibody was used, the cells stained with the surface Abs and washed in ABB were fixed and permeabilized in FOXP3 fixation/permeabilization buffer (30 min, 4 °C) followed by two washings in FOXP3 permeabilization buffer (600 g, 5 min, 4 °C).

The intracellular Ab cocktail was added to the cell pellet, together with anti-TCRb frame antibody labeled with APC (it was not possible for us to get this antibody in the unlabeled carrier-free form allowing in-house isotope conjugation) (the total volume of intracellular Abs was approx. 60 µl, 30 min, 4°C). The cells were washed twice in CSB (1050 g, 5 min, 4 °C). Finally, an APC directed isotope labeled mAb was added (30 min, 4 °C), cells were washed twice in CSB (1050 g, 5 min, 4 °C), fixed in 1 ml freshly diluted 1.6% formaldehyde (10 min, RT) and centrifuged (1050 g, 5 min, 4 °C).

#### 4.2.3. Mass cytometry samples acquisition

After incubation in 125nM 191/193Ir in Maxpar® Fix and Perm Buffer (at least 12 hours, up to 2 weeks), the cells were washed twice in CSB (1050 g, 5 min, RT) and once in Maxpar® Cell Acquisition Solution (CAS, 1050 g, 5 min, RT). Then they were diluted up to 1×10^6^ cells/ml in 15 % EQ^TM^ Four Element Calibration Beads (Standard BioTools Inc.) in CAS and filtered through a 35 μm nylon mesh cell-strainer cap (BD Biosciences, San Jose, CA). The samples were acquired using Helios (Standard BioTools Inc.), CyTOF software version 6.7.1014. The instrument was tuned for acquisition using Tuning Solution (Standard BioTools Inc.) according to the manufacturer’s instructions. The noise reduction (the cell length 7–150, lower convolution threshold 200) was applied during the acquisition. The signal was normalized using a Standard BioTools Inc. algorithm, which is based on the ‘Bead Passport’ concept.

#### 4.2.4. Analysis of mass cytometry data

After export to listmode data (FCS format), files were further processed using FlowJo (v10, BD Biosciences). First, nucleated cells were gated by metal-tagged DNA intercalator 191/193Ir, and thymus versus peripheral blood was debarcoded by manual gating (Supplementary Figure 1). Next, PBMC were gated on CD3^pos^ lineage negative and TCR γδ negative T cells. The thymus cells were gated on lineage negative, viable (platinum negative) and TCR γδ negative thymocytes. Thus, only TCR αβ T cells and their progenitors in the thymus were used for further processing by the *tviblindi* algorithm described later. When manual gating was performed prior to *tviblindi*, the FlowJo workspace file was used to save manual gate positions, so that the gated cells could be displayed in the GUI, to help the interpret the *vaevictis* plot.

The *tviblindi* algorithm was programmed in R, C++ and Python. The source code is deposited on GitHub (https://github.com/stuchly/tviblindi).

The listmode data was transformed using *arcsinh* with cofactor 5 for subsequent analysis (*99*). After data interrogation by *tviblindi*, an ‘enhanced’ FCS file was created, containing artificial channels with newly computed variables (2-dimensional embedding coordinates, pseudotime, pathway membership identifier, labels and selected points on trajectories) and a graphical output for figures was created using FlowJo layouts.

### 4.3. FACS sorting

Human thymocytes and PBMC obtained as described above for mass cytometry experiments were stained with appropriate amount of fluorescently labeled monoclonal antibodies, at 1 million cells/50 µl for 30 minutes on ice in the dark (see Supplementary Table FACS panel). After incubation, cells were washed in ice-cold PBS and sorted using BD FACS Aria III cell sorter into tubes with 10 µl of ice-cold Platinum Direct cell lysis buffer with Proteinase K (ThermoFisher Scientific). The obtained subsets (see Supplementary Figure 10) were then directly processed according to the manufacturer’s instructions to minimize any loss of material.

### 4.4. TREC analysis

Extracted DNA was used for the quantification of T-cell receptor excision circles (TREC) and T cell receptor alpha constant gene region (TCRAC) via previously established real-time PCR assay (*100,101*). Estimated number of cell divisions was calculated based on the ratio between TREC and TCRAC copies in these populations.

All the background information on *tviblindi* design including used versions of R packages, dataset used for testing, dimensionality reduction, pseudotime estimation, topological clustering and biological interpretation of re-used datasets is available in the Supplementary note. Please, see the supplementary note also for detailed description and guide through the user interface and comparison with other state-of-the-art-methods (*102–136*).

## Notes

https://github.com/stuchly/tviblindi

### Competing Interest Statement

The authors have declared no competing interest.

### Summary of Updates

This version expands the evaluation of the framework on complex scRNA-seq datasets, the discussion of hyperparameters, and adds additional TI methods for comparison.

