## Supplementary Table Mass panel 1 for "Deconstructing Complexity: A Computational Topology Approach to Trajectory Inference in the Human Thymus with *tviblindi*"

### Supplementary Table CyTOF/FACS panel

| HUMAN | | | | | Mass panel 1 | Mass panel 2 | Mass panel 3 | FACS panel | | | | Spectral panel | | | |
|---|---|---|---|---|---|---|---|---|---|---|---|---|---|---|---|
| CD marker | clone | metal | cat# (MaxPar)/LOT (home-made) | Manufacturer | | | | CD marker | clone | fluorochrome | Manufacturer | CD marker | clone | fluorochrome | Manufacturer |
| BARCODES and markers applied with the barcode, before pooling the samples | | | | | | | | | | | | | | | |
| CD45 | HI30 | Y89 | 3089003B | Fluidigm | x | x | x | | | | | | | | |
| CD45 | MEM-28 | 115In | 181010 | Exbio | x | x | x | | | | | | | | |
| Annexin V biotin | | | 640904 | BioLegend | x | x | x | | | | | | | | |
| cisPt | | 198Pt | | Fluidigm | x | x | x | | | | | | | | |
| SURFACE | | | | | | | | | | | | | | | |
| CD123 | 6H6 | 106Cd | 200310 | BioLegend | | x | x | | | | | | | | |
| DR | L243 | 111Cd | 200310 | BioLegend | | | x | | | | | | | | |
| CD95(Fas) | DX2 | 113Cd | 200310 | BioLegend | | x | x | CD95 | LT95 | PE-Cy7 | Exbio | CD95 | DX2 | BUV737 | BD |
| TCRab | BW242/412 | 141Pr | 190208 | Miltenyi Biotec | | x | x | | | | | | | | |
| CD28 | CD28.2 | 142Nd | 190809 | BioLegend | x | x | x | | | | | | | | |
| Anti-Biotin | 1D4-C5 | 143Nd | 3143008B | Fluidigm | | x | | | | | | | | | |
| CD69 | FN50 | 144Nd | 3144018B | Fluidigm | x | x | x | | | | | | | | |
| CD4 | RPA-T4 | 145Nd | 3145001B | Fluidigm | x | x | x | CD4 | OKT4 | BV510 | Biolegend | CD4 | OKT4 | BV510 | Biolegend |
| CD3 | UCHT1 | 146Nd | 200925 | Exbio | x | x | x | CD3 | UCHT1 | PerCP_Cy5.5 | Exbio | CD3 | UCHT1 | PerCP_Cy5.5 | Exbio |
| CD8 | MEM-31 | 148Nd | 181129 | Exbio | x | x | x | CD8 | SK1 | APC-H7 | BD | CD8 | SK1 | APC-H7 | BD |
| CD34 | 581 | 149Sm | 3149013B | Fluidigm | x | x | x | | | | | | | | |
| CD5 | L17F12 | 150Nd | 190919 | Exbio | x | | | | | | | | | | |
| CD2 | TS1/8 | 151Eu | 3151003B | Fluidigm | x | x | x | | | | | | | | |
| TCRgd | 11F2 | 152Sm | 3152008B | Fluidigm | | x | | | | | | | | | |
| HLA-DR | L243 | 152Sm | 181108 | BioLegend | x | | | | | | | | | | |
| CD279(PD-1) | EH12.2H7 | 153Eu | 190729 | Exbio | | x | | | | | | | | | |
| TIGIT | MBSA43 | 154Sm | 3154016B | Fluidigm | | x | x | | | | | | | | |
| CD279(PD-1) | EH12.2H7 | 155Gd | 3155009B | Fluidigm | | | x | | | | | | | | |
| CD56 | B159 | 155Gd | 3155008B | Fluidigm | | x | | | | | | | | | |
| CD71 | MEM-75 | 156Gd | 200925 | Exbio | x | x | x | | | | | | | | |
| CD1a | SK9 | 158Gd | 180227 | Exbio | x | x | x | CD1a | HI49 | PE | Exbio | CD1a | HI149 | BUV395 | BD |
| CD197/CCR7 | G043H7 | 159Tb | 3159003A | Fluidigm | | | x | | | | | | | | |
| TCR_Va7.2 | 3C10 | 160Gd | 191018 | BioLegend | x | | | | | | | | | | |
| CD38 | HIT2 | 160Gd | 181221 | Exbio | | | x | CD38 | HIT2 | A700 | Exbio | CD38 | HIT2 | A700 | Exbio |
| CD27 | O323 | 161Dy | 161111 | BioLegend | x | | | | | | | | | | |
| CD152(CTLA-4) | 14D3 | 161Dy | 3161004B | Fluidigm | | x | x | | | | | | | | |
| CD25 | MEM-181 | 162Dy | 190621 | Exbio | x | x | x | CD25 | MEM-181 | APC | Exbio | CD25 | MEM-181 | APC | Exbio |
| CD45RA | MEM-56 | 163Dy | 180905 | Exbio | x | x | x | CD45RA | MEM-56 | PE-DyLight-594 | Exbio | CD45RA | L48 | PEDazzle594 | Exbio |
| CD127 | eBioRDR5 | 165Ho | 190919 | eBioscience | x | x | x | CD127 | HIL-7R-M21 | FITC | BD | CD127 | HIL-7R-M21 | FITC | BD |
| CD44 | BJ18 | 166Er | 3166001B | Fluidigm | x | x | x | | | | | | | | |
| CD27 | L128 | 167Er | 3167006B | Fluidigm | | x | x | CD27 | M-T271 | BV605 | BD | CD27 | M-T271 | BV605 | BD |
| CD19 | HIB19 | 169Tm | 3169011B | Fluidigm | x | x | x | CD19 | HIB19 | BV421 | Biolegend | CD19 | HIB19 | BV421 | Biolegend |
| CD56 | NCAM16.2 | 169Tm | 190322 | BD Biosiences | x | | x | CD56 | NCAM16.2 | BV421 | BD | CD56 | NCAM16.2 | BV421 | BD |
| CD13 | WM15 | 169Tm | 200227 | Exbio | x | x | x | CD13 | WM15 | BV421 | Biolegend | CD13 | WM15 | BV421 | Biolegend |
| CD33 | WM53 | 169Tm | 3169010B | Fluidigm | x | x | x | CD33 | WM53 | BV421 | Biolegend | CD33 | WM53 | BV421 | Biolegend |
| CD16 | 3G8 | 169Tm | 161111 | Exbio | x | | x | CD16 | 3G8 | BV421 | Biolegend | CD16 | 3G8 | BV421 | Biolegend |
| Biotin | 1D4-C5 | 170Er | 3170003B | Fluidigm | x | | x | | | | | | | | |
| NKT | 6B11 | 170Er | 3170015B | Fluidigm | | x | | | | | | | | | |
| CD38 | HIT2 | 172Yb | 3172007B | Fluidigm | | x | | | | | | | | | |
| CXC3CR1 | 2A9-1 | 172Yb | 3172017B | Fluidigm | | | x | | | | | | | | |
| TCRgd | B1 | 173Yb | 191118 | BioLegend | x | | x | | | | | | | | |
| CD161 | HP-3G10 | 173Yb | 180321 | BioLegend | | x | | | | | | | | | |
| HLA-I | W6/32 | 195Pt | 170912 | Exbio | x | | | | | | | | | | |
| HLA-I | W6/32 | 196Pt | 170912 | Exbio | | x | x | | | | | | | | |
| CD47 | CC2C6 | 209Bi | 3209004B | Fluidigm | x | | x | | | | | | | | |
| CD16 | 3G8 | 209Bi | 3209002B | Fluidigm | | x | | | | | | | | | |
| | | | | | | | | | | | | CCR7 (CD197) | G043H7 | BV785 | Biolegend |
| | | | | | | | | | | | | S1P1 | SA258H3 | PE | Biolegend |
| | | | | | | | | | | | | IL18R (CD218a) | H44 | PE | Biolegend |
| | | | | | | | | | | | | CCR6 (CD196) | 11A9 | PE/Cy7 | BD |
| | | | | | | | | | | | | DAPI | | | |
| INTRACELLULAR | | | | | | | | | | | | | | | |
| Ki67 | Ki-67 | 141Pr | 190910 | BioLegend | x | | | | | | | | | | |
| cPARP | F21-852 | 143Nd | 3143011A | Fluidigm | x | | x | | | | | | | | |
| P-Tyr | P-Tyr-01 | 147Sm | 190809 | Exbio | x | x | x | | | | | | | | |
| pRb[Ser807/811] | J112-906 | 150Nd | 3150013A | Fluidigm | | x | x | | | | | | | | |
| TCRVbeta3.1 | JOVI.1 | 152Sm | 200109 | BioTechne | | | x | | | | | | | | |
| CyclinB1 | GNS-1 | 153Eu | 3153009A | Fluidigm | x | | | | | | | | | | |
| BCL-2 | Bcl-2/100 | 153Eu | 190118 | Exbio | | | x | | | | | | | | |
| Bim | C34C5 | 154Sm | 181129 | Cell Signaling | x | | | | | | | | | | |
| pSLP-76[Y128] | J141-668.36.58 | 155Gd | 190919 | BD Biosiences | x | | | | | | | | | | |
| pS6[S240/244] | D68F8 | 159Tb | 190118 | Cell Signaling Tecl | x | | | | | | | | | | |
| FoxP3 | 259D/C7 | 159Tb | 3159028A | Fluidigm | | x | | | | | | | | | |
| Helios | 22F6 | 160Gd | 190322 | BioLegend | | x | | | | | | | | | |
| Tdt | E17-1519 | 164Dy | 3164015B | Fluidigm | x | x | x | | | | | | | | |
| Gata3 | TWAJ | 167Er | 3167007A | Fluidigm | x | | | | | | | | | | |
| Ki-67 | B56 | 168Er | 3168007B | Fluidigm | | x | x | | | | | | | | |
| T-bet | REA102 | 171Yb | 180906 | Miltenyi Biotec | x | x | x | | | | | | | | |
| pS6 [S235/S236] | N7-548 | 172Yb | 3172008A | Fluidigm | | | | | | | | | | | |
| RUNX3 | 527327 | 174Yb | 161220 | R&D Systems | x | | | | | | | | | | |
| BLNK | 2B11 | 174Yb | 200109 | BD Biosiences | | x | x | | | | | | | | |
| Perforin | B-D48 | 175Lu | 3175004B | Fluidigm | x | | | | | | | | | | |
| pHistoneH3[Ser28 | HTA2 | 175Lu | 3175012A | Fluidigm | | x | x | | | | | | | | |
| TCR betaF1 APC | 8A3 | | 566052 | BD Biosiences | x | | x | | | | | | | | |
| SECONDARY INTRACELLULAR | | | | | | | | | | | | | | | |
| Anti-APC | APC003 | 176Yb | 3176007B | Fluidigm | x | | x | | | | | | | | |
