## Supplementary figures for "Deconstructing Complexity: A Computational Topology Approach to Trajectory Inference in the Human Thymus with *tviblindi*"

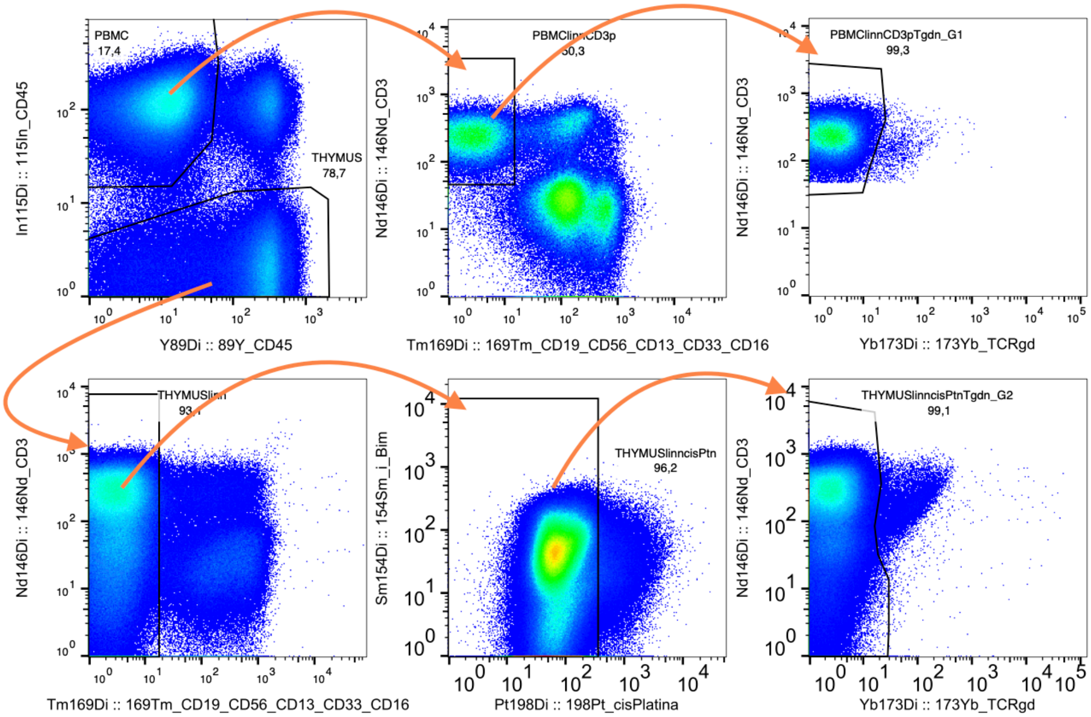


**Supplementary Figure 1: Gating strategy for CyTOF data from thymus stained with Mass panel 1** After gating the nucleated cells as positive for metal-tagged DNA intercalator 191/193Ir (not shown), the thymus versus PBMC were debarcoded by gating. Next, PBMC were gated to select CD3 positive cells negative for lineage markers and TCR $\gamma\delta$. Thymocytes were gated to select live cells (platinum negative), negative for lineage markers and TCR $\gamma\delta$. After gating, the cells from thymus and PBMC were merged and used for visualization and for trajectory inference by *tviblindi*.

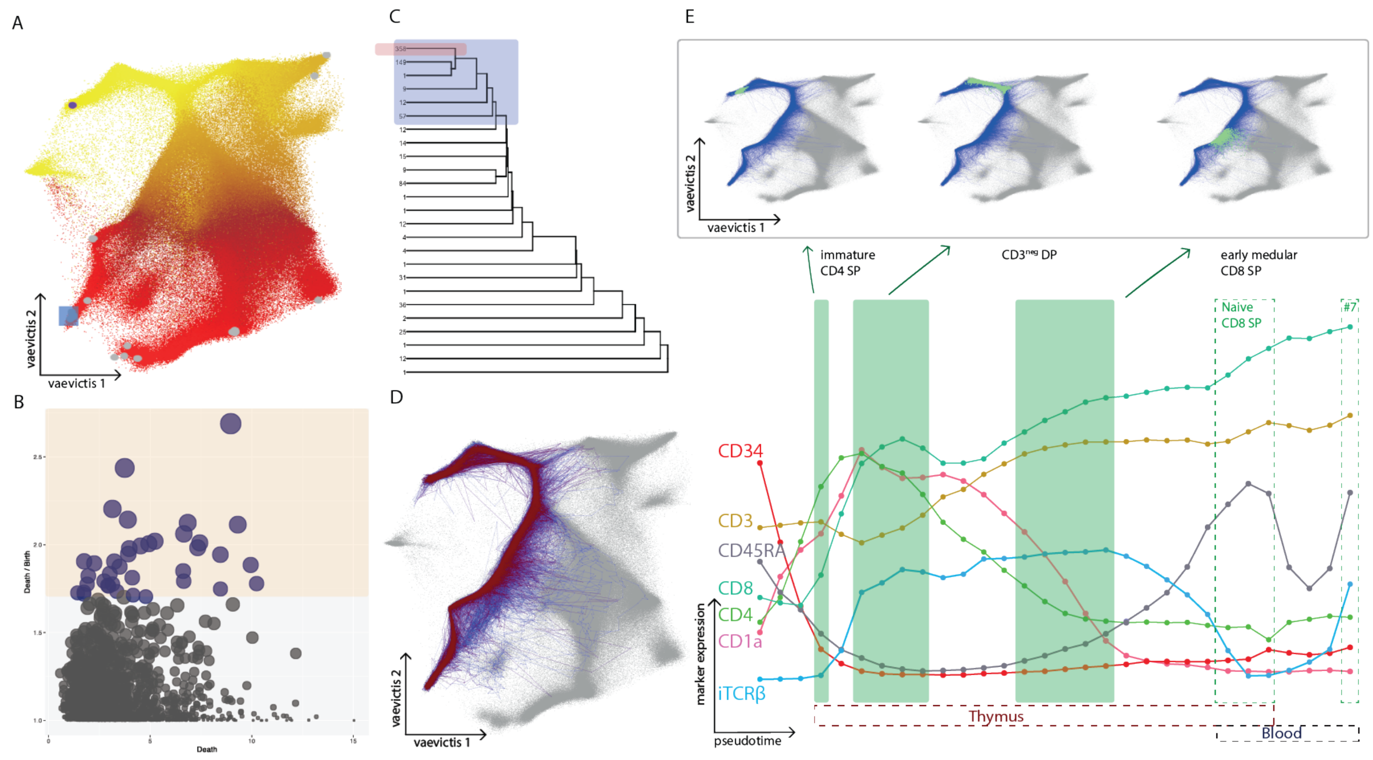


**Supplementary Figure 2: *tviblindi* analysis of trajectories leading to CD8** (A) *vaevictis* plot of the T-cell developmental space (thymus and T cells from peripheral blood) where the estimated pseudotime is shown as yellow-to-red color gradient with $CD34^{pos}$ progenitors as the population of origin (purple dot). Discovered endpoints of development are shown (gray dots). Selected endpoint representing the $CD8^{pos}$ effector memory T cell (#7) is highlighted (gray rectangle). (B) Persistent homology graph where sparse regions (holes) in the point cloud of measured cells are selected (orange rectangle). (C) Dendrogram of clustered trajectories shows a larger group of random walks (blue rectangle) and a subcluster of 358 random walks (red rectangle). (D) *vaevictis* plot of the above selected trajectories shows their topology. (E) Pseudotime line plot showing the average expression of selected markers along the developmental pseudotime (bottom), selected areas of interest (green rectangles) and their position shown (green dots in insert) on the *vaevictis* plot of T-cell development.

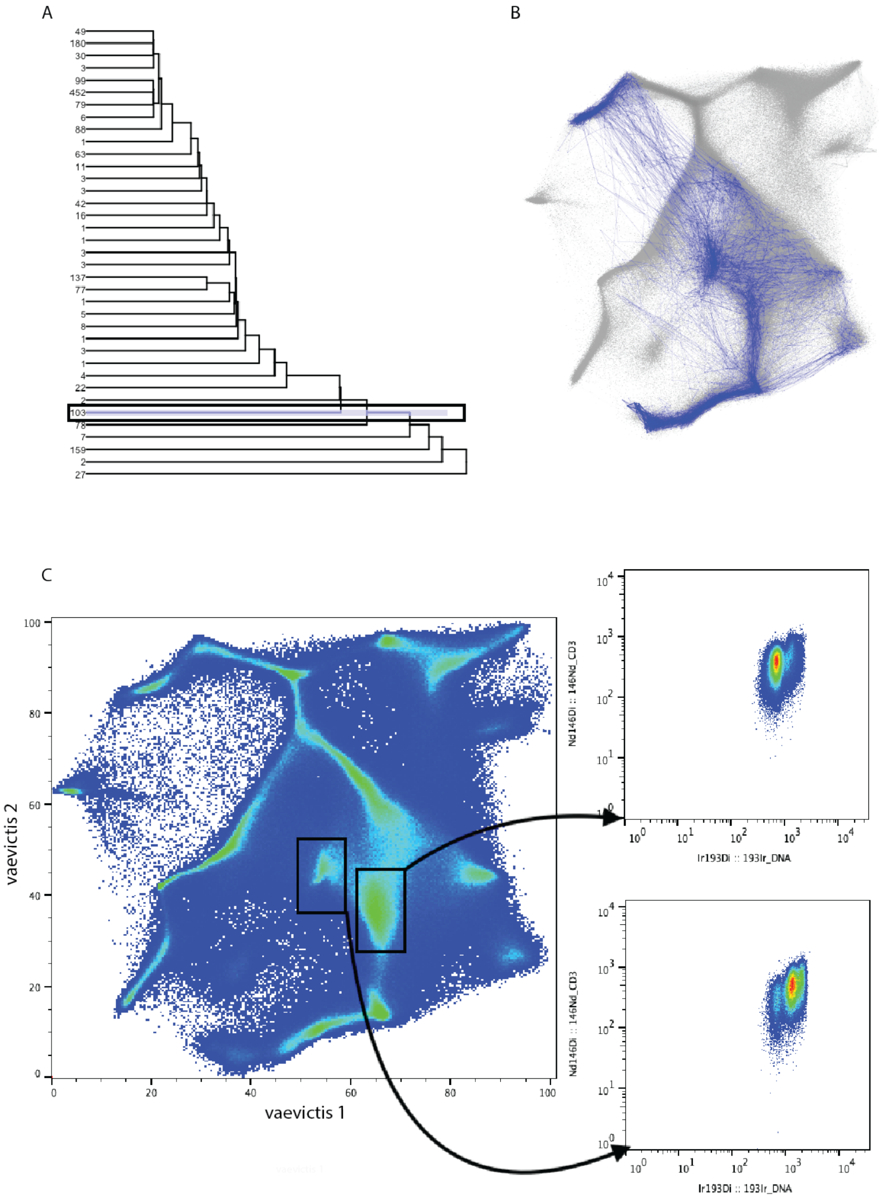


**Supplementary Figure 3: Artifact caused by analysis of doublets** (A) Dendrogram representing the clustered random walks leading to endpoint #1. A leaf containing 103 random walks was selected for further analysis (blue rectangle). (B) *vaevictis* plot of a trajectory showing a strange connection between immature CD4 SP and the „hub“ population. (C) Hub population has double DNA content (bottom dot plot) compared to mature CD4 SP T cells (top dot plot). The doublets are difficult to exclude from datasets containing high numbers of dividing cells (such as thymocytes).

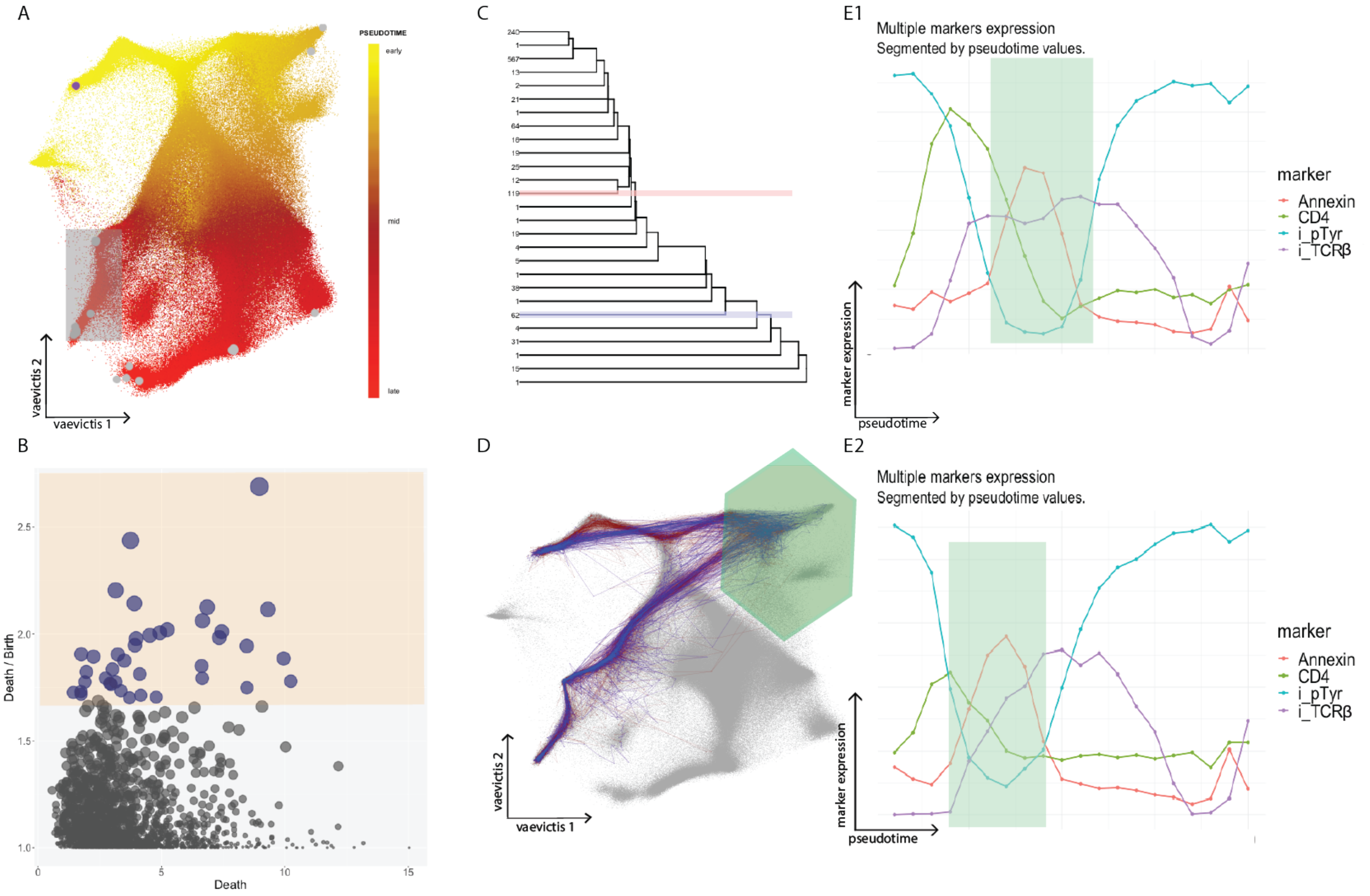


**Supplementary Figure 4: Developmental checkpoints on trajectories to CD8 SP T cells** (A) The same *vaevictis* plot of T-cell development in thymus and peripheral blood as seen in Figure 4A. Estimated pseudotime is represented by a yellow-to-red color gradient with CD34$^{pos}$ progenitors as the population of origin (purple dot). Gray dots indicate the discovered developmental endpoints. The gray rectangle highlights the endpoints selected for further investigation. All CD8 SP endpoints in peripheral blood (naive and memory CD8 SP) were selected to increase the number of interrogated random walks. (B) Persistent homology diagram where sparse regions in the point cloud of measured cells were selected (orange rectangle). Apart from the canonical trajectories (Supplementary Figure 5), two abundant leaves can be selected Va (blue rectangle) and Vb (red rectangle). (D) *vaevictis* plot of the selected trajectories in respective colors shows their topology. (E) Pseudotime line plots, which depict the average expression of selected markers along the trajectories Va (E1) and Vb (E2). The region of apoptosis is highlighted in green. The first part of the trajectory Va (until the apoptotic region) represents the positive/negative selection of DP cells in thymic cortex. The first part of the trajectory Vb represents $\beta$ selection. The latter part of the trajectories (after the apoptotic region) represents the negative selection of CD8 SP cells in reverse direction (same situation as in the CD4 compartment described in the main text and Figure 5).

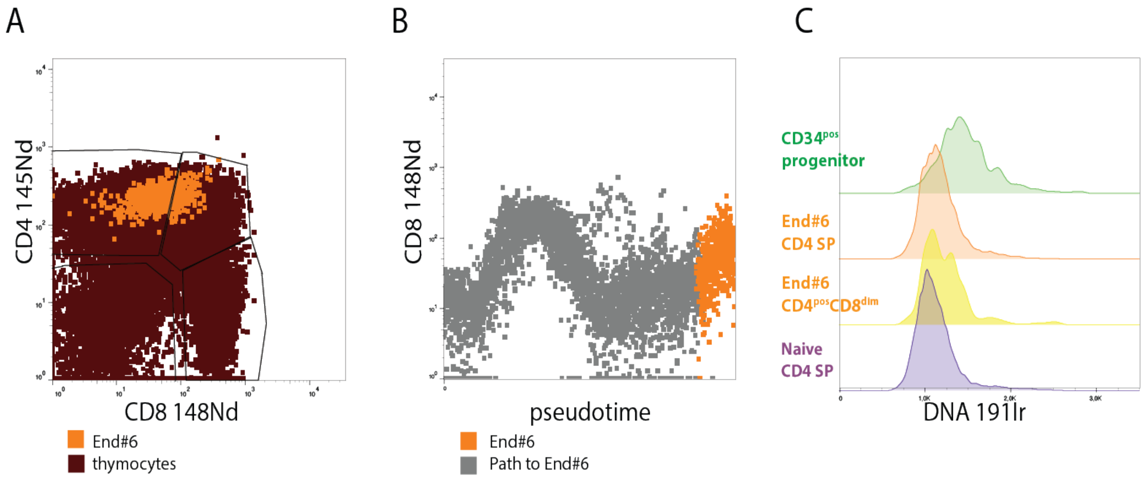


**Supplementary Figure 5: Initial characterization of End#6 cells** (A) Overlay of End#6 subset (highlighted in orange) dot plot over the conventional gating of thymocytes (brown background). (B) Thymocytes belonging to the trajectory to End#6, showing a late upregulation of CD8 on the pseudotime vs. CD8 dot plot. (C) Histograms of the DNA content in dividing CD34$^{pos}$ progenitors (green), End#6 CD4 SP (orange) and End#6 CD4$^{pos}$ CD8$^{dim}$ (yellow) cells compared to naive CD4 SP thymocytes (purple). Note that while events with more than one copy of DNA are present in CD34$^{pos}$ subset, other subsets do not contain high DNA content.

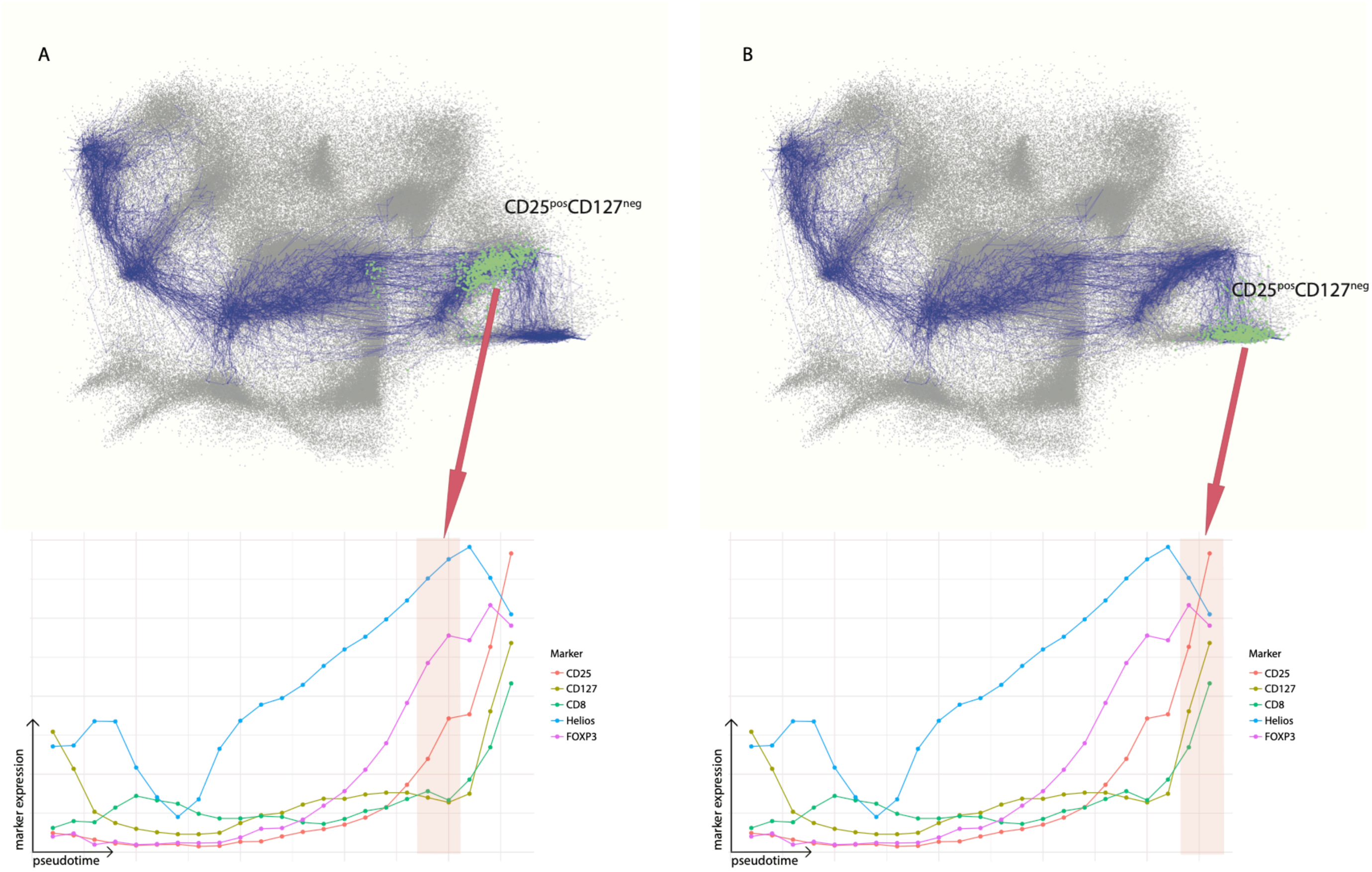


**Supplementary Figure 6: *tviblindi* analysis of End#6 cells stained with Mass panel 2** The pseudotime line plots showing the dynamics of the development of CD8, Helios and FOXP3 markers in the trajectory leading to the population corresponding to End#6 in the modified panel (Mass panel 2). As in the original panel, we discovered two clearly defined populations CD25$^{pos}$ CD127$^{neg}$ (A bottom) and CD25$^{pos}$ CD127$^{pos}$ (B bottom), which can be identified based on the expression dynamics of CD25 and CD127 markers. The populations (highlighted in red rectangles) along with random walks are shown on the *vaevictis* plot above. The cell populations highlighted in green (top panels) correspond to the cells selected by the red rectangles.

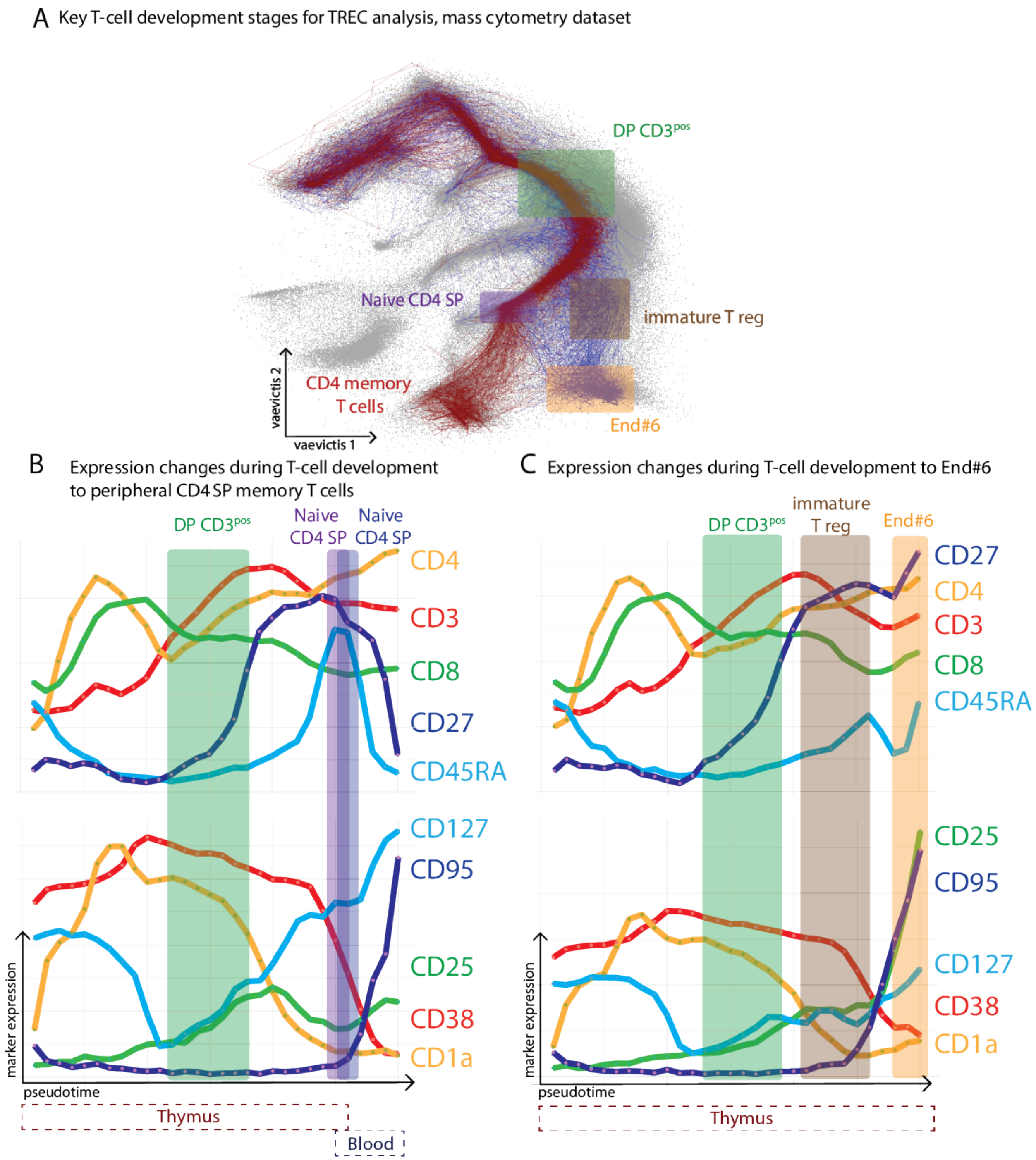


**Supplementary Figure 7: *tviblindi* analysis of CyTOF data from thymus stained with Mass panel 3** (A) *vaevictis* plot of thymocytes and PBMC stained with Mass panel 3 (see Supplementary Table). Key T-cell developmental stages selected for further analysis and investigation are highlighted: CD3$^{pos}$ DP cells (green), naive CD4 SP (purple), immature Treg (brown), End#6 cells (orange) and CD4 memory T cells (red). (B) Pseudotime line plot depicting the expression of developmental markers on the trajectory leading to conventional CD4 SP cells in thymus and peripheral blood as labeled below the plot. DP CD3$^{pos}$ T cells are positive for CD38, CD1a and negative for CD27, CD45RA. Naive CD4 SP have low expression of CD38 and are negative for CD1a and positive for CD27 and CD45RA. (C) Pseudotime line plot depicting the expression of developmental markers on the trajectory leading via immature Tregs (brown) to End#6 cells (orange). Immature Treg cells are positive for CD38, negative for CD1a and CD45RA, positive for CD25 and negative for CD127. The End#6 cells are negative for CD38, CD45RA, positive for CD127, positive for CD95 and highly positive for CD25.

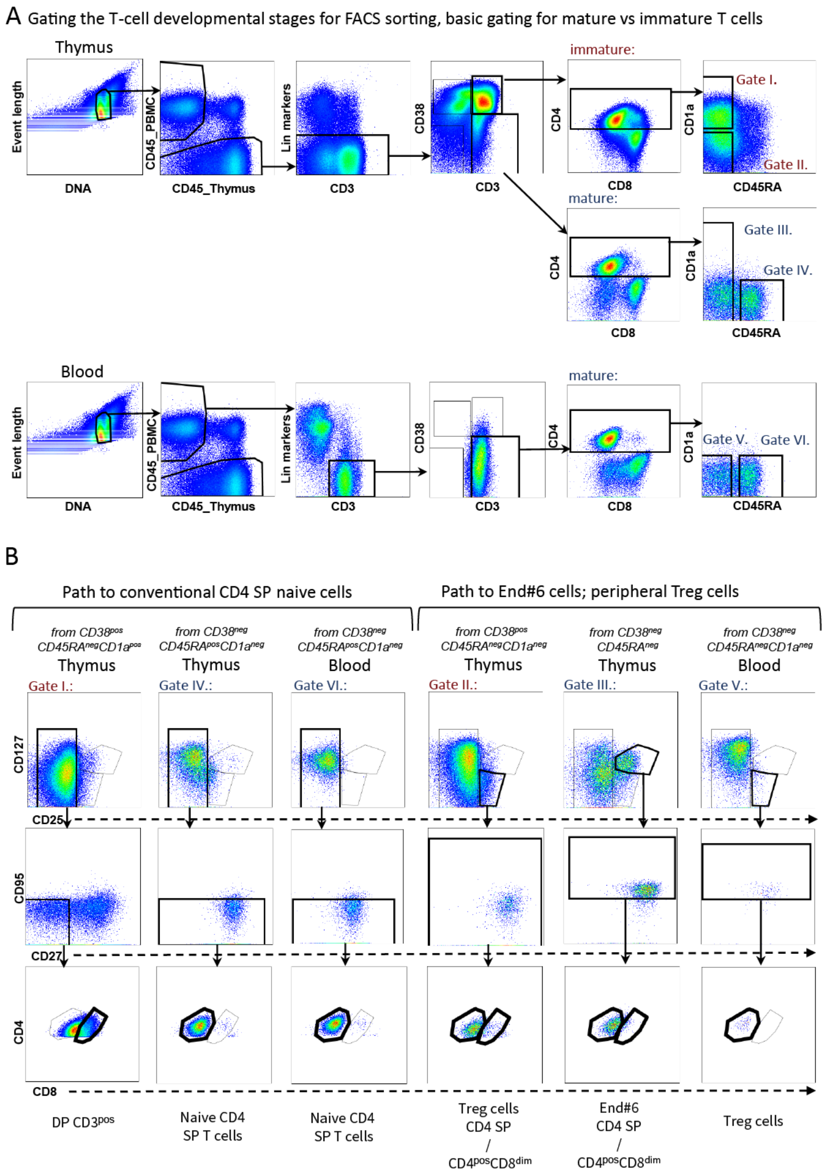


**Supplementary Figure 8: Conventional gating of populations of interest stained with Mass panel 3** (A) Gating strategy of samples from thymus and peripheral blood stained with Mass panel 3 (see Supplementary Table) which was used to design a sorting panel. The surface markers, which distinguish the populations of interest (labeled as Gate I. to Gate VI.) were used, based on their expression shown in Supplementary Figure 7. Gates containing immature CD38$^{pos}$ cells are labeled in red font, gates containing mature CD38$^{neg}$ cells are labeled in blue font. (B) Each population (Gate I. to VI.) as defined in part A is gated further based on the CD25xCD127 expression, maturity (CD95xCD27) and CD4xCD8 expression. The populations, which were finally designated for sorting are indicated by bold frame of their respective gates.

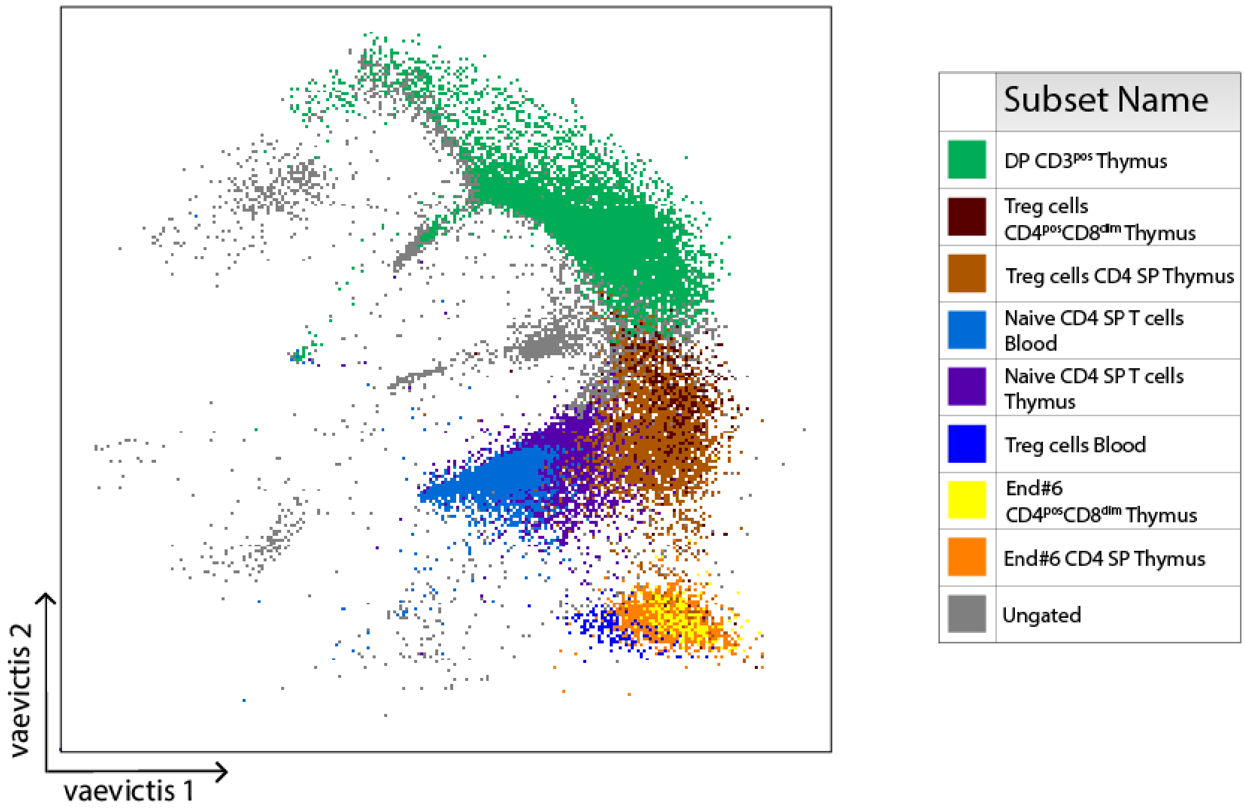


**Supplementary Figure 9: Overlay of conventionally gated populations on *vaevictis* plot** *vaevictis* plot of the thymus and PBMC with an overlay of manual gates as described in Supplementary Figure 8 validating the appropriate position of the key cell populations. Each color indicates one cell population, color code to the right.

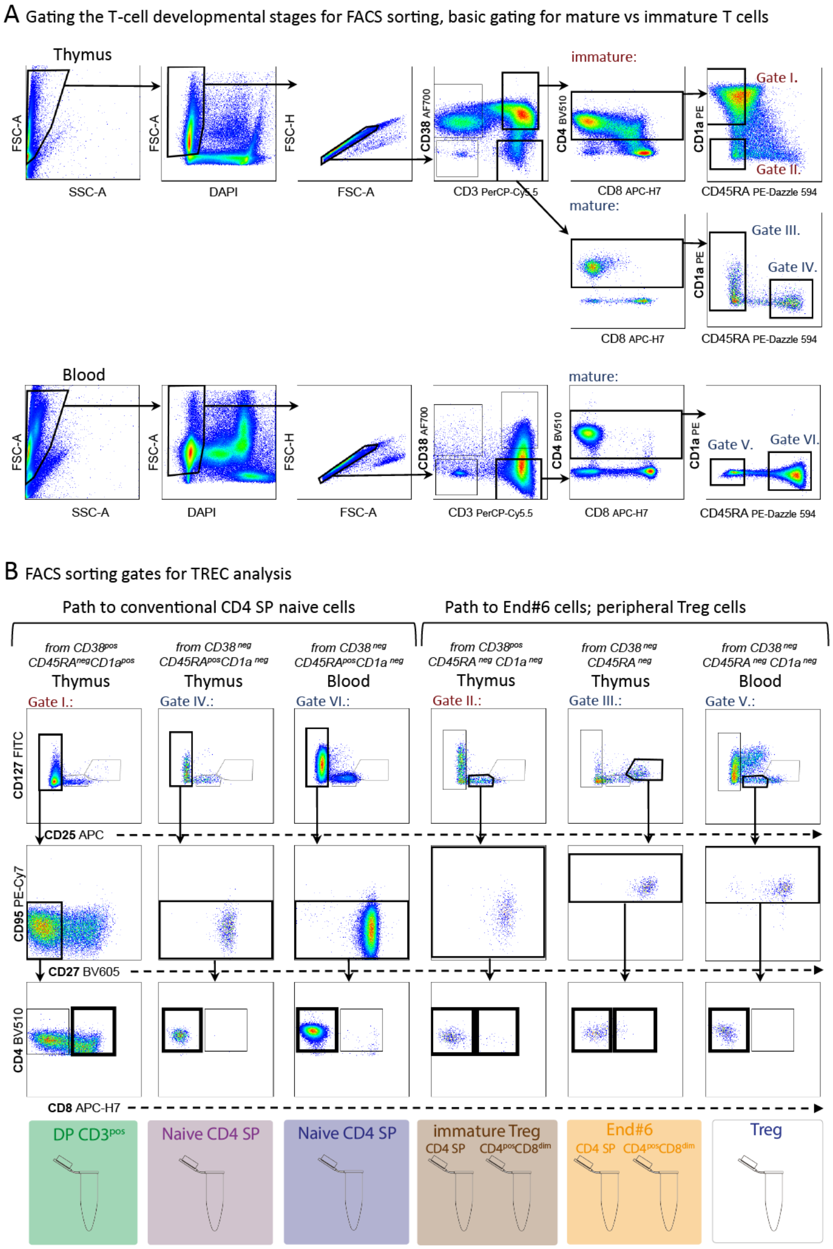


**Supplementary Figure 10: Gating strategy for sorting** (A) Common gating scheme of thymocytes and PBMC on FACS Aria sorter (refer to Supplementary Table). (B) Gating strategy of the key stages, which mimicked the CyTOF Mass panel 3 gating shown above in Supplementary Figure 8. Eight T-cell populations were sorted: DP $CD3^{pos}$ (green), naive CD4 SP from thymus (purple), naive CD4 SP from peripheral blood (blue), immature Tregs CD4 SP and $CD4^{pos}$ $CD8^{dim}$ (brown), End#6 cells CD4 SP and $CD4^{pos}$ $CD8^{dim}$ from thymus (orange) and Tregs from peripheral blood (white).
