## Supplementary note for "Deconstructing Complexity: A Computational Topology Approach to Trajectory Inference in the Human Thymus with *tviblindi*"

**5 Connectome** **28**

**6 Graphical user interface (GUI)** **28**

**7 Running the analysis** **35**

**8 Performance evaluation** **37**

**11 Software acknowledgement 80**

**References 82**

### 1 Datasets

In this section we give a brief description of datasets used in this supplementary note.

#### 1.1 *dyntoy* dataset

For the purposes of basic demonstrations we created an artificial dataset using the *dyntoy* R package (Cannoodt and Saelens (2022)) as follows:

```
devtools::install_github('dynverse/dyntoy')
library(dyntoy)
n_events <- 10000
n_features <- 3896 #randomly chosen value from typical range
set.seed(12345)
d <- dyntoy::generate_dataset(
  id           = 'tviblindi_dyntoy_test',
  model        = 'connected',
  num_features = n_features,
  num_cells    = n_events
)
```

Package *dyntoy* simulates single-cell expression data organized into trajectories. It creates a random milestone network (Figure 1a). The *milestones* (named *M#*) can be understood as populations of cells. We used the first 25 principal components for downstream analysis. To further highlight some issues that come up in trajectory inference, we modified this dataset by up-sampling a population labeled *M10*, by iterating over all points assigned to this population and adding random convex combinations of their 30 nearest neighbours. This resulted in a new dataset with $35,795$ points where $26,532$ points lie in the population *M10*; we will refer to this dataset as the 'up-sampled' *dyntoy* dataset (see Figure 1d,e).

##### 1.2 Human thymus & PBMC

A mass cytometry dataset from human thymus and peripheral mononuclear cells from healthy donors (PBMC) was measured using *Mass panel 1.* (Supplementary Table, Mass panel 1). The analyzed dataset consists of 1, 182, 802 events and 34 dimensions (markers).

##### 1.3 Human thymic development scRNA-seq

A single-cell RNA-seq dataset presented in Park et al. (2020) was used. Data were downloaded from `https://zenodo.org/record/5500511`. The `seurat_object` was created from the raw counts (`HTA07.A01.v02.entire_data_raw_count.h5ad`) restricted to the subset of T cells. Annotations of cells were retrieved from `HTA08.v01.A06.Science_human_tcells.h5ad`. Data were preprocessed by *Seurat* (Hao et al. (2021)) and *Rmagic* (van Dijk et al. (2018)). The preprocessed dataset consists of 76, 994 events and 2000 dimensions.

```
require(Seurat)
s.genes <- cc.genes$s.genes
g2m.genes <- cc.genes$g2m.genes
seurat_object <- NormalizeData(seurat_object)
seurat_object <- FindVariableFeatures(seurat_object)
seurat_object <- CellCycleScoring(seurat_object,
                  s.features = s.genes, g2m.features = g2m.genes, set.ident = TRUE)
seurat_object <- ScaleData(seurat_object,
                  vars.to.regress = c("sample","method","S.Score","G2M.Score"))
seurat_object <- RunPCA(seurat_object, npcs = 50, verbose = FALSE)
```

```
require(Rmagic)
data<-magic(t(seurat_object@assays$RNA@scale.data))$result
```

##### 1.4 Bone marrow atlas

A CITE-seq atlas of human bone marrow published by Zhang et al. (2024) was used. The h5ad object containing RNA counts was downloaded from https://www.synapse.org/Synapse:syn53222756 and loaded into R with the schard package (https://github.com/cellgeni/schard); ADT counts were downloaded from https://www.synapse.org/Synapse:syn53222702. The dataset contains 71,460 cells from 4 donors, which we integrated with Seurat's RPCA method with the following commands:

```
options(future.globals.maxSize = 10e+09)
library(Seurat)

# convert h5ad to seurat
library(schard)
BM <- schard::h5ad2seurat("adata_combined_rna_adt_annotated-titrated.h5ad")

# Add ADT counts
adt <- fread("ADTs-Titrated-TotalVI-log2-transposed.zip")
adtm <- as.matrix(adt[,2:ncol(adt)])
rownames(adtm) <- adt$uid
adtm <- adtm[colnames(BM),]
BM[['ADT']] <- CreateAssayObject(data=t(adtm))

# Remove ADT counts from the RNA assay slot
```

```
BM <- subset(BM, features = setdiff(rownames(BM), tail(rownames(BM), 142)))

# Split the RNA assay into layers by sample
BM[["RNA"]] <- split(BM[["RNA"]], f = BM[["sample"]][,1])

# Normalize data and find variable features
BM <- NormalizeData(BM, normalization.method = "LogNormalize")
BM <- FindVariableFeatures(BM)
BM <- ScaleData(BM)
BM <- RunPCA(BM)

# Integrate layers
BM_integrated <- IntegrateLayers(
  object = BM, method = RPCAIntegration,
  orig.reduction = "pca", new.reduction = "integrated.dr",
  normalization.method = "LogNormalize",
  verbose = FALSE)
BM_integrated <- RunUMAP(BM_integrated,reduction = "integrated.dr",dims = 1:50)

#Create a list to be used in section 10
BMlist <- list(
        integrated_pca = Embeddings(BM_integrated, "integrated.dr"),
        meta = BM_integrated@meta.data)
```

##### 1.5 Atlas of mouse gastrulation

An RNA-seq atlas of mouse gastrulation from E6.5 to E8.5 published by Pijuan-Sala et al. (2019) was used. The dataset was downloaded from `https://content.cruk.cam.ac.uk/jmlab/atlas_data.tar.gz`. The atlas contains (after filtration) 116,312 cells from 36 samples which are well corrected. The dataset is loaded into Seurat with the corrected principal components provided by the authors with the following code:

```
# Load data
metadata <-read.table("meta.tab",
           header = TRUE, sep = "\t", stringsAsFactors = FALSE)
corrected_pcas <- readRDS("corrected_pcas.rds")

# Select overlapping cells
overlap <- intersect(metadata$cell, rownames(corrected_pcas$all))

metadata <- metadata[match(overlap, table = metadata$cell),]
corrected_pcas <- corrected_pcas$all[overlap,]

# Create a list to be used in section 10
MGlist <- list(
        pca = corrected_pcas,
        meta = metadata[,c("sample", "stage", "celltype")],
        origin_name = "Epiblast",
        umap = metadata[,c("umapX", "umapY")])
```

#### 2 Dimensionality reduction

To meet the requirements for data visualization in trajectory inference (TI) problems, i.e. continuity, robustness against the differences in sizes of cell populations and straightforward extrapolation to new datasets, we implemented a variational autoencoder model based on ideas in Szubert et al. (2019); Ding et al. (2018), called *vaevictis*. The objective function is a weighted sum of reconstruction error, variational regularization (divergence from a latent prior distribution) and two regularization terms: the loss function value adapted from $t$-SNE and triplet loss from *ivis*. For additional speedup (which makes the time complexity virtually independent of dataset size) and improved robustness against differences in population size, the dataset is by default first clustered using $k$-means clustering and a fixed number of representatives are sampled from each cluster. This method is implemented in a separate GitHub repository: `https://github.com/stuchly/vaevictis`.

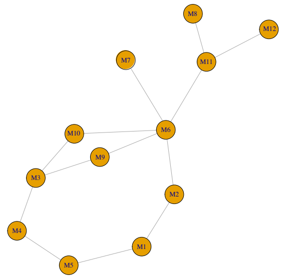

(a) Topology of *dyntoy* data

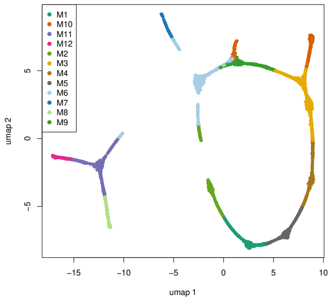

(b) 2D UMAP embedding of *dyntoy* data

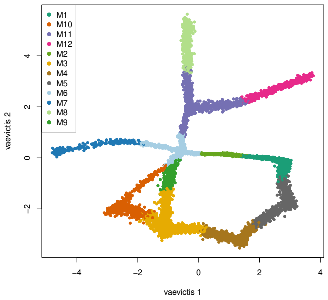

(c) 2D *vaevictis* embedding of *dyntoy* data

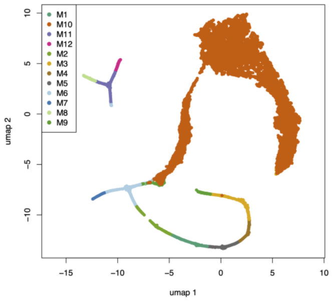

(d) 2D UMAP embedding of up-sampled *dyntoy* data

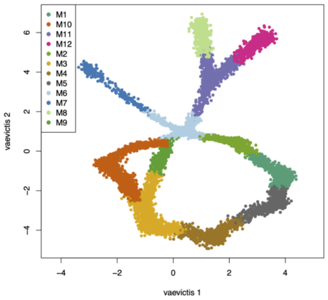

(e) 2D *vaevictis* embedding of up-sampled *dyntoy* data

Figure 1: *dyntoy* data structure preservation with dimensionality reduction

Figure 1 illustrates the performance of *vaevictis* in producing a 2-dimensional embedding of the original and the up-sampled *dyntoy* dataset(subsection 1.1), as compared to a corresponding UMAP embedding. While the overall structure of this data (panel $a$) is preserved by *vaevictis* (panel $c$), even for data with numerically unbalanced populations (panel $d$; the population *M10* was enlarged to contain $\sim 74\%$ of all cells, see subsection 1.1), UMAP preserves global structure comparatively worse (panel $b$ and $d$).

### 3 Pseudotime estimation & random walks simulation

#### 3.1 Theoretical concepts

Let $G = (V, E)$ be $k$ nearest neighbor graph ($k$-NNG) (or any suitable connected graph over the data), where $V$ and $E$ stand for vertices (cells) and edges of the graph respectively. We have the corresponding *distance matrix* $S$

$$S_{ij} = \begin{cases} d(v_i, v_j) & (v_i, v_j) \in E \\ \infty & \text{otherwise} \end{cases} \tag{1}$$

where $d$ is a distance function (we use Euclidean distance). For modeling diffusion on $G$ we also consider the *transition matrix* $T$

$$T_{ij} = \begin{cases} f(S_{ij}) & (v_i, v_j) \in E \\ 0 & \text{otherwise} \end{cases} \tag{2}$$

where $f$ is a suitable kernel function. Our default choice is the Gaussian kernel

$$f(x) = e^{-x^2/\varepsilon},$$

where $\varepsilon$ is determined from the distribution of vertices as follows: for an edge $(v_i, v_j) \in E$ $\varepsilon = \text{median}\{d(v_i, v_k)^2, (v_i, v_k) \in E\}$. For convenience we define the *transition probability matrix* $P$

$$P = D^{-1}T \tag{3}$$

where $D$ is a diagonal matrix of row-sums of $T$.

We will calculate the *pseudotime* of each vertex of $G$ in the terms of expected length of random walks until a vertex is reached.

**Definition 1** (Random walk). *We define a random walk (of possibly infinite length) on $G = (V, E)$ as a series $v^0, v^1, \ldots$ of vertices in $G$. Each $v^{i+1}$ is chosen from the set $\{v_j : \{v_l, v_j\} \in E, v_l = v^i\}$ with probability $p_{v_j} = P_{lj}$ ($P$ defined in Equation 3).*

If we fix the starting point $v_0$ of random walks we can define the *pseudotime* as one of the following quantities.

**Definition 2** (Pseudotime). *Let $v_0 \in V$ be fixed. We define pseudotime $\tau_{v_0}(v)$ for each $v \in V$ as either* **expected hitting time** *(the expected length of a random walk sequence starting in $v_0$ before it reaches $v$) or* **expected hitting distance** *(the expected distance traveled by a random walk starting in $v_0$ before it reaches $v$). For both definitions it holds that $\tau_{v_0}(v_0) = 0$.*

The *expected hitting time* is calculated (for $v_j \neq v_0$) as follows (e.g. in Förster et al.

(2022)):

$$\tau_{v_0}(v_j) = \sum_{v_i:\{v_i,v_j\}\in E} P_{ij}(\tau_{v_0}(v_i)+1) \text{ if } v_j \neq v_0,\ \tau_{v_0}(v_0)=0$$

The generalization to *expected hitting distance* is straightforward:

$$\tau_{v_0}(v_j) = \sum_{v_i:\{v_i,v_j\}\in E} P_{ij}(\tau_{v_0}(v_i)+S_{ij}) \text{ if } v_j \neq v_0,\ \tau_{v_0}(v_0)=0$$

Rewriting this in matrix form (with components corresponding to $v_0$ dropped), we get

$$\boldsymbol{\tau} = P\boldsymbol{\tau} + (P \odot S)\mathbf{1}$$

where $\odot$ denotes the Hadamard (element-wise) product and $\mathbf{1}$ is a vector of ones. We can solve for $\tau$ the equation

$$(I-P)\boldsymbol{\tau} = (P \odot S)\mathbf{1} \tag{4}$$

where $I$ denotes the identity matrix. Note that if $T$ is symmetric (which is by default enforced by setting $T_{ij} = max(T_{ij}, T_{ji})$) the Equation 4 can be rewritten as $D^{1/2}(I - P)D^{-1/2}D^{1/2}\tau = D^{1/2}(P \odot S)\mathbf{1}$, where $D^{1/2}(I-P)D^{-1/2}$ is a symmetric matrix. The Equation 4 is a sparse system which can be efficiently solved by iterative methods, such as the (bi)conjugate gradient method, even for graphs with millions of vertices. This allows to keep the resolution of pseudotime on the single-cell level[1].

Once pseudotime is estimated, we can use it to simulate finite random walks. We do it

[1]Note that we are solving a boundary problem with the boundary condition $\tau_{v_0}(v_0) = 0$. It is straightforward to generalize this condition to $\tau_{v_I}(v_I) = \phi(v_I)$ with an arbitrary function $\phi$ and arbitrary set of indices $I$. This way, we can prescribe the values of pseudotime at any vertex. As long as the adjacency matrix, with rows and columns corresponding to $I$ dropped, represents a connected graph, the problem is well posed and has a unique solution.

by creating a directed acyclic graph (DAG) $G' = (V, E')$, where $E' = \{(v_i, v_j) \in E,\ \tau(v_j) > \tau(v_i)\}$. Random walks on a DAG are necessarily finite, and their final vertices are suitable candidates of developmental endpoints. We simulate large number of walks on $G'$ and record the endpoints for further analysis (e.g. on Figure 2 in gray), where we study the possible trajectories (represented by sets of spatially coherent random walks) by which the endpoints can be reached.

The main motivation for introducing the *expected hitting distance* was our need for robust pseudotime estimation, even for the cases of significant accumulation of cells at some stages of development. The *expected hitting time* tends to rise quickly within compact, numerically large populations, and this would yield undesirable points of attraction when simulating developmental pathways in our downstream analysis. To illustrate this issue, we took the up-sampled *dyntoy* dataset (see subsection 1.1 and Figure 2) and set the point closest to the centroid of population *M4* as $v_0$. If pseudotime is calculated as *expected hitting time*, population *M10* forms a sink which 'traps' any random walk entering it (see the color gradient on Figure Figure 2$c$). In contrast, pseudotime based on *expected hitting distance* allows random walks to pass through population *M10* (Figure 2$d$). Upon inspection of Figure 2$b$, we see that passing through *M10* would be equivalent to going back in time when using *expected hitting time* formulation.

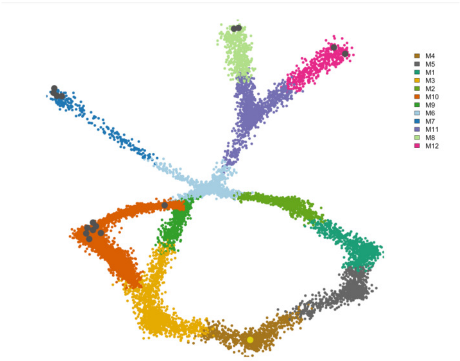

(a) *vaevictis* embedding of *dyntoy* data with marked populations

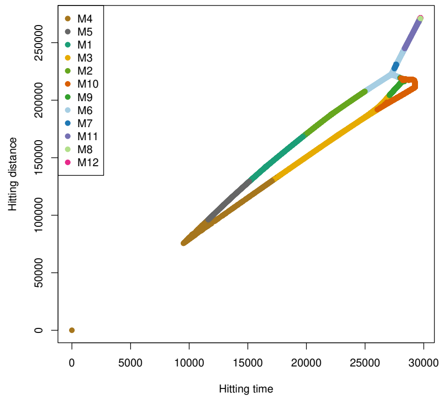

(b) Comparison of ‘expected hitting time‘ and ‘expected hitting distance‘ plotted for each point, colored by population

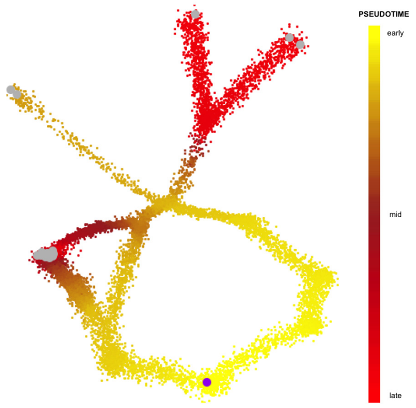

(c) Pseudotime: hitting time

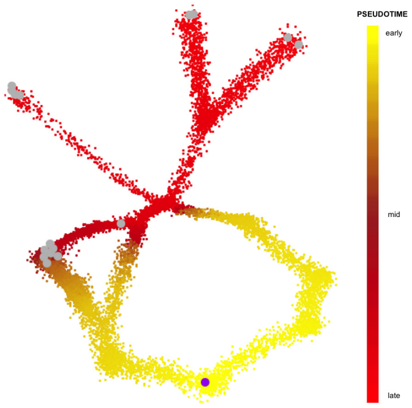

(d) Pseudotime: hitting distance

Figure 2: Pseudotime calculation strategies

##### 3.2 On numerical aspects of the solution

One of the crucial advantages of describing *pseudotime* as a solution of a linear system $Ax = b$ (Equation 4) is the possibility to measure the accuracy of our solution as the relative error $\frac{\|Ax-b\|}{\|b\|}$. Although iterative methods can solve the system (provided the solution exists) to any precision (limited by machine precision of floating point arithmetic), in practice the desired precision might not be possible to reach (e.g. because of an ill-conditioned system).

Although this happened rarely in our experiments, it is an issue worthy of attention. In most cases, non-convergence can be rectified by changing the kernel function (for instance, Gaussian kernel seems to work better than exponential kernel on data from flow cytometry) or adjusting the transformation of the data (an *asinh* transformation is usually a sensible choice).

To summarize, the reported error of the numerical solution for pseudotime values offers an important diagnostic tool in pseudotime estimation.

### 4 Topological clustering of random walks

Our approach to clustering random walks is based on the idea presented in Pokorny et al. (2014). Detailed descriptions of relevant mathematical concepts and tools (algebraic topology, persistent homology) can be found in Hatcher (2000); Edelsbrunner and Harer (2022).

#### 4.1 Basic notions of simplicial homology

For convenience, we define basic terms used throughout this document here.

**Definition 3** ($p$-simplex). *We define a* $p$-**simplex** *as a convex hull of* $p+1$ *affinely independent points.*

*A $p$-simplex $\sigma$ is a point for $p = 0$, a line segment for $p = 1$, a triangle for $p = 2$, a tetrahedron for $p = 3$, etc. We use the notation $\sigma = [v_0, v_2, \dots, v_p]$ where $v_i$ are vertices. The convex hull of a nonempty subset of vertices is called a* **face** *of the simplex.*

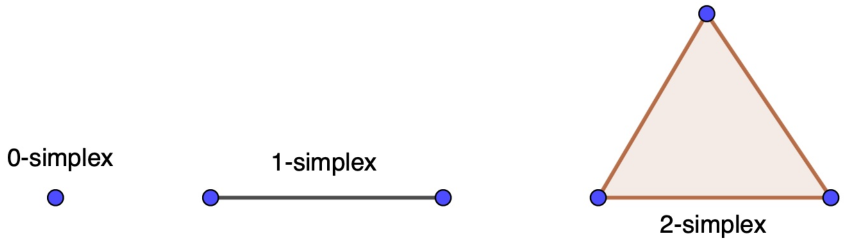

Figure 3: Simplices

**Definition 4** (Simplicial complex)**.** *A* **simplicial complex** $K$ *is a set of simplices that satisfies the following conditions:*

*1. Every face of a simplex from* $K$ *is also in* $K$*.*

*2. The non-empty intersection of any two simplices* $\sigma_1, \sigma_2 \in K$ *is a face of both* $\sigma_1$ *and* $\sigma_2$*.*

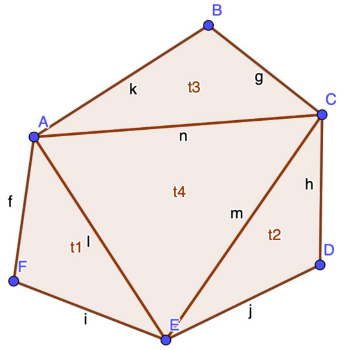

Figure 4: Simplicial complex

**Definition 5** ($p$-chain group)**.** *For a simplicial complex* $K$*, the group* $C_p(K)$ *of* $p$**-chains** *of* $K$ *is defined as a formal linear combination of* $p$*-simplices:*

$$C_p(K) = \left\{ \sum_i m_i \sigma_i | m_i \in \mathbb{Z}/2\mathbb{Z} \right\},$$

*where $\sigma_i$ are p-simplices of K.*

**Note** *that the coefficients $m_i \in \mathbb{Z}/2\mathbb{Z}$. In general, these coefficients may be from any ring. This choice simplifies many things (e.g. no issues with orientation arise, as $-1 = 1$), but we lose some descriptive power.*

**Definition 6** (Boundary)**.** *The* **boundary homomorphism** $\partial_p : C_p(K) \to C_{p-1}(K)$ *is defined simplex-wise for a simplex $\sigma = [v_0, v_2, ..., v_p]$ as*

$$\partial\sigma = \sum_{j=0}^{p} [v_0, \dots, v_{j-1}, \hat{v}_j, v_{j+1}, \dots, v_p],$$

*where $\hat{v}_j$ indicates deletion of vertex $v_j$.*

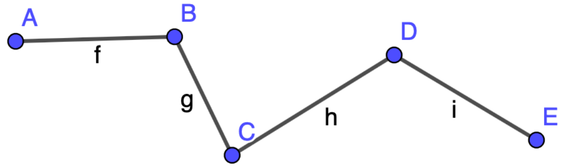

Figure 5: 1-chain $c = f + g + h + i = [A, B] + [B, C] + [C, D] + [D, E]$

$$\begin{aligned}\partial c = \partial f + \partial g + \partial h + \partial i &= \partial[A, B] + \partial[B, C] + \partial[C, D] + \partial[D, E] \\ &= A + B + B + C + C + D + D + E = A + E\end{aligned}$$

(coefficients in $\mathbb{Z}/2\mathbb{Z}$)

The boundary operator can be expressed as a matrix

$$\partial_{i,j} = \begin{cases} 1, & \text{if } \sigma_i \text{ is a co-dimension 1 face of } \sigma_j \\ 0, & \text{otherwise} \end{cases}$$

**Definition 7** (Reduced (boundary) matrix)**.** *Let R be a matrix with elements in $\mathbb{Z}/2\mathbb{Z}$ and $Low(j)$ be the row index of the lowest 1 in the column j of the matrix R and $Low(j)$*

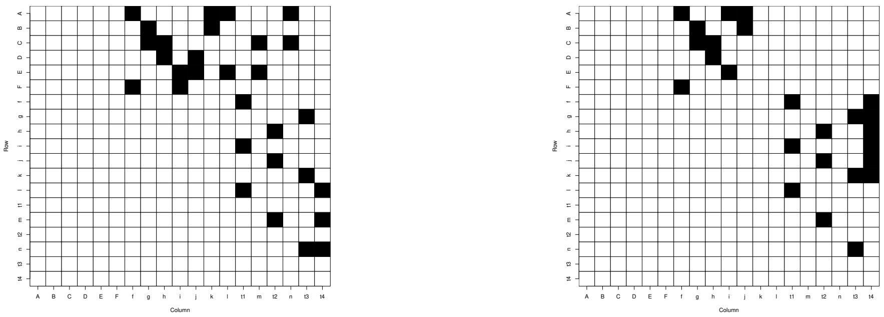

(a) Boundary matrix ⟶ (b) Reduced boundary matrix

Figure 6: Boundary matrix of simplicial complex from Figure 4, 1s are marked as black squares

*undefined on zero columns. We say that matrix $R$ is* **reduced** *if for any two non-zero columns $i$, $j$, $i \neq j$ $Low(i) \neq Low(j)$. See Figure 6.*

Matrix reduction can be performed by efficient algorithms and is essential for calculation of persistence (Definition 11). Matrix reduction corresponds to left-to-right column addition of columns representing boundaries of the same dimension, hence the columns of *reduced boundary matrix* correspond to boundaries of simplicial complexes (sums of boundaries of some $p$-simplices).

**Definition 8** (Boundaries & Cycles)**.** *We call*

$p$-**boundaries***: the group $B_p = Im\ \partial_{p+1}$*

$p$-**cycles***: the group $Z_p = Ker\ \partial_p$*

$p$-**th homology group**[2]*: $H_p = Z_p/B_p$*

**Definition 9** (Filtration)**. Filtration** *is a sequence of simplicial complexes that are ordered by inclusion. Hence, if $K_1$ and $K_2$ are complexes where $K_1$ appears before $K_2$ in the filtration,*

[2]Note that by fundamental lemma of homology $\partial_p \partial_{p+1} c = 0$ holds for any $p+1$-chain $c$ and consequently $B_p \subset Z_p$.

*then* $K_1 \leq K_2 : K_1$ *is a sub-complex of* $K_2$.

*By* **filtration of a simplicial complex** $K$ *we mean a sequence*

$$\emptyset = K_0 \subseteq K_1 \subseteq K_2 \subseteq \ldots \subseteq K_n = K$$

*These inclusions induce a sequence of homology groups connected by homomorphisms*

$$f_p^{i,j} : H_p(K_i) \to H_p(K_j),\ i \leq j$$

*Filtration thus gives a sequence of homology groups*

$$0 = H_p(K_0) \to H_p(K_1) \to H_p(K_2) \to \ldots \to H_p(K_n) = H_p(K).$$

**Definition 10** (Filtration induced by a monotonic function)**.** *Let* $K$ *be a simplicial complex and* $f : K \to \mathbb{R}$ *a monotonic function on* $K$*, that is* $f(\sigma) \leq f(\tau)$ *whenever* $\sigma$ *is a face* $\tau$*. Let* $a_1 < a_2 \ldots < a_n$ *be the values of function* $f$ *on the simplices in* $K$ *and* $a_0 = -\infty$*, then the sequence* $K_i = f^{-1}(-\infty, a_i]$ *is a* **filtration of** $K$ **induced by** $f$*. We call* $a_i$ **filtration values***.*

**Definition 11** (Persistence and essential classes)**.** *Given a filtration,* $p$**-th persistent homology groups** *are defined as* $H_p^{i,j} := Imf_p^{i,j}$*, for* $0 \leq i \leq j \leq n$*. (i.e.* $H_p^{i,j} = Z_p(K_i)/(B_p(K_j) \cap Z_p(K_i))$*).*

*We say that a homology class* $\gamma \in H_p(K_i)$ *is* **born** *at* $K_i$ *if* $\gamma \notin H_p^{i-1,i}$*. Furthermore we say, that a class* $\gamma$ *born at* $K_i$ **dies** *entering* $K_j$ *if* $f_p^{i,j-1}(\gamma) \notin H_p^{i-1,j-1}$ *but* $f_p^{i,j}(\gamma) \in H_p^{i-1,j}$*.*

*If the filtration is induced by a monotonic function we define the* **persistence** *of class* $\gamma$

*which was born at $K_i$ and died at $K_j$ as pers*$(\gamma) = a_j - a_i$. *Sometimes it is convenient to consider* **index persistence** $= j - i$.

*Throughout the text we will refer to the subcomplexes or corresponding filtration values where a homology class is born or where it dies as* **birth** *and* **death** *of the class, respectively.*

*We say that a p-cycle c represents an* **essential class** *in the simplicial complex $K$ iff there is no $(p+1)$-chain $k' \in C_{p+1}(K)$ such that $c = \partial k'$. In other words, essential classes are persistent homology classes which never die.*

If we consider a *reduced boundary matrix* with columns and rows ordered by filtration, then every non-zero column $j$ corresponds to a *death* of a homology class. Moreover, this class was *born* at $K_{\mathrm{Low}(j)}$, thus the *index persistence* of this homology class is $j - \mathrm{Low}(j)$.

##### 4.2 Triangulation of a point cloud & random walks

The triangulation of a high-dimensional point cloud is a computationally intensive task. The complexity of building triangulations used in lower-dimensional problems (such as *Delauney triangulation*) grows exponentially with the dimension, while other triangulations, better suited for high-dimensional problems (like the *Vietoris-Rips complex*) can become very large if we need sufficient cover of the space. We decided, therefore, to use the *witness complex* (Silva and Carlsson (2004)), implemented in Kachanovich (2015). *Witness complex* is built from a smaller set of landmark points and uses the original point cloud to witness the existence of connections between points. In our implementation we first cluster the points into a large number of clusters (625 by default) by $k$-means clustering, and use the centroids of the clusters as landmark points. Obviously, the vertices of *witness complex* differ from the vertices in $G'$ where the random walks were simulated. We need to triangulate the random walks as well. This procedure is straightforward: we simply contract the vertices to the corresponding cluster and remove loops from the new contracted random walks. Thus, after this *triangulation* step, each random walk corresponds to a connected 1-chain in the *witness complex*.

A witness complex can be built up to a preset dimension of simplices. Since we need only 2-simplices to classify 1-chains, we limit the dimension to 2, and effectively create a 2-skeleton. The filtration values of simplices are then calculated the same way as they are in alpha complex filtration (Rouvreau (2015)).

The choice of landmark points (clusters) can limit the resolution of the interactive aggregation of random walks. We did not encounter such an issue in practice and the simplicial complex built with default parameters always offered rich enough structure to achieve the desired level of detail. Potential issues can however be rectified by increasing the number of clusters or by introducing a custom clustering vector. Of note the triangulation of random walks takes place during the interactive session and the filtration can be recalculated independently of all other modules.

###### 4.2.1 Data denoising

Technical noise is always present in single-cell data. The presence of unwanted data points in sparse regions which are used for characterization of the shape of our point cloud can make the use of persistent homology less effective at describing our data. In order to reduce this noise and to help capture the topology correctly, we apply the denoising technique, suggested in Carlsson et al. (2008).

Taking advantage of a previously built $k$-NNG, we replace the coordinates of each data point with the averaged coordinates of some $l$ of its nearest neighbors. The denoised version of our expression matrix is used to construct the witness complex.

##### 4.3 Cycle representation

In order to classify random walks with respect to homology classes, we need a suitable representation of their dissimilarity. The basic idea of this approach is outlined in Pokorny et al. (2014). We, however, improve on the cited results by allowing for *essential classes*, and consequently the classification of random walks with different endpoints (e.g. developmental branching).

Every random walk can be understood as a connected 1-chain with at least one vertex in its boundary (the cell-of-origin vertex $v_0$) common to all other random walks. Consider first a set of 1-chains $\{c^0, c^1, \ldots, c^N\}$ with a common boundary (i.e. origins and endpoints of all corresponding triangulated random walks are the same). We can now fix a reference 1-chain (e.g. $c^0$) and consider 1-*cycles* $\{c^0 + c^1, c^0 + c^2, \ldots, c^0 + c^N\}$. If all cycles in this set are boundaries of some 2-chain, they can be uniquely expressed as linear combinations (over $\mathbb{Z}/2\mathbb{Z}$) of columns of the *reduced boundary matrix* (as described in Pokorny et al. (2014)). However, if some of these *cycles* represent an *essential class*, we need a more elaborate solution, implemented in Algorithm 1. If we encounter a *cycle* which is not a *boundary* during the calculation of the representation, we simply form an auxiliary *cycle* representing the "essential part" and add it as another column into the *reduced boundary matrix*. The filtration value corresponding to the *death* of this class is $\infty$, but for the purpose of visualization on the persistent diagram we assign arbitrary values to all *essential classes*, so that they have the same *death/birth* ratio and the *death* filtration value both higher than any non-essential classes.

If we fix a reference 1-chain $c^0$, we can uniquely represent this way any 1-chain $c^i$ with the same boundary by expressing the *cycle* $c^0 + c^i$ by a set of columns of (possibly updated) *reduced boundary matrix*. Note that $c^0 + c^0$ is an empty 1-chain. This representation will be used in subsection 4.4 to hierarchically cluster random walks.

What remains to be discussed is the case of classification of random walks with different endpoints. We implement two options. The first option is to simply choose the endpoint with highest *pseudotime*, set it as the endpoint of all random walks in question, and connect the other endpoint(s) to it by the shortest path within the simplicial complex. This approach is convenient in the case of closely clustered endpoints that represent the same developmental fate. The second option is to add a 1-simplex between selected endpoints. Since there were no 1-simplices connecting these endpoints (otherwise the random walks in question would end in the endpoint with the higher *pseudotime*), this will effectively create one or more new *essential classes*. Random walks differing by an *essential class* will be split into major branches at the root of the dendrogram (see subsection 4.4).

We close this subsection by a detailed description of the algorithms used and by providing a proof of correctness. Since we use the field $\mathbb{Z}_2$ as the coefficients ring, we can understand simplicial complexes (and consequently $p$-chains) as sets of simplices. To simplify further exposition we introduce following notation and arithmetic:

**Definition 12** (Notation and arithmetic of $p$-chains & $\text{Low}^{-1}$). *Given a simplicial complex* $K = \{\sigma_1, \sigma_2, \ldots, \sigma_N\}$ *and a p-chain* $c = \{\sigma_{i_1}, \sigma_{i_2}, \ldots, \sigma_{i_k}\}$*, we define the* **index representation** $c_I$ *of* $c$ *as the tuple* $c_I = (i_1, i_2, \ldots, i_K)$.

*For any two p-chains* $a_I$, $b_I$ *we define (in agreement with the arithmetic over* $\mathbb{Z}_2$*)*

$$a_I + b_I := a_I \triangle b_I \text{ (the symmetric difference of the 2 sets).}$$

*By a slight abuse of notation, we will define the* **index representation of a set of $p$-chains** $C = \{c^1, \ldots, c^k\}$ *as* $C_I := (c^1 + c^2 + \ldots + c^k)_I$.

*We will also assume that the indices in the index representation tuple are always ordered with respect to the filtration in descending order, i.e. for any p-chain c,* $c_I[1]$ *has the highest* *filtration value among all simplices forming the p-chain.*

*For the reduced boundary matrix $R$ let us further define the map $Low^{-1}$:*

*$Low^{-1}(i) = j$ if $i$ is the highest row index of a non-zero element in a column $j$ of matrix $R$ and $Low^{-1}(i) = -1$ if there is no such column.*

---

**Algorithm 1** One cycle representation

---

**Input:** $c_I$ cycle, $R$ reduced boundary matrix with corresponding $\mathrm{Low}^{-1}$ function
**Output:** $r$ representation (tuple of column indices) of the cycle $c_I$ as columns of (updated) boundary matrix $R$, updated boundary matrix $R$

```
procedure GET_REPRESENTATION_OF_CYCLE(c_I, R)
    r ← ∅
    e_I ← ∅
    b_I ← ∅
    N ← number of columns of R
    while c_I ≠ ∅ do
        j ← Low^{-1}(c_I[1])
        if j>0 then
            r ← r ∪ j
            c_I ← c_I + R[, j]_I
            b_I ← b_I + R[, j]_I
            e_I ← e_I
        else
            e_I ← e_I + c_I[1]
            c_I ← c_I + c_I[1]
            r ← r
        end if
    end while
    if e_I ≠ ∅ then
        R ← [R, e_I] (adding a column with 1s at positions e_I and 0s elsewhere)
        Low^{-1} ← update Low^{-1} function with respect to the new R.
        r ← r ∪ (N + 1)
    end if
    return r, R
end procedure
```

---

**Algorithm 2** Cycles representation

**Input:** $C = \{c_I^1, c_I^2, \ldots, c_I^L\}$ set of cycles, $R$ reduced boundary matrix with corresponding $\mathrm{Low}^{-1}$ function

**Output:** Representation $= \{r^1, r^2, \ldots, r^L\}$ representations (tuples of column indices of (updated) boundary matrix $R$) of cycles $C$

```
for i ∈ 1 . . . L do
    r^i, R ← GET_REPRESENTATION_OF_CYCLE(c_I^i, R)
end for
return Representation
```

**Proposition 4.1: Correctness of algorithms 1 and 2**

i) Algorithm 1 finds unique representation for any $p$-cycle $c_I$ in maximum of $\max(c_I)$ steps in the form $c_I = \hat{b}_I + \hat{e}_I$, where $\hat{b}_I$ is a $p$-boundary and $\hat{e}_I$ is a $p$-cycle representing essential class(es).

ii) Matrix $R$ is updated in the $i$-th step of algorithm 2 *iff* $\forall j : 0 \leq j < i$ the cycle $c_I^i + c_I^j$ represents an essential class ($c_I^0 := \emptyset$).

*Proof.* Let us denote the original reduced boundary matrix $R_0$ and the number of columns of $R_0$, $N_0$. Notice that for any $e_I$ constructed by algorithm 1 there was no column with lowest non-zero element at $\max(e_I)$ before the update, hence the matrix $R$ is always *reduced* and the columns remain linearly independent after the update(s). Consequently, every element in their span has a unique representation.

(i) Since $c_I[1] = \max(c_I)$ and if $j = \mathrm{Low}^{-1}(c_I[1]) > 0$ then $\max(R[,j]_I) = \max(c_I)$, the value $\max(c_I)$ is reduced at least by 1 in every iteration.

By construction $\emptyset = c_I + b_I + e_I$, hence $e_I$ constructed by algorithm 1 is a cycle. If $e_I \neq \emptyset$, the matrix $R$ is updated by a column containing a cycle representing an essential class. All cycles not representing an essential class can be expressed as a unique sum of boundaries and hence $e_I$ would be $\emptyset$.

Consider a representation $r$ corresponding to the cycle $c_I$. Let us decompose $r$ into disjoint sets $r = r_b \cup r_e$, such that $\min(r_e) > N_0$, $\max(r_b) \leq N_0$ and define $\hat{e}_I := R[, r_e]_I$, $\hat{b}_I := R[, r_b]_I$.

(ii) If there exists $c_I^j$, $0 \leq j < i$ such that the cycle $c_I^i + c_I^j$ does not represent an essential class, then by definition there exists a $p$-boundary $b$ such that $c_I^i = c_I^j + b$. Since $c_I^j$ has a representation within $R$ and $b$ can be expressed as sum of columns of $R_0$, no matrix update is necessary. Conversely, if $\forall 0 \leq j < i$ the cycle $c_I^i + c_I^j$ represents an essential class, then $c_I^i$ cannot be expressed in the terms of the columns of matrix $R$ and $e_I$ constructed by algorithm 1 is non-empty. □

##### 4.4 Topological hierarchical clustering of random walks

Consider a set of $L$ random walks and corresponding 1-chains $\{c^0, c^1, c^2, \ldots, c^L\}$. Let us select $c^0$ as a reference 1-chain (thus having trivial representation) and the representations obtained by algorithm 2 be $\{\emptyset, r^1, r^2, \ldots, r^L\}$. Each $r^i$ corresponds to a set of homological classes each with its own *birth* and *death* filtration values. If we set a threshold on the *death* value and delete all homological classes which *died* before this point from the representation, a natural hierarchical clustering stems from the following idea. Initially, each leaf of the dendrogram contains all random walks with the same representation, then every time a homological class dies as we are increasing the threshold, two leaves/branches of the dendrogram may merge. As soon as all[3] homological classes *die*, the dendrogram is complete.

In practice, we proceed differently. We first choose (using an interactive persistence diagram) homology classes we deem significant based on their persistence. Then we order the homology classes present in any representation in decreasing order with respect to

[3]In practice we set threshold on column indices of corresponding (updated) *reduced boundary matrix.* Thus even *essential classes die* after a finite number of steps

their *death* value. Finally, we split the random walks into two branches[4] based on the presence/absence of the homology class with highest *death* value in their representation. The class with the highest *death* value is dropped and we re-iterate the process on each of the two branches.

#### 5 Connectome

First, a multigraph is created from vertices corresponding to clusters. One directed edge is added between any pair of vertices per each traversal by a random walk. Self-loops are deleted. Then, a new graph is plotted where the edges with the same direction between same vertices are merged, setting edge weights to the number of edges before merging (by default, only the majority-direction edge is kept for any pair of vertices). Vertices of the resulting graph are plotted as pie charts of cell populations present in each cluster. The graph layout employs the same 2D representation(s) as *tviblindi* GUI to facilitate the interpretation. Terminal nodes are automatically detected as vertices with fewer outgoing than incoming edges. The widths of edges represent their weights.

Although connectome offers only a low resolution overview of the dataset, the result is informative even for very complex topologies (Figure 7). On this dataset connectome was run as `Connectome(tv1,qq=0.8)` i.e. only arrows with width above the 80% percentile were plotted.

#### 6 Graphical user interface (GUI)

An essential part of *tviblindi* is GUI which enables the on-demand aggregation of random walks and real-time resolution adjustments. In this section we describe the main features

[4]If one branch is empty, we drop the class and re-iterate

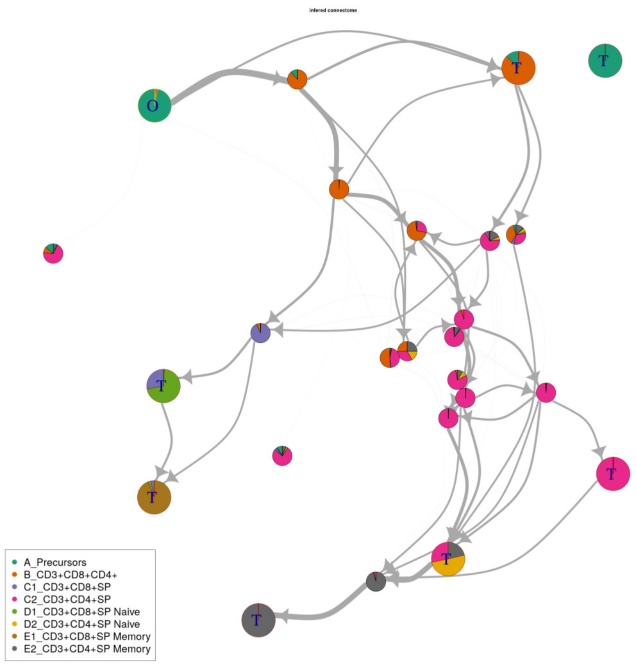

Figure 7: Connectome applied on the human thymus dataset (subsection 1.2)

of this GUI and the analysis process. (See also supplementary videos.) In this text we will refer to the annotated screenshots of the GUI on pages 33 and 34 of this document by numbers in circles, e.g. ①, and squares, e.g. [1], respectively. Please consult section 7 or the documentation of the *tviblindi* package for more details on any function mentioned in the text (function names are typeset as `verbatim`). Note that pop-up tooltips are displayed for button in the GUI when running.

The user usually starts with adjusting visualization and selecting a path model. As every run of the function `DimRed` adds a 2D layout to the *tviblindi* analysis object, we can choose the most appropriate visualization in the drop-down menu ①. Menu ② allows the user to choose the path model (pseudotime and random walks model, for different choice of cell-of-origin, different parameter choices for *pseudotime* calculation etc.). The 2-dimensional layout plot is optimized for large datasets (millions of events), but button

(9) can increase point size when needed. Labels of points (7) serve mainly for visualizing labeled populations on the plot invoked by (8), and for tracking population frequencies along the trajectory ((21) second pane).

This first step of the analysis itself is the choice of endpoints (3). The user selects some black dots in the layout via rectangular selection at once, or iteratively by pressing (4) after each selection. The selected endpoints are listed (as row indices in the data matrix) together with the number of random walks which end in a particular endpoint. Pressing (4) moves the endpoints in the current rectangular selection (*Selected terminal nodes* frame) to *Terminal nodes marked for further analysis*.

Once the user is satisfied with the endpoints selection, they can calculate their representation by pressing (5). Checkbox (6) switches between the two options for the multiple endpoints treatment (see subsection 4.3).

At this point the endpoints of interest are selected and the representation of all random walks ending in any of these endpoints are calculated. The user can then switch to the pane *Homology classes by persistence selection* [1] and select homology classes they deem significant, based on their persistence. By default, the $y$ axis of the diagram corresponds to the *death/birth* ratio[5]. This ratio describes the significance of the hole (sparse region) corresponding to the homology class. The larger this value, the sparser the interior of the hole relative to the density of its boundary, and the higher the significance for the random walk classification. The $x$ axis corresponds to the *death* value of each homology class plotted: this value corresponds to the *size* of the corresponding hole, and to the height of the branching on the dendrogram [4]/(13). For the artificial data used on the screenshots, the situation is clear: there are two *essential classes*, one corresponding to the larger hole in

[5]This gives better resolution for homology classes with large differences in size than the more standard *death - birth*. This can be changed using checkbox [3], resulting in a common persistence diagram.

the data and one created by the selection of two disparate developmental fates[6]. Another significant homology class corresponds to the smaller hole in the data.

The selection process is equivalent to the selection of endpoints, and by pressing button [2] the dendrogram [4] is updated.

Using the dendrogram (13), the user can select leaves/branches to aggregate random walks into trajectories. Two trajectories can be built at the same time by selecting a trajectory group A or B (11). The selection process is the same as before: pressing the "+" button (14) adds selected walks into the selection into the active trajectory, the "fire" button resets the trajectory. Note the slider (23), which adjusts the complexity of the dendrogram by setting a threshold on the minimal fraction of random walks for a node to be shown. Nodes below this threshold are merged.

The user can adjust the level of detail by selecting significant homology classes in [1]. In some cases it is more convenient to focus on a smaller subset of random walks. This can be achieved by pressing the button (17): this makes the analysis focus only on random walks in the active trajectory and the user can analyze its topology in detail, e.g. if inspecting finer branching processes. To go back to the entire set of random walks, the user can press the "fire" button (4) and start over with selection of endpoints.

Selected trajectories are visualized in a 2-dimensional embedding (layout) in pane (18). Change in signal from specific markers along a trajectory can be tracked (21). The development of those markers is shown as a line plot connecting the average values of a specific marker (or points on specific random walks, see below [5]) within a pseudotime segment. By default, the pseudotime values are divided into uniform segments (each containing the same number of points) whose number can be set changed in (20). To focus on either early or late stages of the development, adjust the pseudotemporal scale using (19): the

[6]Note that *essential classes* are infinite from the point of view of homology and their *death* values are chosen arbitrarily for visualization purposes

higher the value, the more resolution is put on early stages, and vice-versa. If the user is interested in *where* in the layout a certain part of development is projected, they can select the segments of interest and press (22). If multiple markers are tracked, the points of all trajectories in the selected segments are highlighted. If only one marker is tracked [5], we can see individual random walks and highlight only points from selected walks. If the user wants to manually clean a selected trajectory based on the evolution of the signal levels from a specific marker, they can manually select some random walks and remove them from the trajectory by pressing the ”flash” button near [6] ”flag”.

It is often useful to export the trajectories/selected points for further analysis. Highlighted points can be pinned by pressing the ”pin” button (24) (the ”thrash” deletes all pinned points). Trajectories can be pinned using (15). When pressing the ”pin” button, the user is prompted to name the trajectory or group of points. Pinned results are stored as list slots in the *tviblindi* object or can be exported as an extended FCS file by pressing the ”download” button (16) (the ”trash bin” button clears pinned trajectories). Once pressed, the user is prompted to name the exported FCS file and the results are stored either in a new FCS file built from the data matrix or added to an existing FCS file (if provided: see section section 7). The following is stored in the FCS file: the active 2D embedding, pseudotime values from the active ”PATH MODEL”, all sets of labels, all pinned trajectories and all pinned highlighted points.

Button (11) saves the corresponding figure as either a PNG or an SVG file (12).

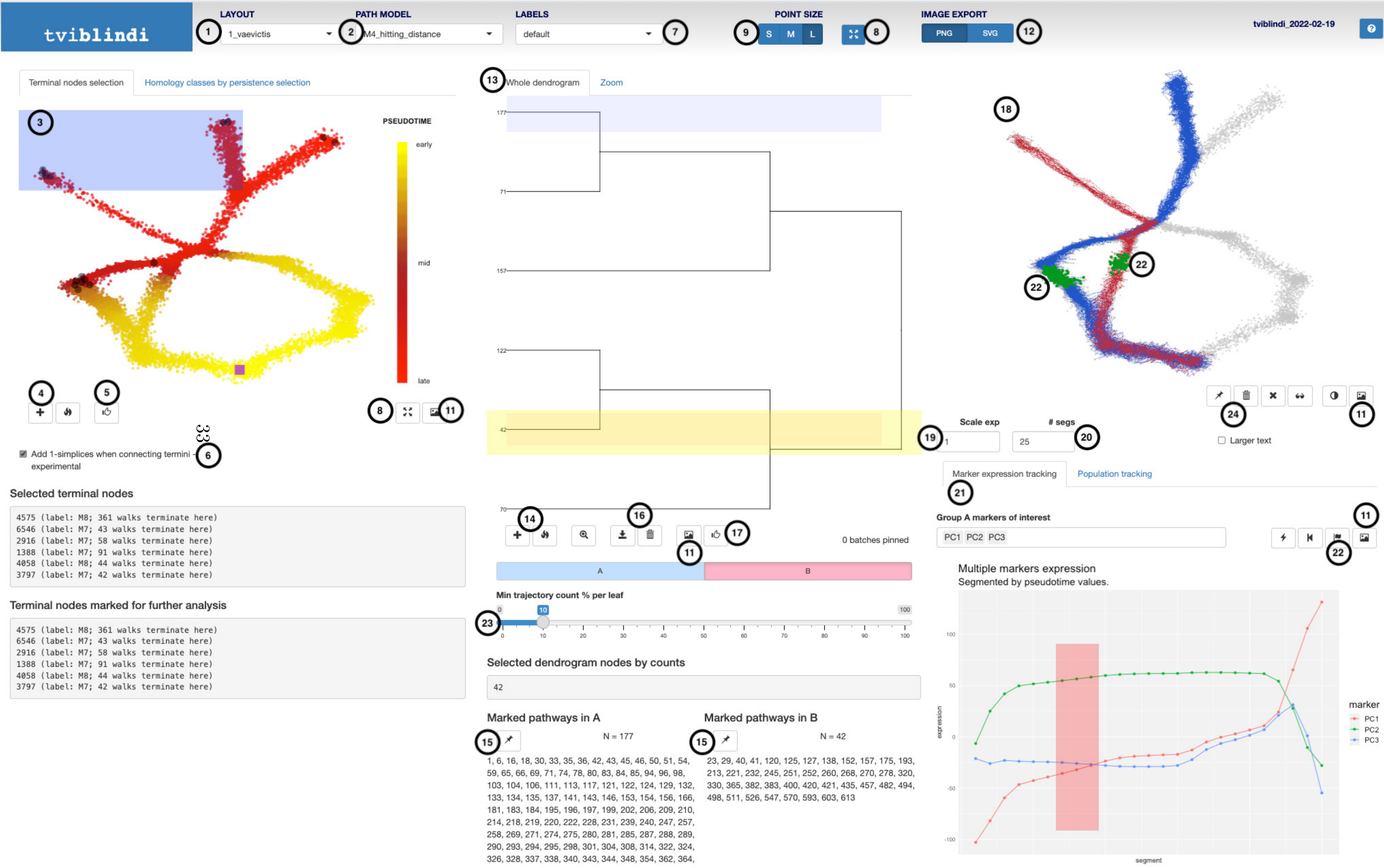
tviblindi
LAYOUT
1_vaevictis
PATH MODEL
M4_hitting_distance
LABELS
default
POINT SIZE
S M L
IMAGE EXPORT
PNG SVG
tviblindi_2022-02-19
Terminal nodes selection
Homology classes by persistence selection
PSEUDOTIME
early
mid
late
Add 1-simplices when connecting termini - experimental
Selected terminal nodes
4575 (label: M8; 361 walks terminate here)
6546 (label: M7; 43 walks terminate here)
2916 (label: M7; 58 walks terminate here)
1388 (label: M7; 91 walks terminate here)
4058 (label: M8; 44 walks terminate here)
3797 (label: M7; 42 walks terminate here)
Terminal nodes marked for further analysis
4575 (label: M8; 361 walks terminate here)
6546 (label: M7; 43 walks terminate here)
2916 (label: M7; 58 walks terminate here)
1388 (label: M7; 91 walks terminate here)
4058 (label: M8; 44 walks terminate here)
3797 (label: M7; 42 walks terminate here)
Whole dendrogram
Zoom
0 batches pinned
A
B
Min trajectory count % per leaf
Selected dendrogram nodes by counts
42
Marked pathways in A
N = 177
1, 6, 16, 18, 30, 33, 35, 36, 42, 43, 45, 46, 50, 51, 54, 59, 65, 66, 69, 71, 74, 78, 80, 83, 84, 85, 94, 96, 98, 103, 104, 106, 111, 113, 117, 121, 122, 124, 129, 132, 133, 134, 135, 137, 141, 143, 146, 153, 154, 156, 166, 181, 183, 184, 195, 196, 197, 199, 202, 206, 209, 210, 214, 218, 219, 220, 222, 228, 231, 239, 240, 247, 257, 258, 269, 271, 274, 275, 280, 281, 285, 287, 288, 289, 290, 293, 294, 295, 298, 301, 304, 308, 314, 322, 324, 326, 328, 337, 338, 340, 343, 344, 348, 354, 362, 364,
Marked pathways in B
N = 42
23, 29, 40, 41, 120, 125, 127, 138, 152, 157, 175, 193, 213, 221, 232, 245, 251, 252, 260, 268, 270, 278, 320, 330, 365, 382, 383, 400, 420, 421, 435, 457, 482, 494, 498, 511, 526, 547, 570, 593, 603, 613
Scale exp
### segs
Larger text
Marker expression tracking
Population tracking
Group A markers of interest
PC1 PC2 PC3
Multiple markers expression
Segmented by pseudotime values.
expression
segment
marker
PC1
PC2
PC3

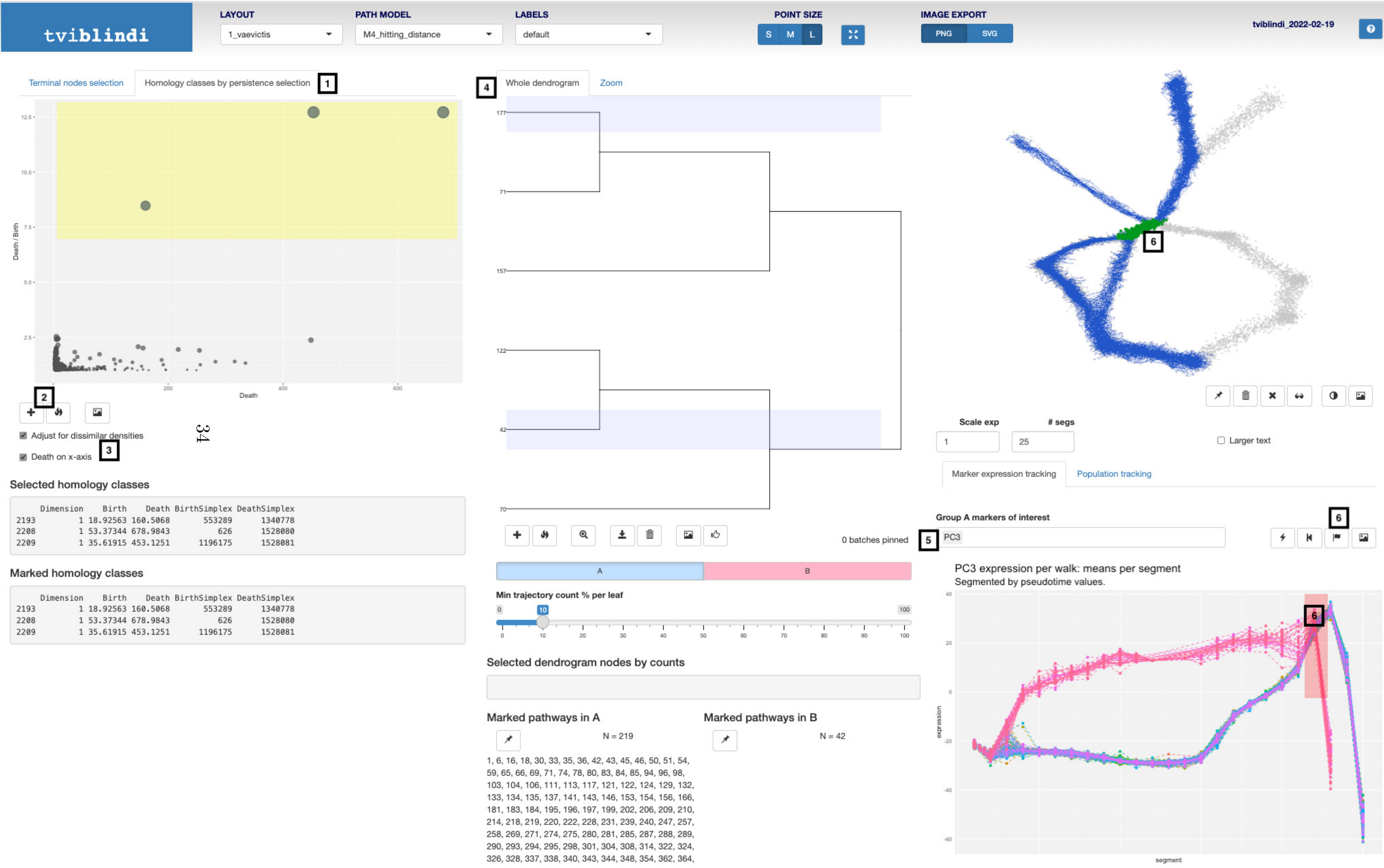

tviblindi
LAYOUT
1_vaevictis
PATH MODEL
M4_hitting_distance
LABELS
default
POINT SIZE
S
M
L
IMAGE EXPORT
PNG
SVG
tviblindi_2022-02-19
Terminal nodes selection
Homology classes by persistence selection
1
Death / Birth
Death
2
Adjust for dissimilar densities
Death on x-axis
3
Selected homology classes
Dimension Birth Death BirthSimplex DeathSimplex
2193 1 18.92563 160.5068 553289 1340778
2208 1 53.37344 678.9843 626 1528080
2209 1 35.61915 453.1251 1196175 1528081
Marked homology classes
Dimension Birth Death BirthSimplex DeathSimplex
2193 1 18.92563 160.5068 553289 1340778
2208 1 53.37344 678.9843 626 1528080
2209 1 35.61915 453.1251 1196175 1528081
4
Whole dendrogram
Zoom
0 batches pinned
A
B
Min trajectory count % per leaf
Selected dendrogram nodes by counts
Marked pathways in A
N = 219
1, 6, 16, 18, 30, 33, 35, 36, 42, 43, 45, 46, 50, 51, 54, 59, 65, 66, 69, 71, 74, 78, 80, 83, 84, 85, 94, 96, 98, 103, 104, 106, 111, 113, 117, 121, 122, 124, 129, 132, 133, 134, 135, 137, 141, 143, 146, 153, 154, 156, 166, 181, 183, 184, 195, 196, 197, 199, 202, 206, 209, 210, 214, 218, 219, 220, 222, 228, 231, 239, 240, 247, 257, 258, 269, 271, 274, 275, 280, 281, 285, 287, 288, 289, 290, 293, 294, 295, 298, 301, 304, 308, 314, 322, 324, 326, 328, 337, 338, 340, 343, 344, 348, 354, 362, 364,
Marked pathways in B
N = 42
6
Scale exp
### segs
1
25
Larger text
Marker expression tracking
Population tracking
Group A markers of interest
PC3
5
6
PC3 expression per walk: means per segment
Segmented by pseudotime values.
expression
segment
6

### 7 Running the analysis

The analytical pipeline consists of several steps which are not completely interdependent and can be (re)run in a different order if necessary (see following graph of the dependence structure (a dashed connection shows that the denoising step is optional).

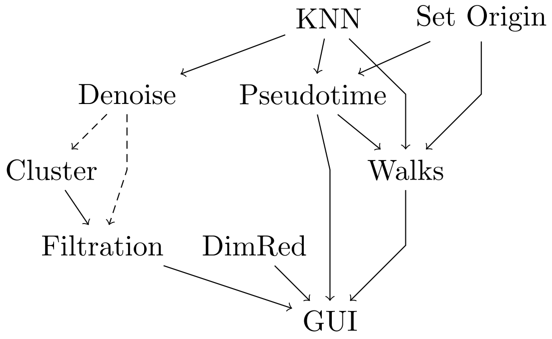

The *R* code needed to run a generic *tviblindi* analysis is shown below.

```
tv1 <- tviblindi(
    data = d,       # matrix of row-wise data, with named columns
    labels = l,     # vector of string labels per each row of d
    fcs = p,        # path to FCS file if d comes from its expression matrix
    events_sel = i  # indices of rows of FCS file, if only a subset is used
)
Set_origin(
    tv1,
    label = o,      # name of progenitor population
    origin_name = n # path model name (new)
)
KNN(tv1)            # build a k-NN graph
Denoise(tv1)        # reduce noise before witness complex construction
```

```
Cluster(tv1) # k-means clustering by default: k = 625 clusters
Filtration(tv1)     # build witness complex
DimRed(tv1)         # dimensionality reduction
Pseudotime(
    tv1,
    origin_name = n # path model name (existing)
)
Walks(
    tv1,
    N = 5000,       # number of finite random walks to simulate
    origin_name = n # path model name (existing)
)

launch_shiny(tv1)   # launch GUI for interactive analysis
```

All functions starting with a capital letter are generic S3 methods and the documentation could be accessed via the class suffix, e.g. `?KNN.tviblindi`. The parameter `origin_name` is optional and serves to distinguish different choices of the cell-of-origin, pseudotime calculation and random walk simulation (different path models; see section 6). The first command deserves clarification. The `fcs`, `events_sel` parameters are optional and connect the *tviblindi* object to an existing FCS file to be 'enhanced' as a part of the analysis. If omitted, the GUI will create a new FCS file out of the data matrix if the user chooses to export an FCS file. Otherwise if these parameters are provided, analysis results will be added to an existing FCS file as additional columns. If only a subset of rows from the FCS file is used for the analysis, the parameter `events_sel` records the indices of these rows as an integer vector.

### 8 Performance evaluation

#### 8.1 Sensitivity to hyperparameters

##### 8.1.1 Number of nearest neighbors

The pseudotime calculation is robust with respect to the number of nearest neighbors considered. The *Spearman correlation coefficient* between pseudotime estimated using 10 nearest neighbors and 50 nearest neighbors was 0.9986367 for the data from subsection 1.2.

The choice of the number of nearest neighbors $k$ used for *random walks* simulation has significant effect on the number of detected endpoints. Lower $k$ leads to higher number of endpoints, while higher $k$ tends to smooth the topology of the graph leading to more global view. Since the *random walks* simulation does not have any effect on the results of other modules (section 7), the user can experiment with this setting very efficiently. But even when larger $k$ causes spurious connection between unrelated population of cells, the topological clustering of *random walks* allows for the separation of different trajectories.

##### 8.1.2 Relaxation parameter for *witness complex* construction

The relaxation parameter `alpha2` (function `Filtration`) controls the size of the simplicial complex. Lower values of `alpha2` lead to smaller simplicial complex which might not give sufficient resolution for the downstream analysis. On the other hand, smaller simplicial complexes lead to significant speedup.

*tviblindi* is highly robust with respect to this parameter. Even with a drastic reduction of `alpha2`, the loss of resolution is not severe. An example of the effect of setting `alpha2` to a very low value is shown in Figure 8. When this parameter is set to 0.001 (compared to the default estimated value of $\sim 6$), the resulting simplicial complex consists of approximately $\sim 25000$ simplices. This leads to the emergence of essential classes (Figure 8b, red rectangle). Consequently, the hierarchical structure in the red group of trajectories (Figure 8c, Figure 8e) is lost - the trajectories in the red group are separated by essential classes, all of which have the same (infinite) size. However, the resolution for the majority of trajectories remains sufficient. Over-reduction of this parameter can be diagnosed by detecting a large number of essential classes. For the purpose of trajectory classification, we recommend the use of a large complex. However, a smaller complex (which still captures the topology well) may be useful for higher-order analysis, such as the Hodge decomposition of vector fields (e.g., RNA velocity).

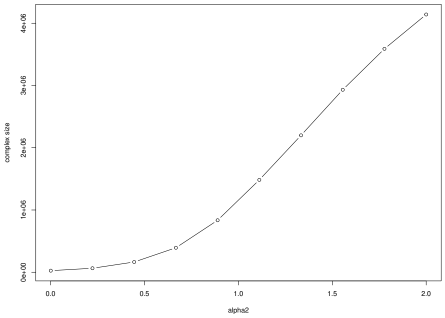

(a) Complex size vs. alpha2

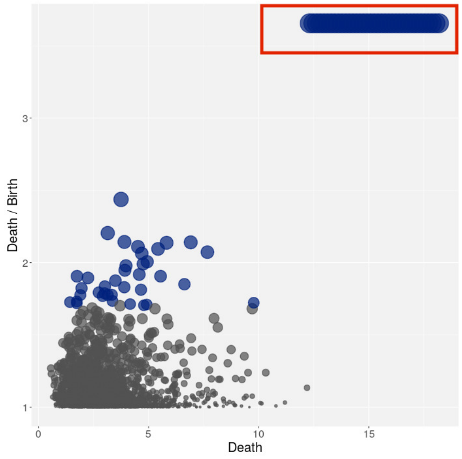

(b) Persistence diagram with essential classes (red rectangle)

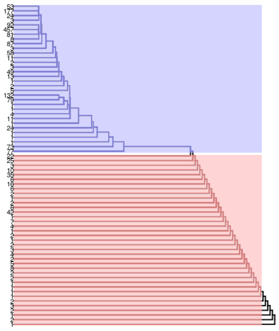

(c) Dendrogram of trajectories - red part aggregated with respect to essential classes

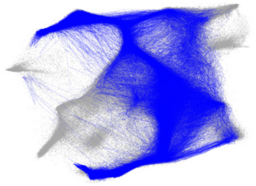

(d) Blue group of trajectories

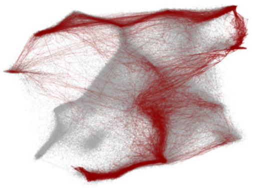

(e) Red group of trajectories

Figure 8: Relaxation parameter alpha2

The default setting is very conservative and creates very large simplicial complex (roughly 1-2 orders of magnitude larger than necessary, according to our tests) while maintaining feasible running times (subsection 8.3).

##### 8.2 Comparison to state-of-the-art methods

We evaluate how *tviblindi* performs in comparison to selected state-of-the-art TI methods. As we focus on analysis of large single-cell datasets, we have chosen methods which scored well in previously published benchmarks, and scale well with sample size Saelens et al. (2019); Stassen et al. (2021). Namely we compared *tviblindi* to *Monocle3* (v1.3.1) Cao et al. (2019), *Stream* (v1.1) Chen et al. (2019), *Palantir* (v1.0.0) Setty et al. (2019), *VIA* (v0.1.89) Stassen et al. (2021), *StaVia* (Via 2.0) Stassen et al. (2024), *CellRank 2* (v2.06) Weiler et al. (2024) and *PAGA* (scanpy==1.9.3) Wolf et al. (2019). All methods were run with either default or recommended setting (when available) for the corresponding dataset. When the implementation enabled a custom embedding, the final output was projected using *vaevictis* projection used in the main text to facilitate the interpretation. For our comparison we use datasets which were thoroughly analyzed by *tviblindi* in the main text.

##### 8.2.1 *tviblindi* on up-sampled *dyntoy* dataset

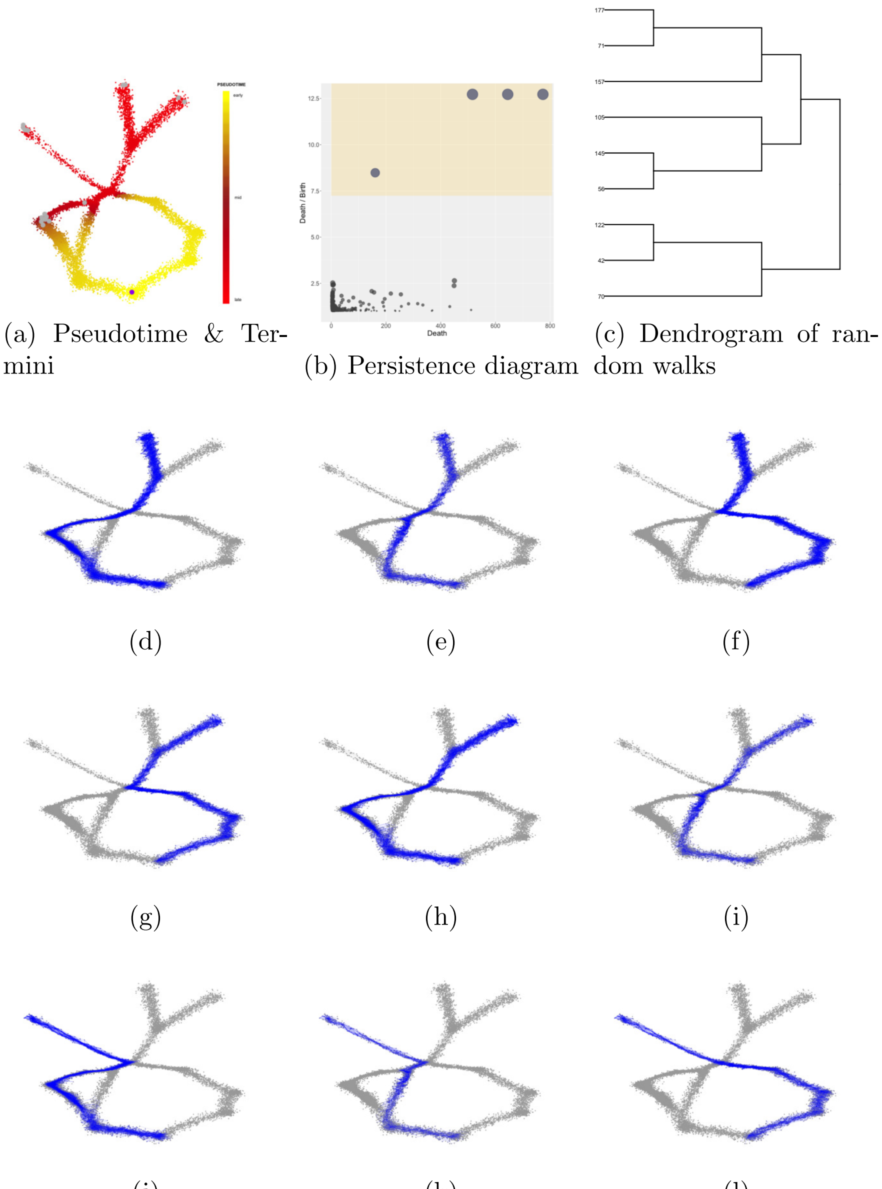

(a) Pseudotime & Termini

(b) Persistence diagram

(c) Dendrogram of random walks

(d)

(e)

(f)

(g)

(h)

(i)

(j)

(k)

(l)

Figure 9: *tviblindi* applied on up-sampled *dyntoy* dataset

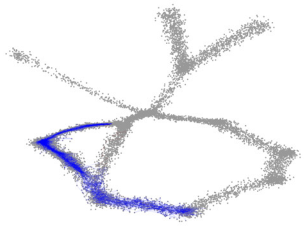

Figure 10: 4% of random walks ended in the up-sampled population

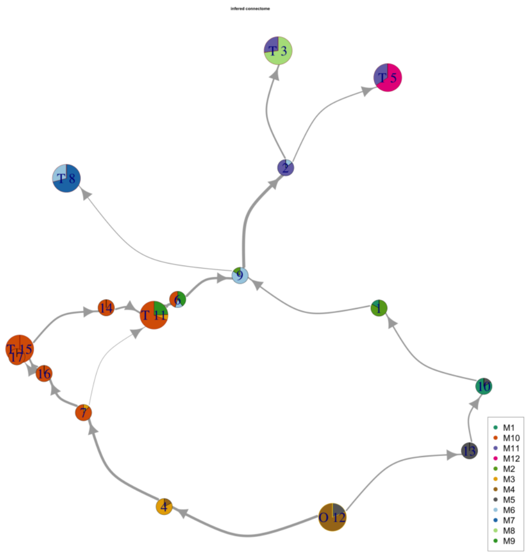

Figure 11: Connectome applied on up-sampled dyntoy dataset

###### 8.2.2 Monocle 3

*Monocle 3* relies heavily on the dimensionality reduction. The drawbacks of this approach become apparent when we apply *Monocle 3* on the artificial dataset while using the default method *UMAP* (Figure 12, Figure 13). The artifacts (e.g. over-separation of populations) introduced by the dimensionality reduction may hamper the interpretability of the result. On the other hand if we use *vaevictis*, *Monocle 3* captures the basic skeleton of dataset and misses only some connections (Figure 14). At the same time the inference of pseudotime based on the graph representation is very robust with respect to uneven distribution of cells when using appropriate dimensionality reduction (Figure 15). When applied the biological dataset the shortcomings are more subtle. In the subsection 1.2 dataset *Monocle* does not detect $\beta$ selection which is supported by low number of events. On the other hand, it over-complicates the interpretation of the dynamics within large CD4 SP and CD8 SP populations Figure 16. *Monocle 3* offers interactive interrogation of the *principal graph* and calculation of *pseudotime* along selected branches (not shown). However the resolution is set beforehand by the dimensionality reduction, clustering and graph construction and cannot be adjusted during the interactive section. In contrast to *tviblindi* which keeps the level of resolution on a single *random walk* and uses the structure of large simplicial complex for on-demand aggregation.

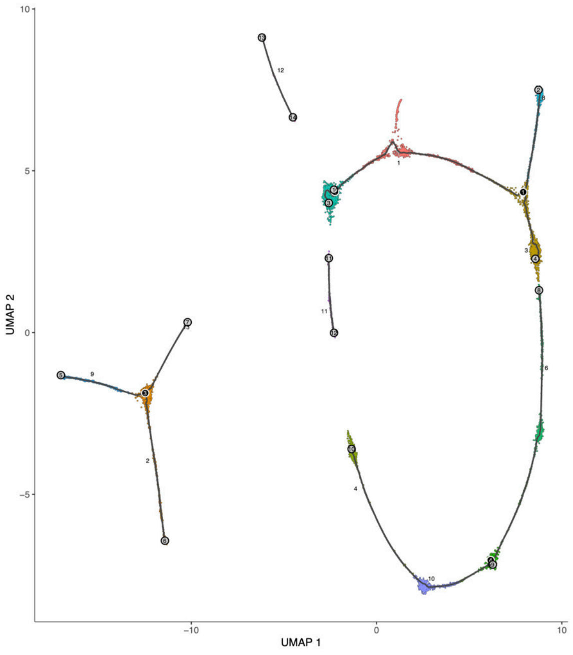

Figure 12: *Dyntoy* dataset subsection 1.1, *UMAP* dimensionality reduction

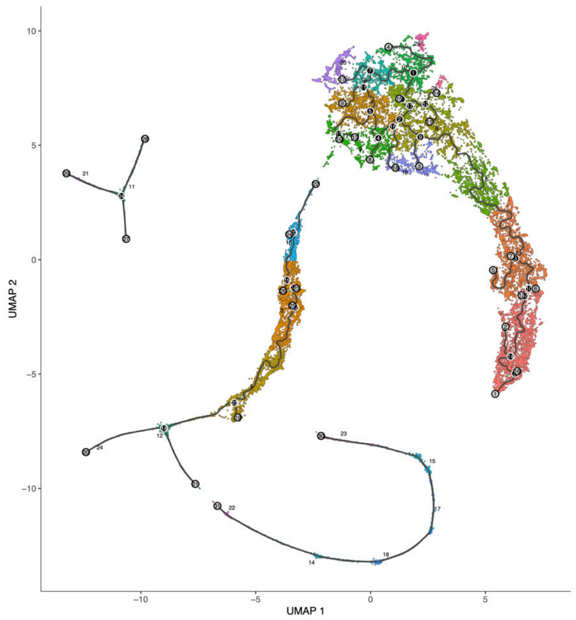

Figure 13: Up-sampled *dyntoy* dataset subsection 1.1, *UMAP* dimensionality reduction

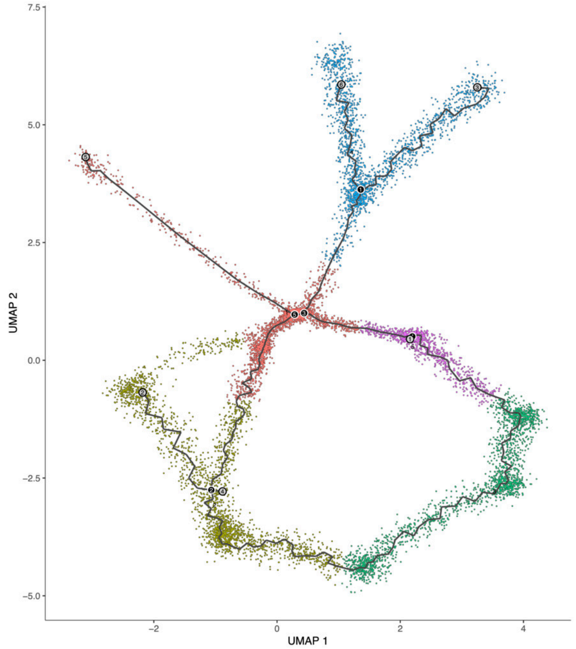

Figure 14: *Dyntoy* dataset subsection 1.1, *vaevictis* dimensionality reduction

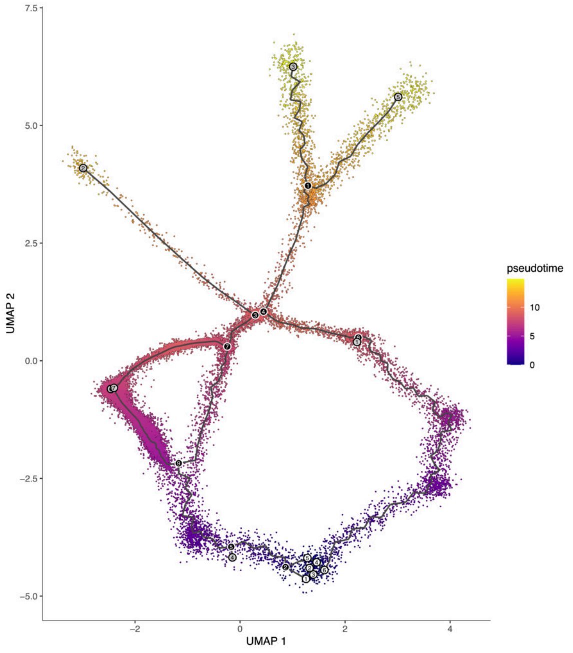

Figure 15: Up-sampled *dyntoy* dataset subsection 1.1 with inferred pseudotime, *vaevictis* dimensionality reduction

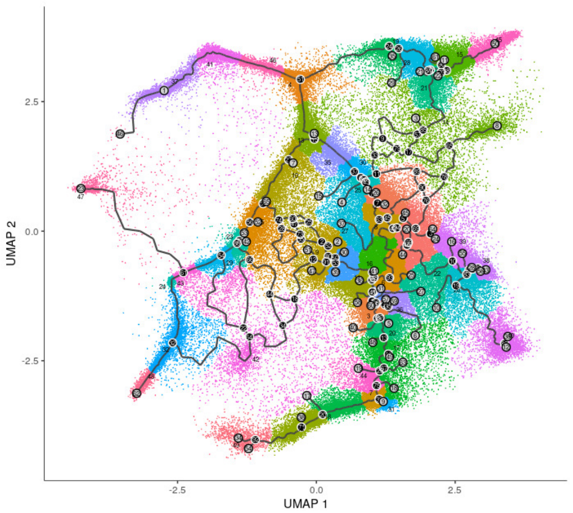

Figure 16: Human thymus & PBMC subsection 1.2, *vaevictis* dimensionality reduction

##### 8.2.3 Stream

Since the underlying structure of the method is a *minimal spanning tree*, *Stream* is not well suited for the analysis of non-tree trajectories (Figure 17, Figure 18). *Stream* successfully discovers developmental terminal points (Progenitors, apoptotic cells and CD4 SP/CD8 SP in peripheral blood) in the human T-cell development data. However the discovered structure is too simplistic and skewed to the most abundant population (immature CD4 SP thymocytes) (Figure 19).

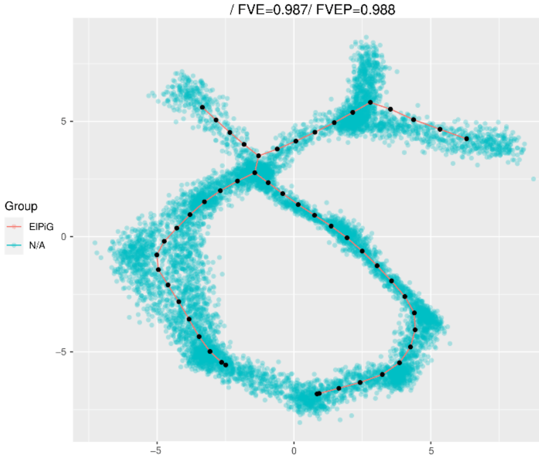

Figure 17: *Dyntoy* dataset subsection 1.1

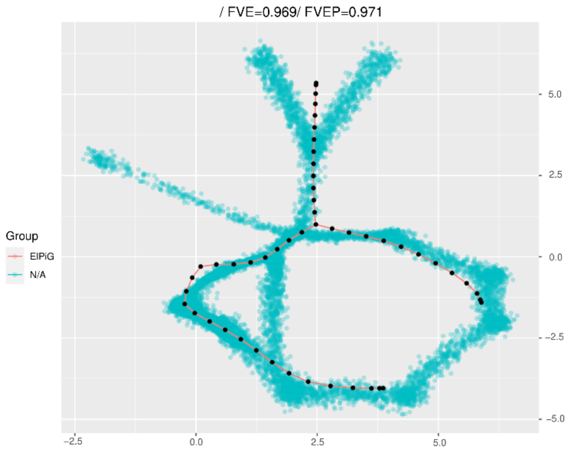

Figure 18: Up-sampled *dyntoy* dataset subsection 1.1

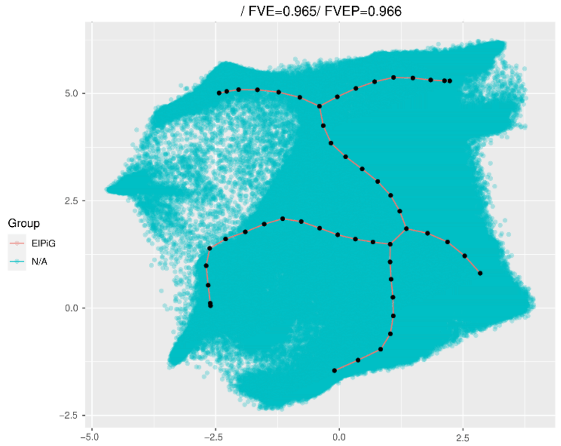

Figure 19: Human thymus & PBMC subsection 1.2

##### 8.2.4 Palantir

*Palantir* is very resilient to the uneven distribution of the cells in all datasets when calculating the pseudotime. It correctly discovers candidate endpoints in the biological dataset (Figure 22) but lacks the toolbox to interrogate subtle features such as complex branching. Although the pseudotime is inferred correctly even in the artificial dataset, *palantir* struggles to detect some of the endpoints (Figure 20, Figure 21).

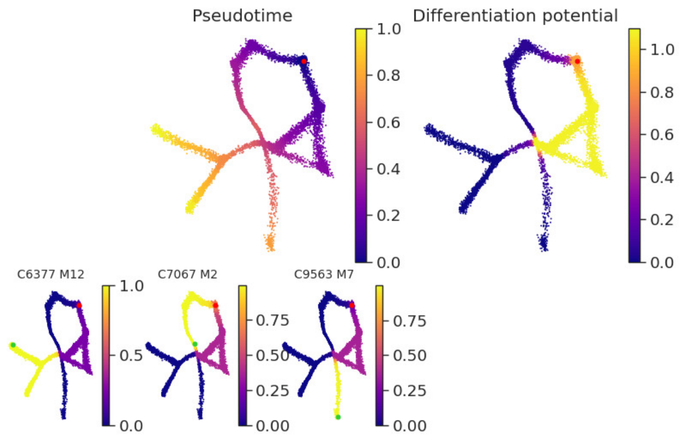

Figure 20: *Dyntoy* dataset subsection 1.1

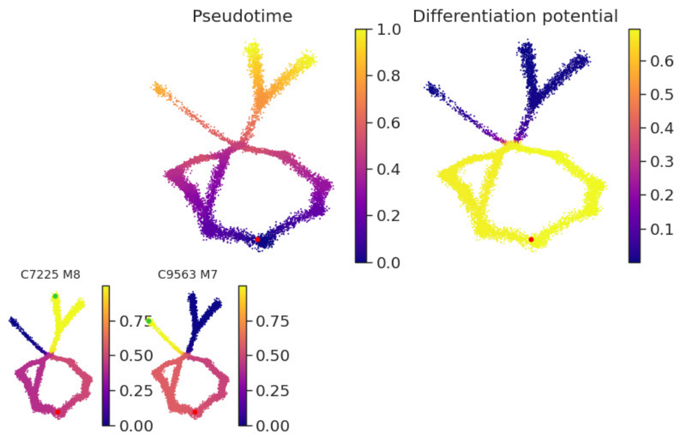

Figure 21: Up-sampled *dyntoy* dataset subsection 1.1

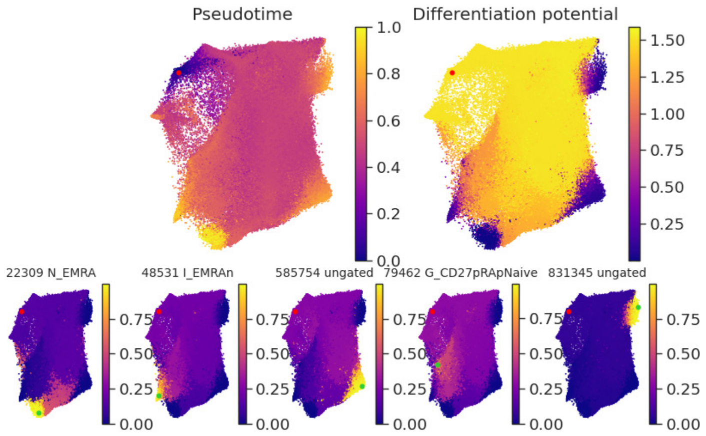

Figure 22: Human thymus & PBMC subsection 1.2

###### 8.2.5 VIA

*VIA* is the only of the tested method with running times comparable to *tviblindi* (see subsection 8.3). It performs relatively well on artificial datasets (Figure 23, Figure 24), although it sometimes misses obvious connections. It is also susceptible to the aforementioned issues with unevenly distributed cells, giving the highest pseudotime value to the over-abundant population *M10* (Figure 25). In the case of the biological dataset *VIA* is able to detect candidate endpoints but fails to discover subtle features (such as $\beta$ selection; Figure 25). The lack of an interactive interrogation or resolution adjustment of the results makes *VIA* difficult to use on complex datasets.

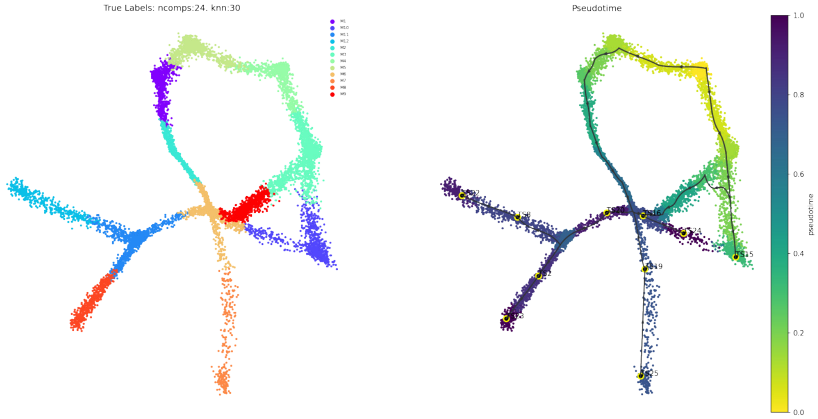

Figure 23: *Dyntoy* dataset subsection 1.1

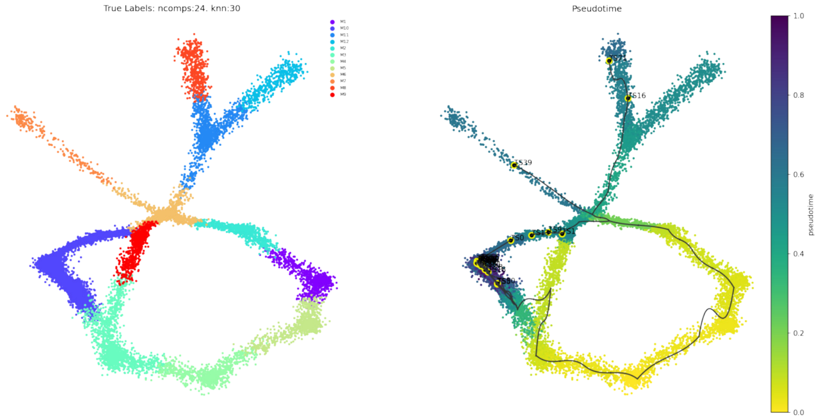

Figure 24: Up-sampled *dyntoy* dataset subsection 1.1

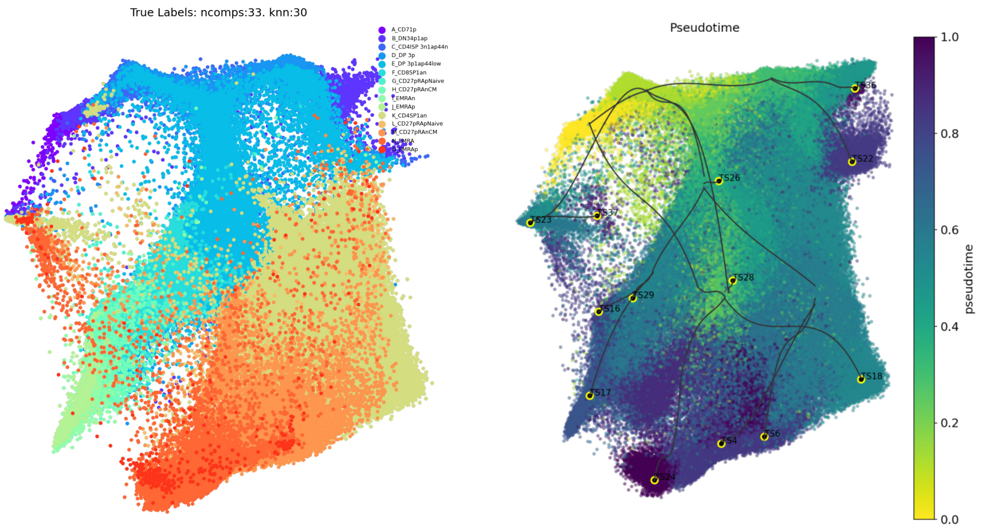

Figure 25: Human thymus & PBMC subsection 1.2

###### 8.2.6 PAGA

*PAGA* performs well on the artificial dataset (Figure 26). However when applied on the up-sampled dataset, the result is surprisingly bad (Figure 27). This is especially surprising when compared with the result of the connectome tool (Figure 11). Our implementation uses the same clustering approach as *PAGA* and we used the same number of nearest neighbors in our analysis. Our approach (inspired by *PAGA*) should improve the result only by the introduction of better layout, directionality & automatic terminal node detection. However the resolution should be comparable.

In general the lack of sensible layout and the lack of terminal node detection makes *PAGA* difficult to interpret on complex datasets (Figure 28).

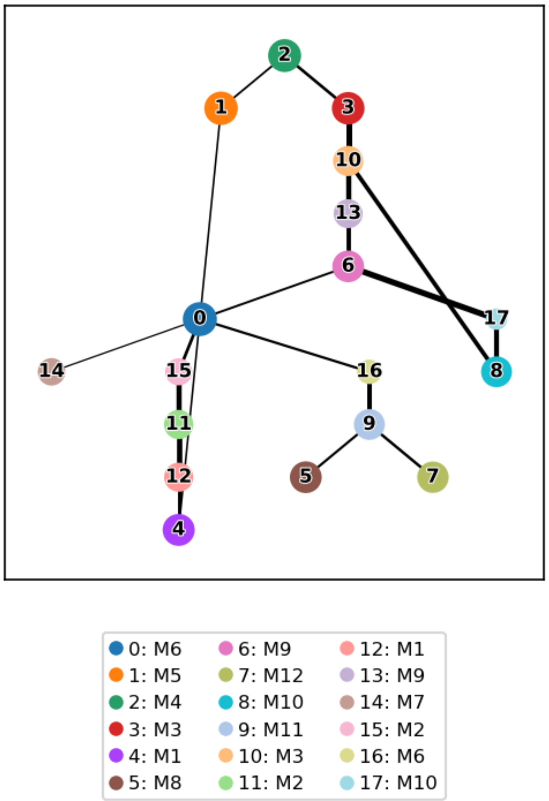

Figure 26: *Dyntoy* dataset subsection 1.1

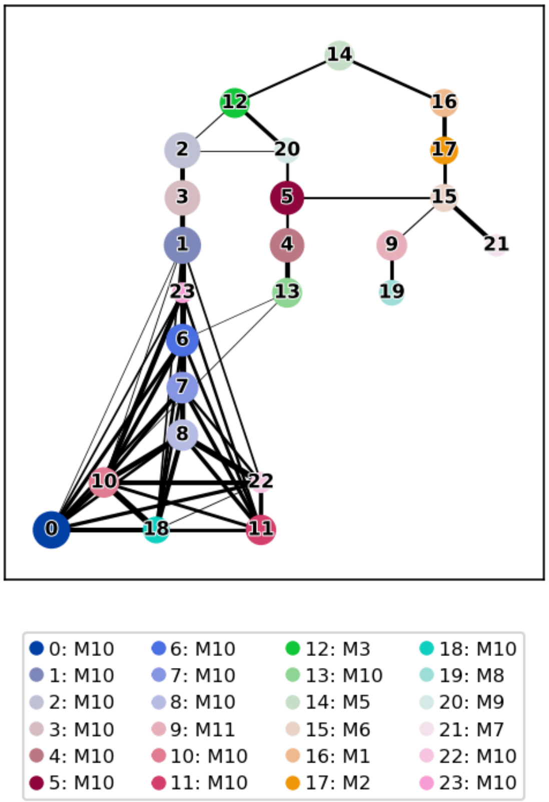

Figure 27: Up-sampled *dyntoy* dataset subsection 1.1

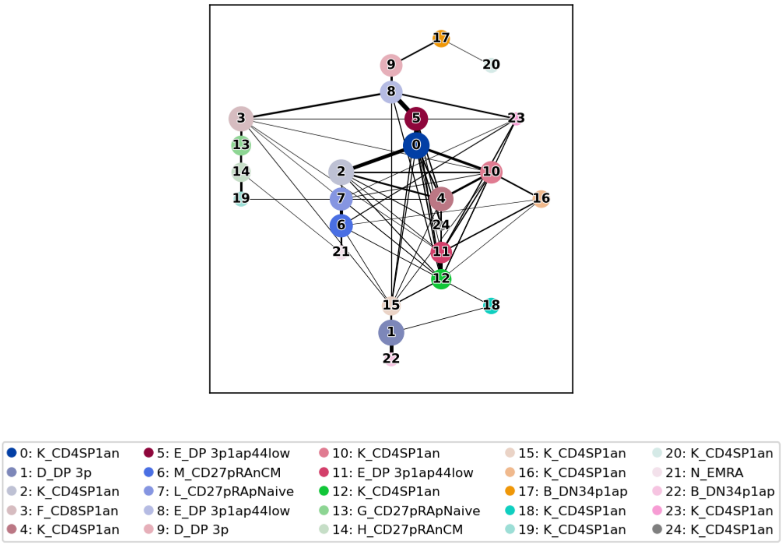

Figure 28: Human thymus & PBMC subsection 1.2

##### 8.2.7 CellRank 2

*CellRank 2* Weiler et al. (2024) correctly detected endpoints in all tested samples, although it also identified some additional endpoints that do not appear to be valid. However, the detection of subtle trajectories (e.g., beta selection, Figure 31) or the proper analysis of converging trajectories (see subsection 10.1, Figure 48) remains challenging for this method.

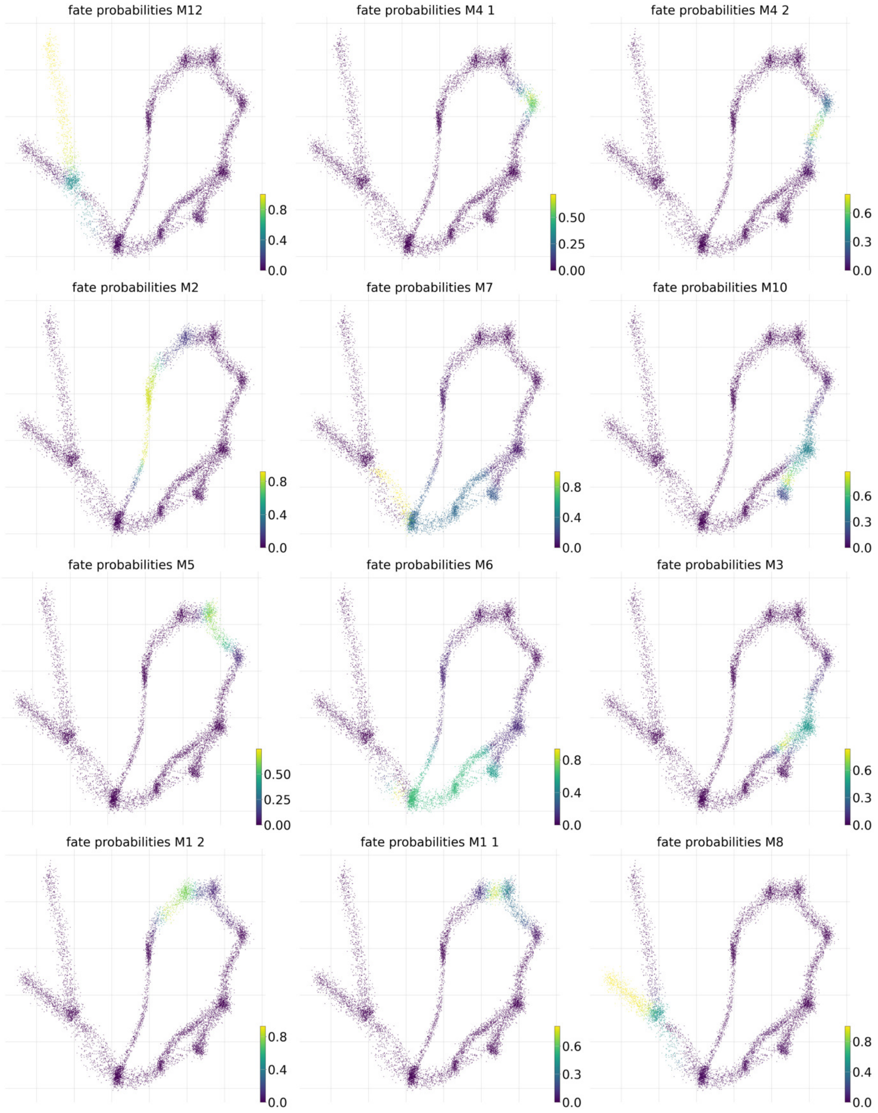

Figure 29: *Dyntoy* dataset subsection 1.1

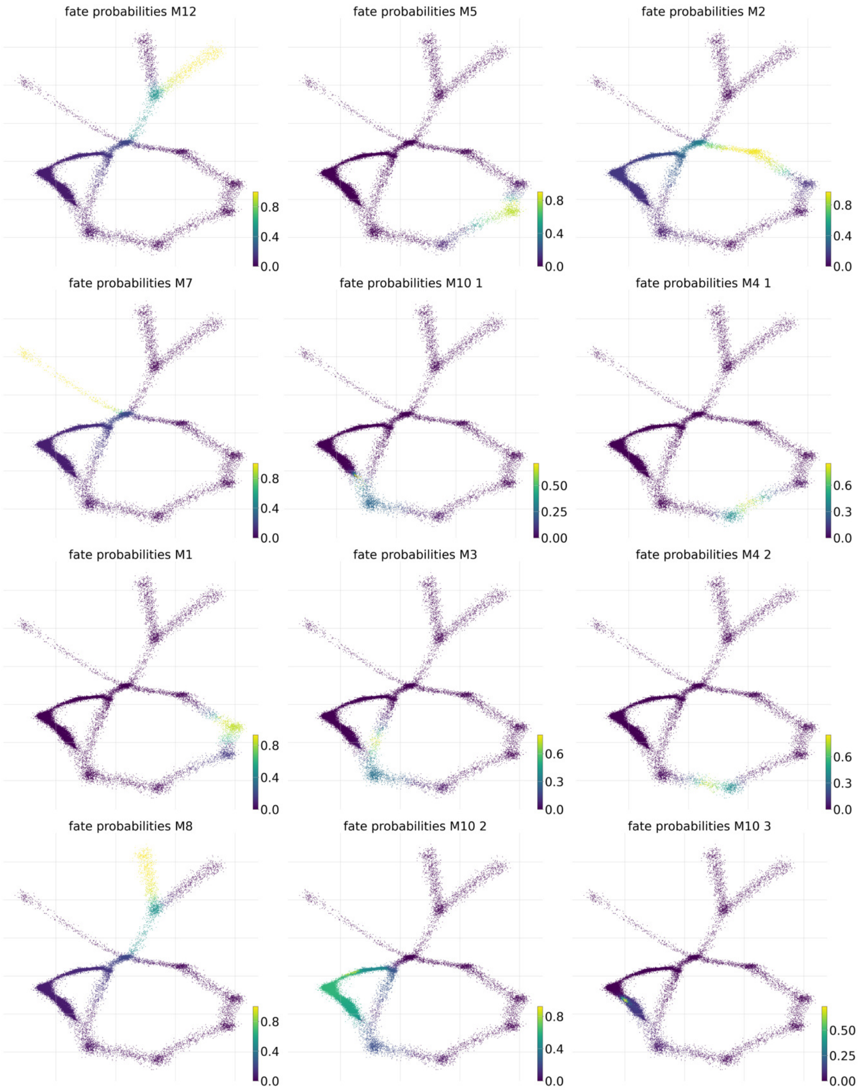

Figure 30: Up-sampled *dyntoy* dataset subsection 1.1

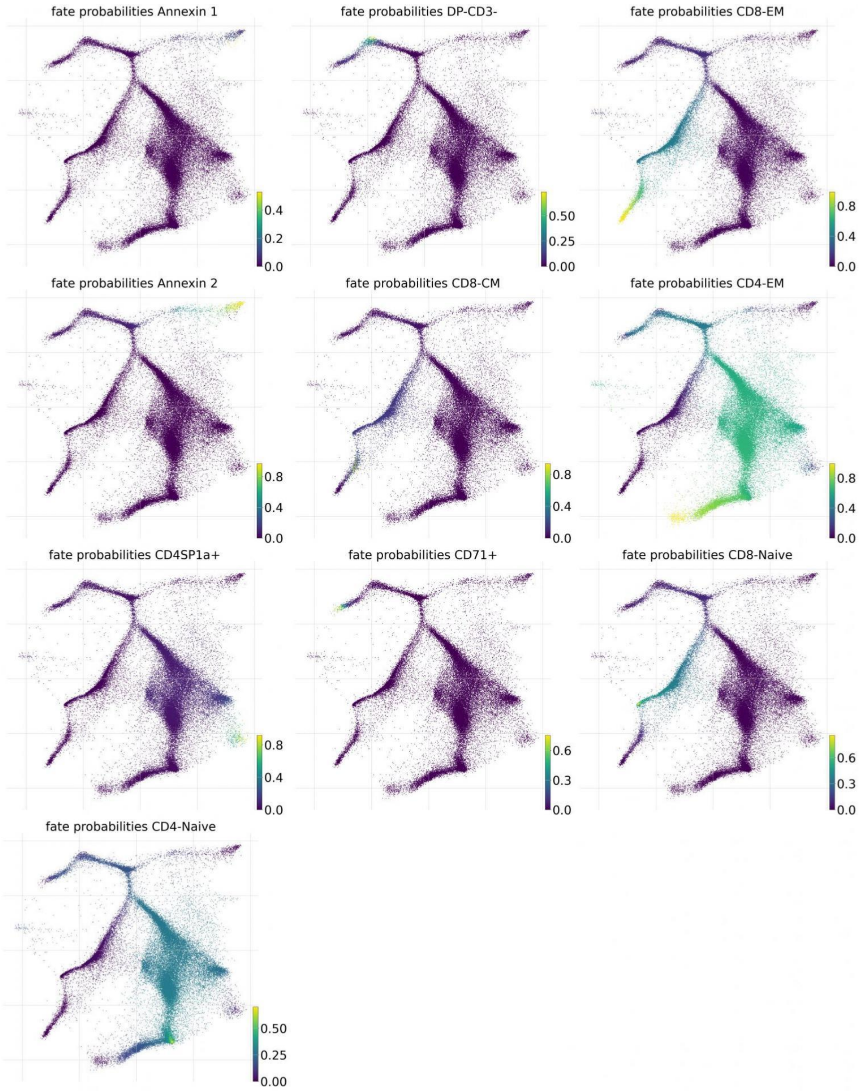

Figure 31: Human thymus & PBMC subsection 1.2

###### 8.2.8 StaVia

New version of *VIA*, *StaVia* Stassen et al. (2024), performed well on artificial data. However, on thymus data, the large number of suggested trajectories makes the results difficult to interpret. Similarly to other methods, StaVia missed the beta selection checkpoint.

Admittedly this method was designed rather for scRNAseq data than for mass cytometry. We have therefore compared *StaVia* to *tviblindi* on 2 scRNAseq datasets in subsection 10.1 and subsection 10.2. *StaVia* performed well, but it still struggles to separate converging trajectories even with the use of *random walks with memory* specifically designed to address this issue (see Figure 48).

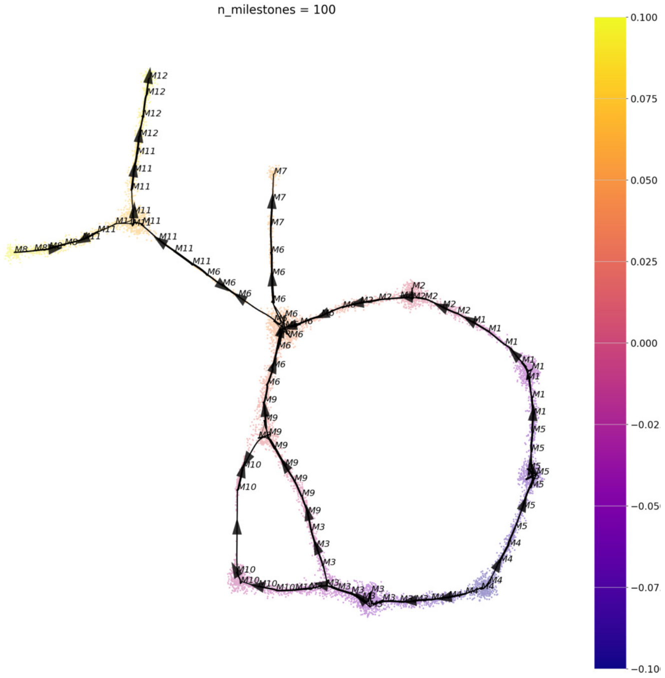

Figure 32: *Dyntoy* dataset subsection 1.1

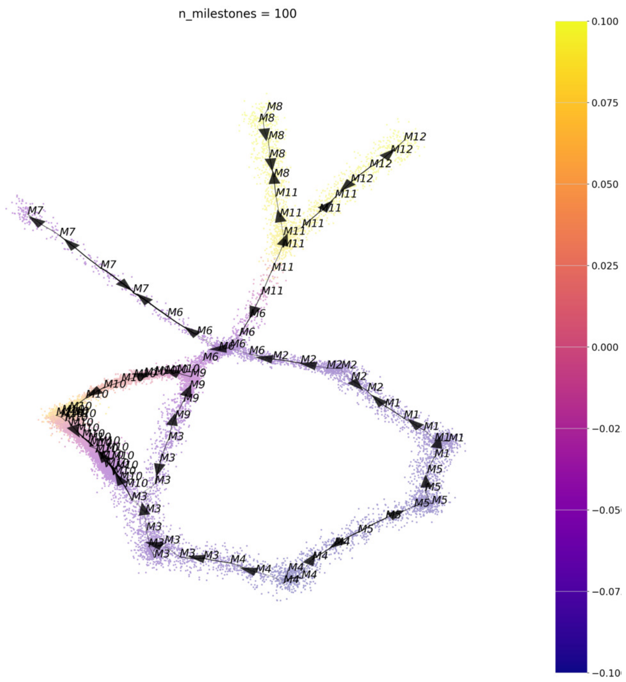

Figure 33: Up-sampled *dyntoy* dataset subsection 1.1

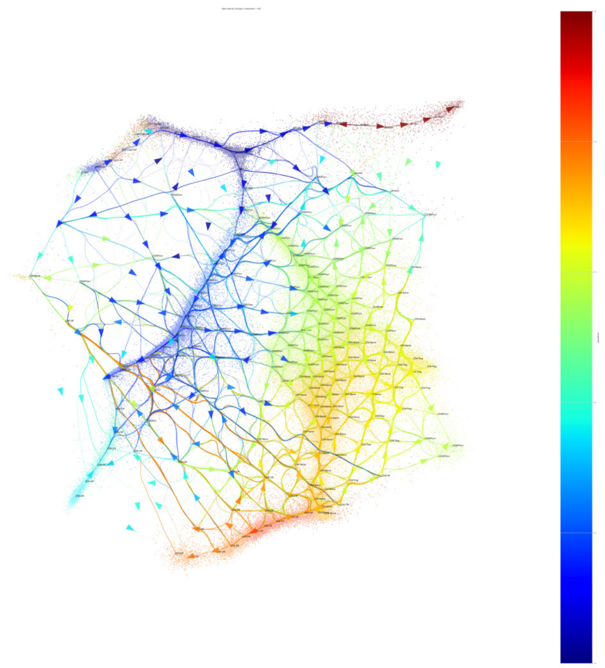

Figure 34: Human thymus & PBMC subsection 1.2

#### 8.3 Running times

*tviblindi* outperforms tested TI methods on medium-sized and large datasets (Table 1) in terms of running time. All methods were run in a dedicated Docker container with 20 CPUs and RAM limited to 200 GB.

The current implementation of *tviblindi* has effectively linear complexity with respect to the number of events for datasets with less then $10^7$ events Figure 35. There is no clear theoretical reason for the complexity to be super-linear and we are currently focusing on the implementation to pinpoint possible overhead.

Table 1: Running times (minutes). Evaluated on the dataset from subsection 1.2.

| n cells | tviblindi | VIA | Palantir | Monocle 3 | STREAM | PAGA | StaVia | CellRank |
|---|---|---|---|---|---|---|---|---|
| **50000** | 0.9 | 0.8 | 3.2 | 4.9 | 10.7 | 0.8 | 1.9 | 9.48 |
| **100000** | 1.7 | 1.5 | 6.5 | 20.4 | 20.8 | 2 | 3.1 | 15.95 |
| **200000** | 4.1 | 3.0 | 14 | 74 | 32.7 | 6 | 6.0 | 26.21 |
| **500000** | 9.7 | 10.6 | 33.4 | X | 62 | 20.4 | 17.4 | 73.84 |
| **All (1,182,80 2)** | 22 | 24.4 | 87.9 | X | 136.7 | 46.1 | 49.7 | 167.15 |

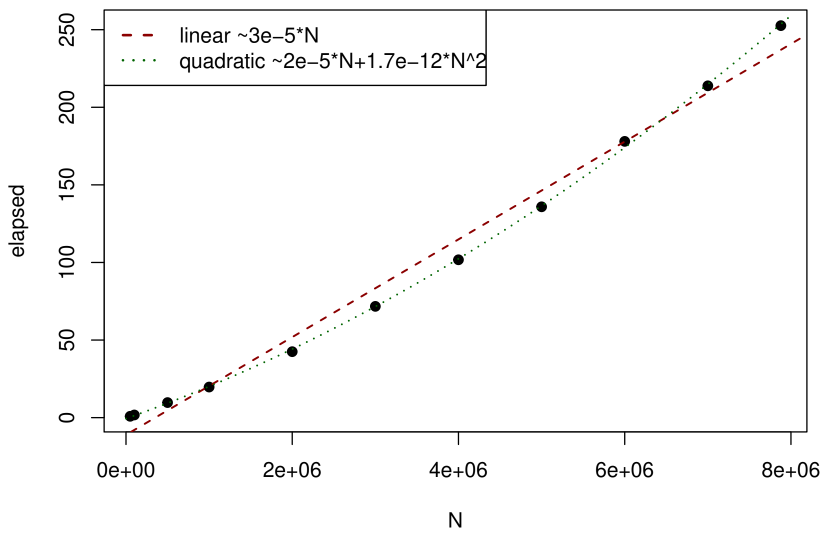

Figure 35: Running times of *tviblindi* with respect to the number of events (minutes).

### 9 Analysis of human thymus single-cell RNA-seq data

In this section we present analysis of the human thymus scRNA-seq dataset (subsection 1.3) using *tviblindi*, which led to the results presented in the Figure 7 of the main text. Imputed[7] data are used as the input for the trajectory inference, scaled counts (no imputation) are

[7]Here, we chose to use data imputed by *Rmagic* to highlight the ability of *tviblindi* to operate directly in high-dimensional space. In general, such strong imputation is not advised, and the use of principal component analysis (PCA) is preferred. However, in this case, we obtained equivalent (see Figure 36) results when using the first 50 principal components (PCs).

shown in line plots.

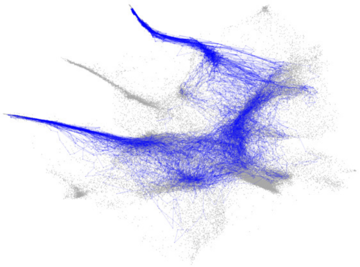

(a) Selected trajectories

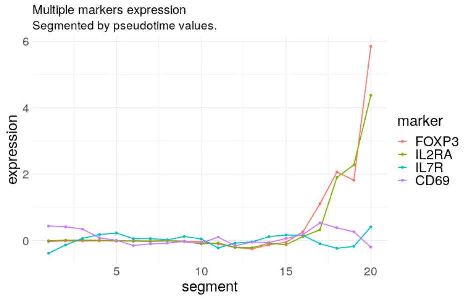

(b) Markers 1

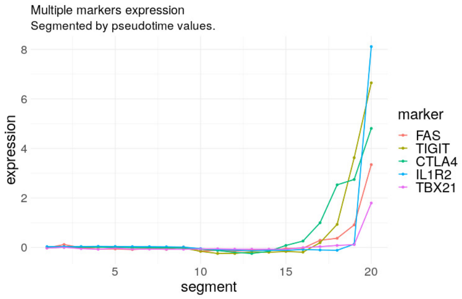

(c) Markers 2

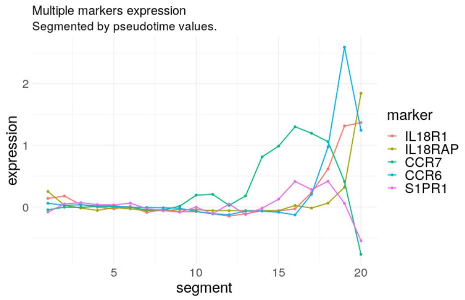

(d) Markers 3

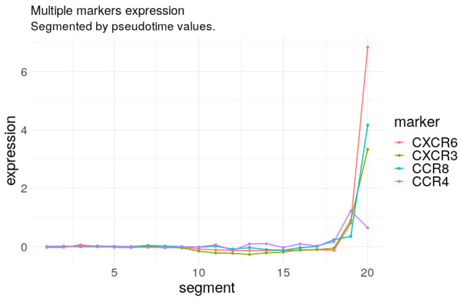

(e) Markers 4

Figure 36: Analysis using the first 50 principal components; the Vaevictis projection is computed on imputed data for straightforward comparison

The outline of the analysis is presented in Figure 37. Double negative - DN (early) - population is set as origin, endpoints corresponding to Treg population are selected (Figure 37a) and significant sparse regions are chosen (Figure 37b) resulting in the dendrogram of random walks (Figure 37c). The dendrogram suggests two distinct groups of trajectories (marked by blue and red rectangles). The blue trajectory is analyzed in detail within the main text. Nonetheless we cannot simply neglect the red one. Although the *persistent homology* suggests a significant difference between these two trajectories, there is no visible distinction on the *vaevictis* plot (Figure 37d). We calculated several[8] alternative *vaevictis* projections, which confirmed that the trajectories are distinct (Figure 37e, Figure 37f). This distinction is most likely a batch effect artifact rather than a biological feature as the main difference observed at the terminal stage of the development turned out to be in the sample source of the cells (compare Figure 37f and Figure 38).

The evolution of the analyzed markers on the red trajectory is consistent with the evolution represented on the blue one. The red trajectory contains a comparatively smaller number of events with the highest pseudotime. The resulting line plots do not capture the pattern at the very end of development (Figure 39).

[8]There is no clear guidance on how to set up *vaevictis* or other dimensionality reduction tools to discover this kind of features. In this particular case increasing the number of clusters for importance sampling from 15 to 35 worked, but the outcome depends on the clustering method. The result confirms the power of topological data analysis.

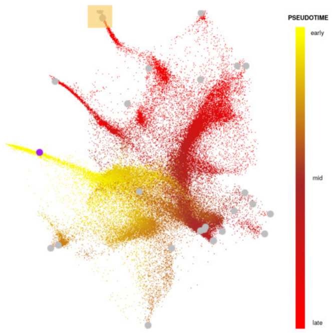

(a) Pseudotime & Termini

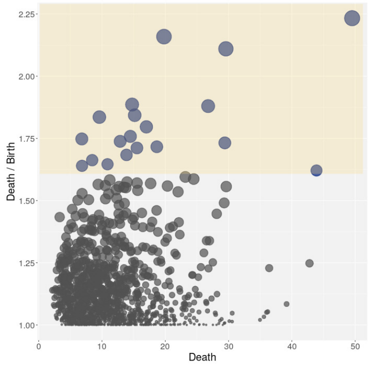

(b) Persistence diagram

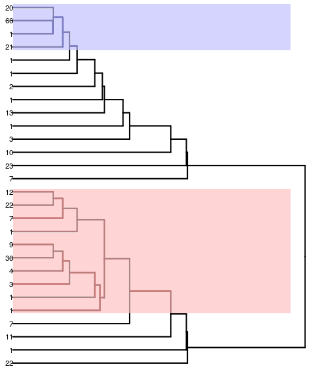

(c) Dendrogram of random walks

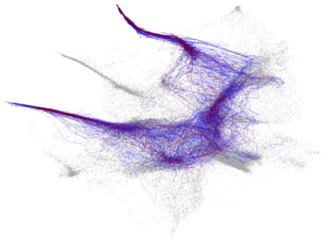

(d) Selected trajectories

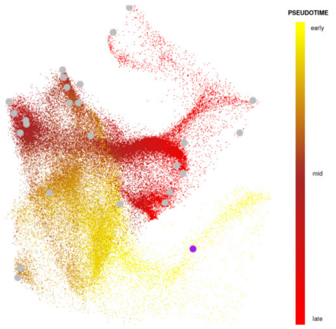

(e) Alternative *vaevictis* projection

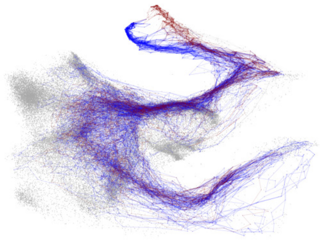

(f) Selected trajectories

Figure 37

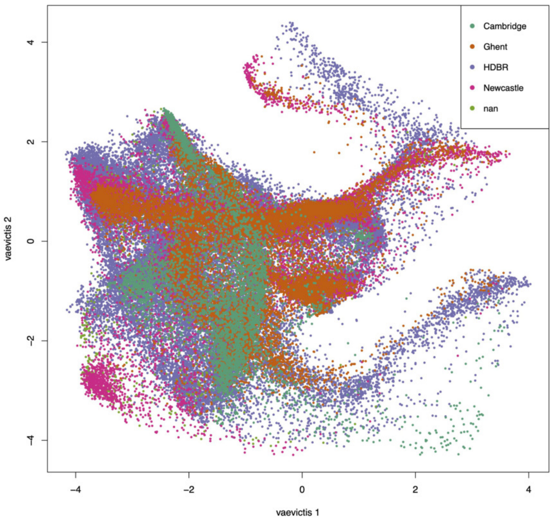

Figure 38: Alternative *vaevictis* projection, colored by sample source

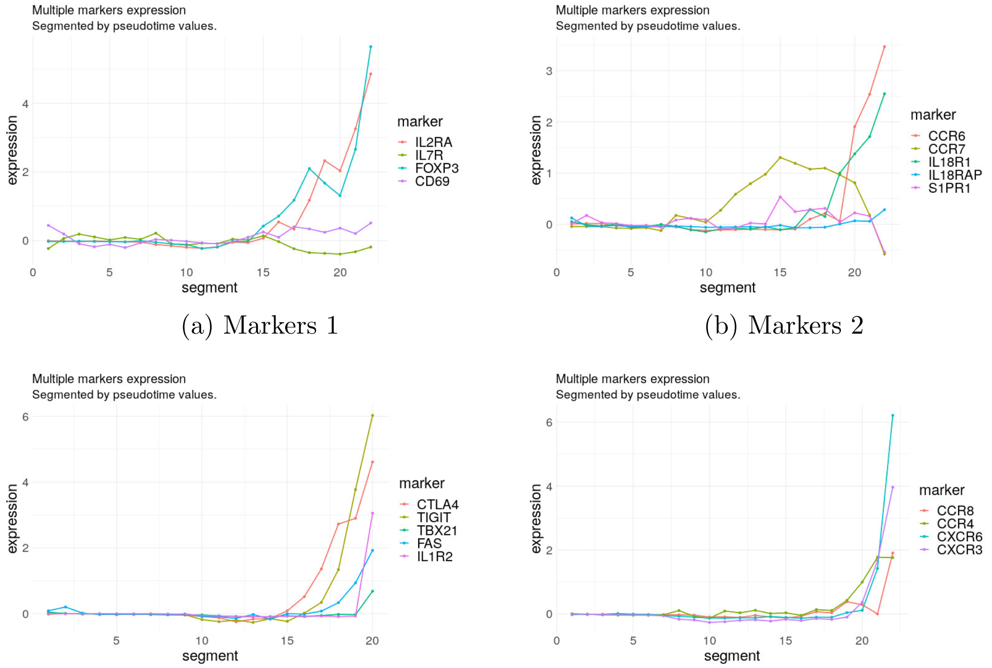

(a) Markers 1

(b) Markers 2

(c) Markers 3

(d) Markers 4

Figure 39: Pseudotime line plots of the red trajectory

### 10 Evaluation of *tviblindi* on scRNAseq data

#### 10.1 *tviblindi* evaluation on Bone marrow atlas dataset

```
tv1<-tviblindi(data=as.matrix(BMlist$integrated_pca),
     labels=list(lab1=BMlist$meta$Level.3.Multimodal,
     lab2=BMlist$meta$Level.1,lab3=BMlist$meta$Level.2,
     lab4=BMlist$meta$Level.3.RNA))
Set_origin(tv1,label = "HSC-1")
KNN(tv1,method = "balltree")
Denoise(tv1)
Cluster(tv1) #kmeans clustering
Filtration(tv1)
DimRed(tv1,upsample = list(N=1500), use.denoised = TRUE)
DimRed(tv1, use.denoised = TRUE)
Pseudotime(tv1)
Walks(tv1,N=10000)
```

In this section we analyze the scRNaseq dataset subsection 1.4 using *tviblindi* and compare the results to selected state-of-the art methods.

See the code used for the analysis above. When working with scRNA-seq data, we typically use exact $k$-NNG calculation (`method="balltree"`) via the vantage-point tree algorithm, as the number of events is usually smaller. Additionally, for the two-dimensional representation, it is preferred to base the calculation on denoised data[9] (`use.denoised=TRUE`). See Figure 49 for comparison. Finally, we can leverage the ability of *vaevictis* to integrate annotation information using `upsample=list(N=1500)`. This overrides the default impor-

[9]Recall that dimensionality reduction is used for visualization purposes only

tance sampling based on clustering, and the representatives are sampled according to the annotation in the first label set (`lab1`). To compare with *vaevictis* run using the default setting `DimRed(tv1, use.denoised=TRUE)`, see Figure 41. The hematopoetic stem cells (`HSC-1`) population is fixed as the origin.

See the basic overview in Figure 40. We begin the analysis (see Figure 42 for a detailed workflow) by selecting the candidate endpoints. The selected endpoints are highlighted in blue text (Figure 42a). For the sake of clarity, we exclude cells annotated as NK/T (black rectangle) from the analysis, as T cells are not directly developmentally connected to bone marrow precursors, making the interpretation of results difficult. We select all highlighted endpoints[10] simultaneously, which yields essential homological classes corresponding to the separation of endpoints (Figure 42b, red rectangle). Next, we select all major leaves on the dendrogram (Figure 42c) and plot the respective trajectories (Figure 43 and Figure 44).

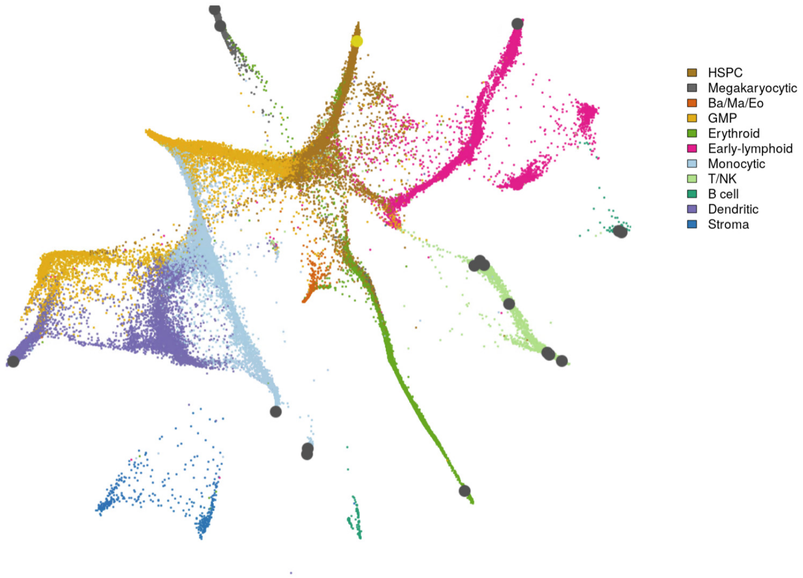

Figure 40: *vaevictis* projection colored by population

[10] We do not select the monocytic endpoint below the blue text at this stage, as the resulting dendrogram would be too complex.

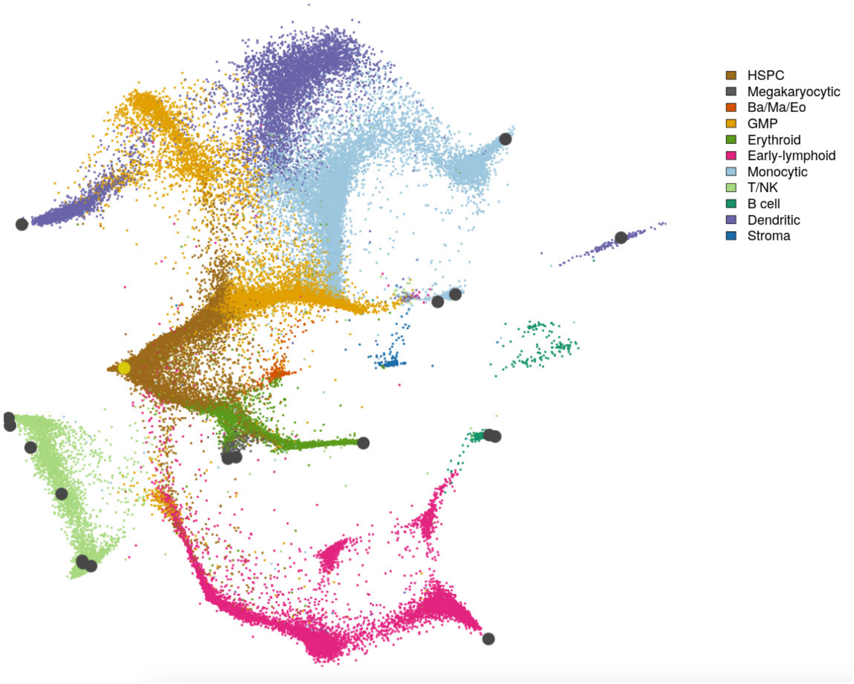

Figure 41: *vaevictis* projection colored by population - default setting without cell annotation information

We notice that one of the trajectories selected on the dendrogram is heterogeneous and focus on it (Figure 48). Finally, we examine the second monocytic endpoint (Figure 46). Most of the trajectories correspond to those already investigated on the previous dendrogram; however, one trajectory corresponds to the development of the basophil/eosinophil/-macrophage population, which is connected (possibly artifactually) to this population on the $k$-NNG. *tviblindi* enables the identification of such connections and facilitates their correct interpretation.

We now compare the results to *CellRank 2* (Figure 47) and *StaVia* (Figure 49). Both tools performed well in identifying the correct endpoints. However, they do not provide a means to adjust the resolution or dissect compound trajectories. For example, consider the development of plasmacytoid dendritic cells. It is well known that these cells can

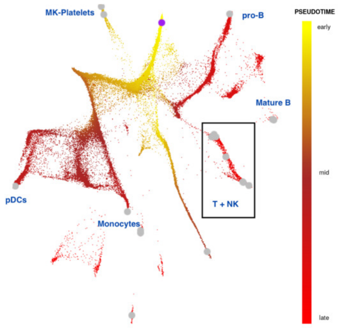

(a) Pseudotime & Termini, T + NK (black rectangle) excluded from further analysis

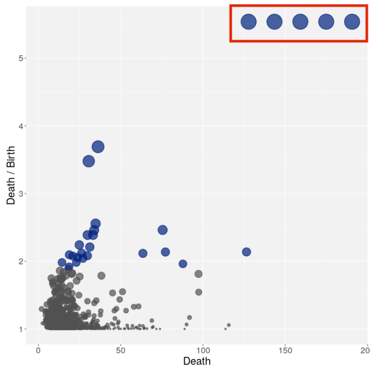

(b) Persistence diagram with essential classes in red rectangle

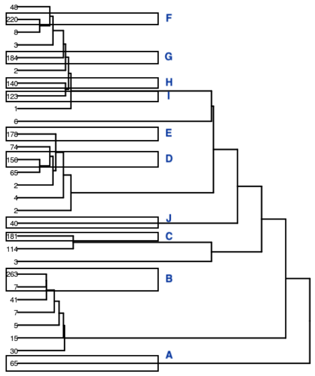

(c) Dendrogram of random walks

Figure 42

develop via multiple pathways, such as from lymphoid and myeloid precursors (Reizis (2019)). Both tools correctly identify the potential precursors but fail to separate the respective trajectories (Figure 48). *tviblindi*, on the other hand, suggests multiple coherent trajectories.

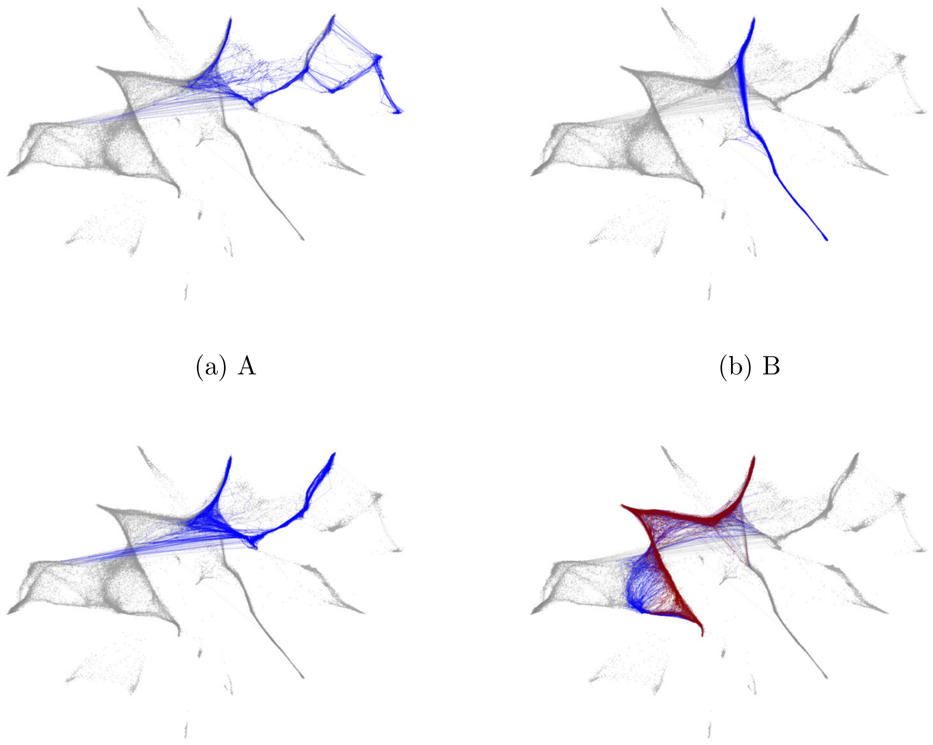

(a) A

(b) B

(c) C

(d) D+E

Figure 43

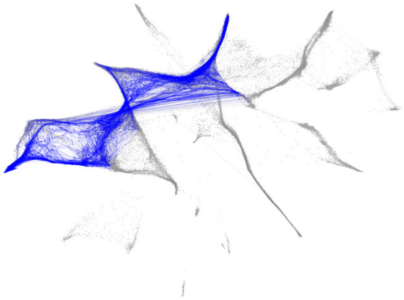

(a) F

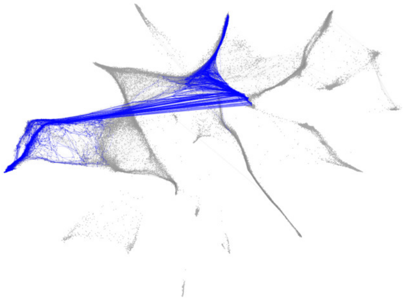

(b) G

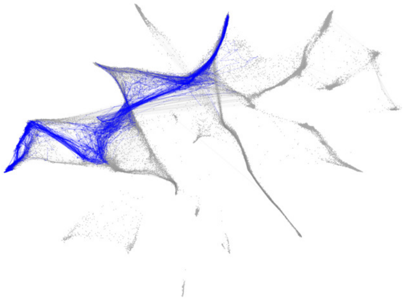

(c) H

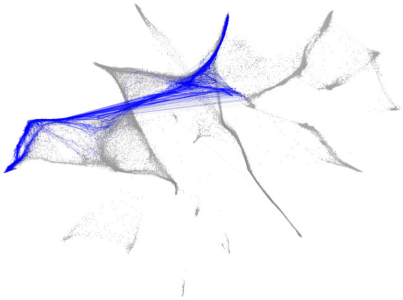

(d) I

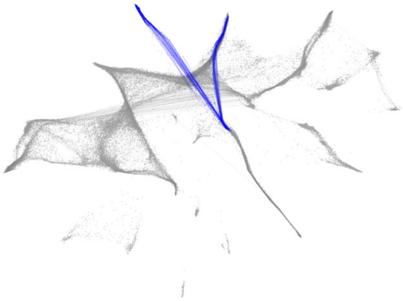

(e) J

Figure 44

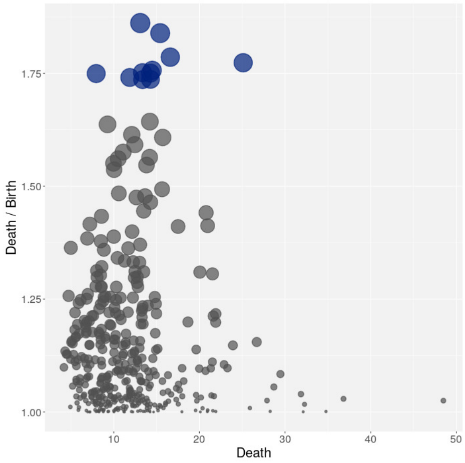

(a) F - Persistence diagram

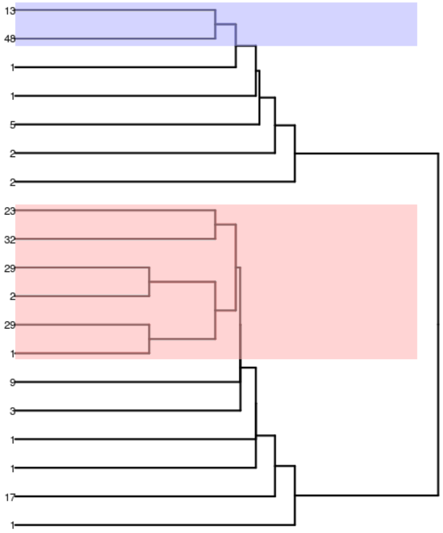

(b) F - dendrogram

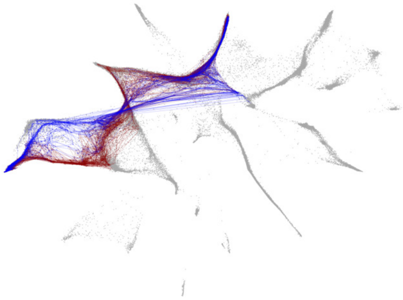

(c) F - 2 alternatives

Figure 45

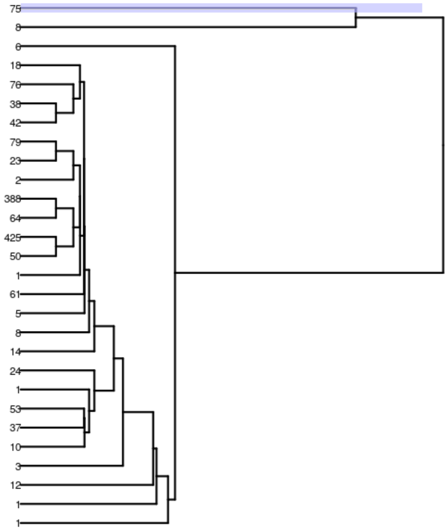

(a) Non-classical monocytes dendrogram

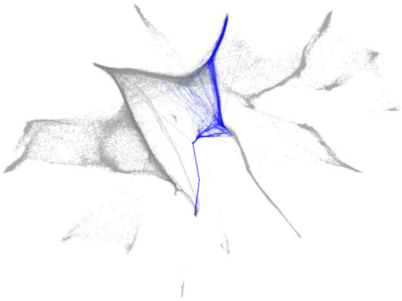

(b) Non-classical monocytes - trajectory through BaMaEo

Figure 46

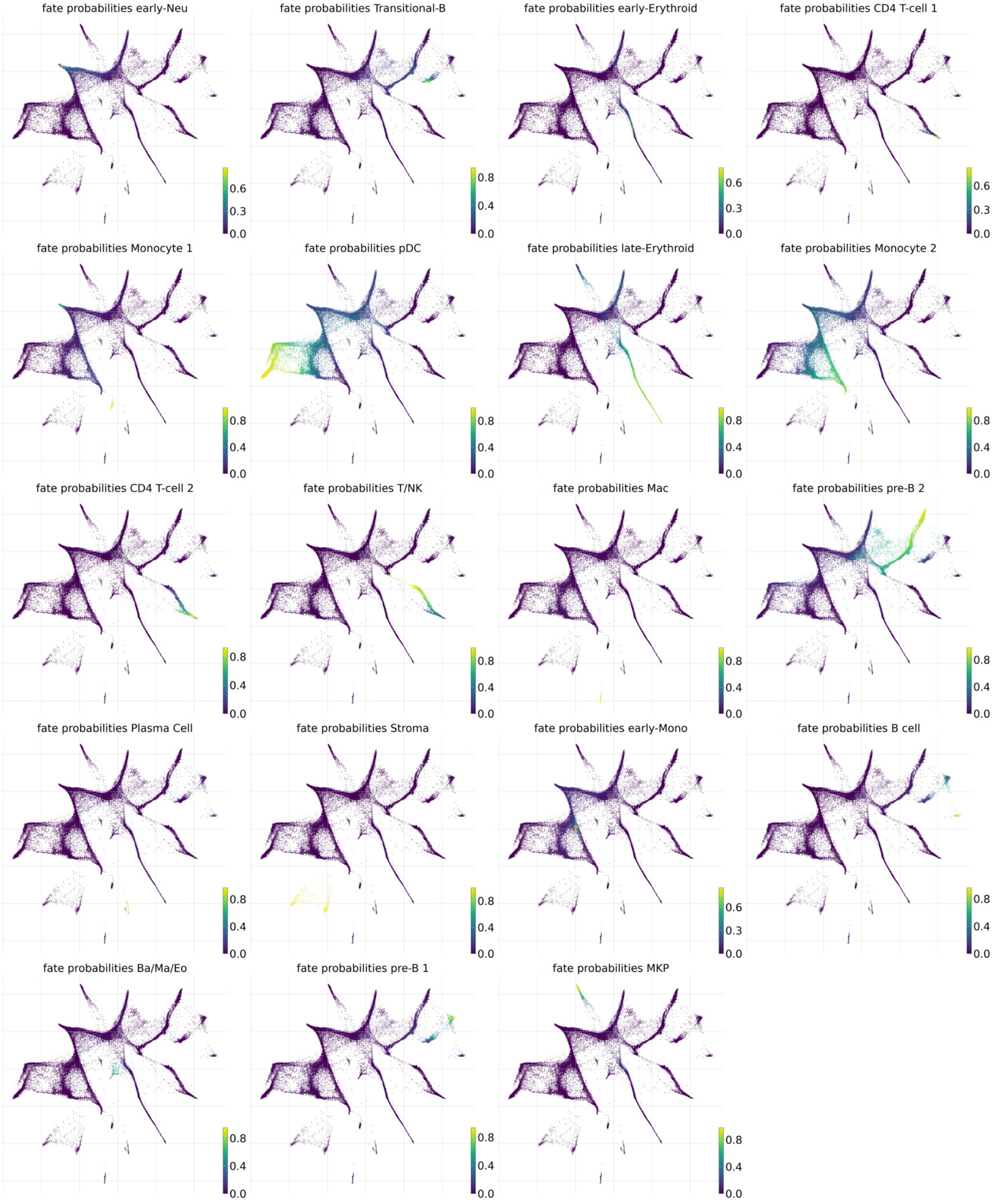

Figure 47: Bone marrow data subsection 1.4 - fates detected by CellRank 2

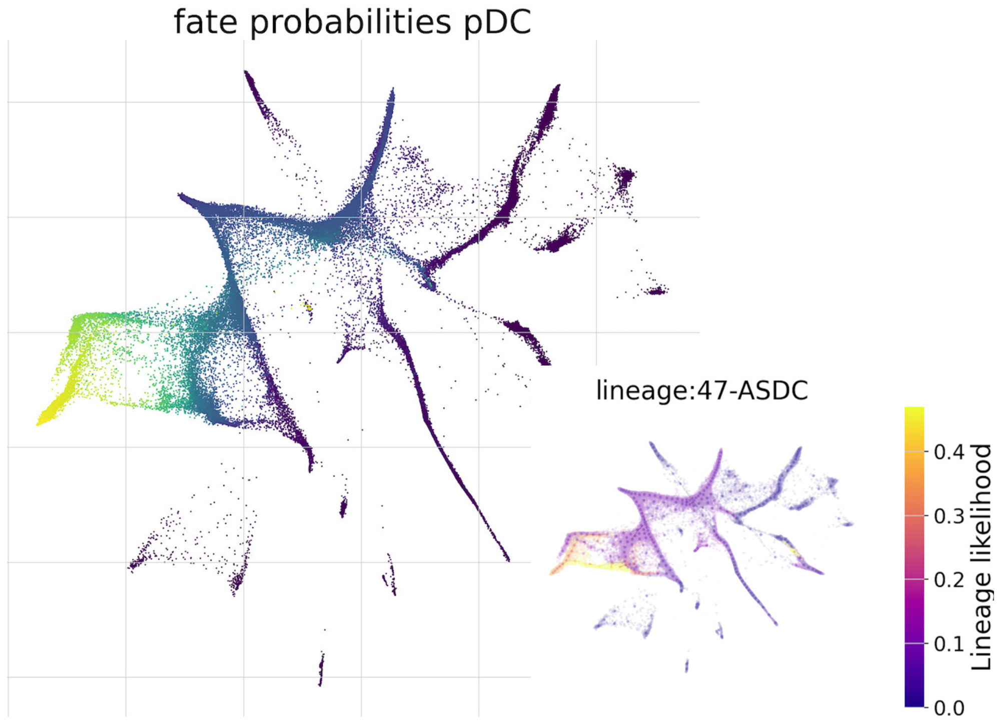

(a) CellRank2 - fate probabilities (b) StaVia - lineage likelihood

Figure 48: StaVia and CellRank2 - pDCs fate

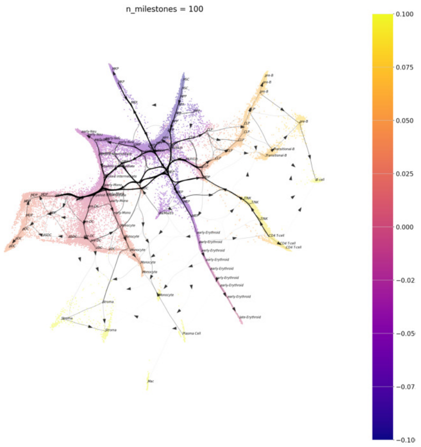

(a) *vaevictis*

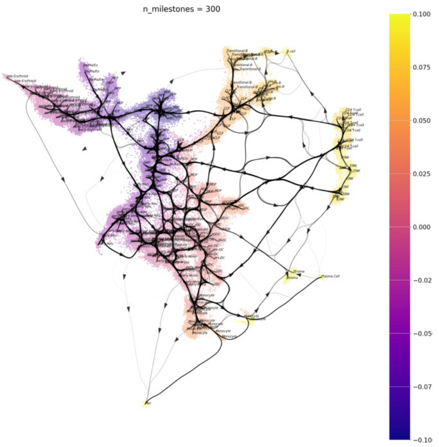

(b) UMAP

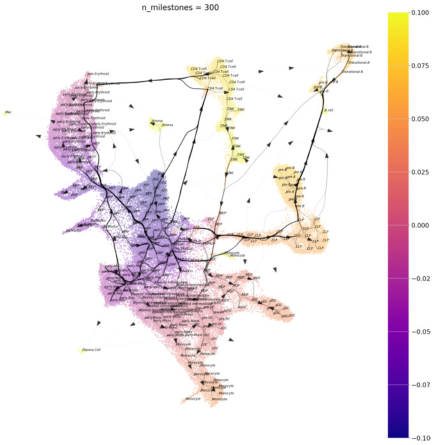

(c) StaVia

Figure 49: Comparison of different projections with *StaVia* atlas view overlay

#### 10.2 *tviblindi* evaluation on Mouse gastrulation dataset

```
tv1<-tviblindi(data=as.matrix(MGlist$pca),
     labels=list(stage=MGlist$meta$stage,
     celltype=MGlist$meta$celltype))

Set_origin(tv1,label = "Epiblast")
KNN(tv1,method = "balltree")
Denoise(tv1)
Cluster(tv1) #kmeans clustering
Filtration(tv1)
DimRed(tv1,upsample = list(N=1500), use.denoised = TRUE)
DimRed(tv1, use.denoised = TRUE)
DimRed(tv1,layout = MGlist$umap)
Pseudotime(tv1)
Walks(tv1,N=5000)
```

In this section we analyze the scRNaseq dataset of mouse gastrulation data (subsection 1.5) using *tviblindi*. This dataset was analysed using *StaVia* and *CellRank 2* in Stassen et al. (2024).

See the code used for the analysis above. When working with scRNA-seq data, we typically use exact $k$-NNG calculation (`method="balltree"`) via the vantage-point tree algorithm, as the number of events is usually smaller. Additionally, for the two-dimensional representation, it is preferred to base the calculation on denoised data[11] (`use.denoised=TRUE`). See Figure 49 for comparison. Finally, we can leverage the ability of *vaevictis* to integrate annotation information using `upsample=list(N = 1500)`. This overrides the default im-

[11] Recall that dimensionality reduction is used for visualization purposes only

portance sampling based on clustering, and the representatives are sampled according to the annotation in the first label set (`stage`). The `Epiblast` is fixed as the origin.

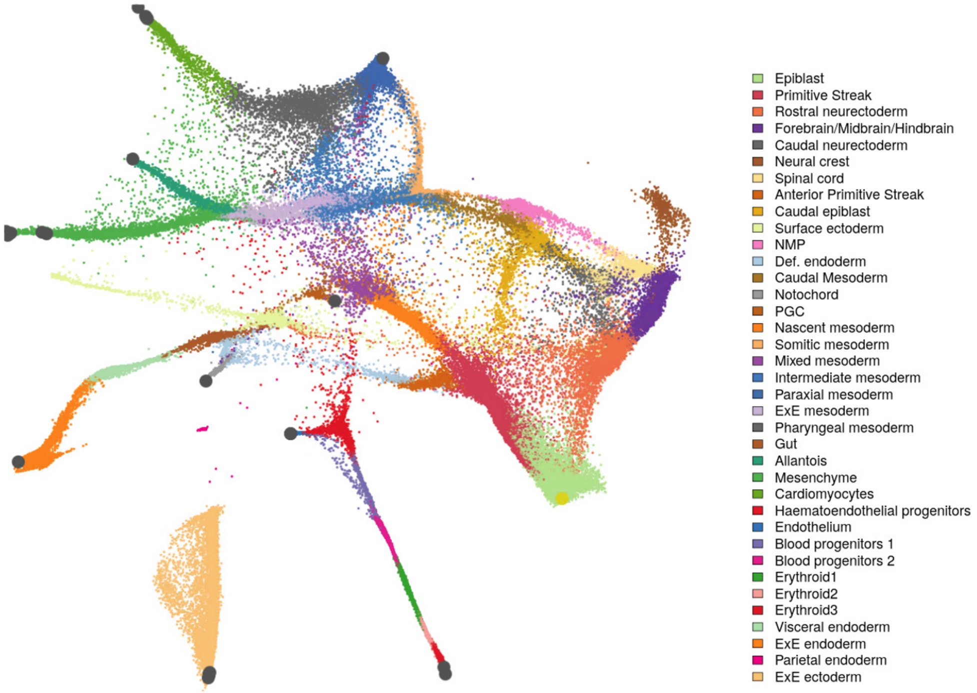

Figure 50: *veavictis* projection colored by population

See the basic overview in Figure 50. As the gastrulation atlas is an extremely complex dataset, we present two examples of development: one toward the extra-embryonic endoderm to highlight a pattern of multiple sources for a particular cell population (in this case, the gut), and another toward cardiomyocytes to illustrate a different pattern of multiple sources.

For the first pattern, see Figure 51. The extraembryonic endoderm is selected as an endpoint. The majority of the trajectories correspond to the development of the embryonic gut. This pattern is usually easy to identify and *StaVia* did not have any issues with it.

Another pattern, similar to the one observed in the development of plasmacytoid den-

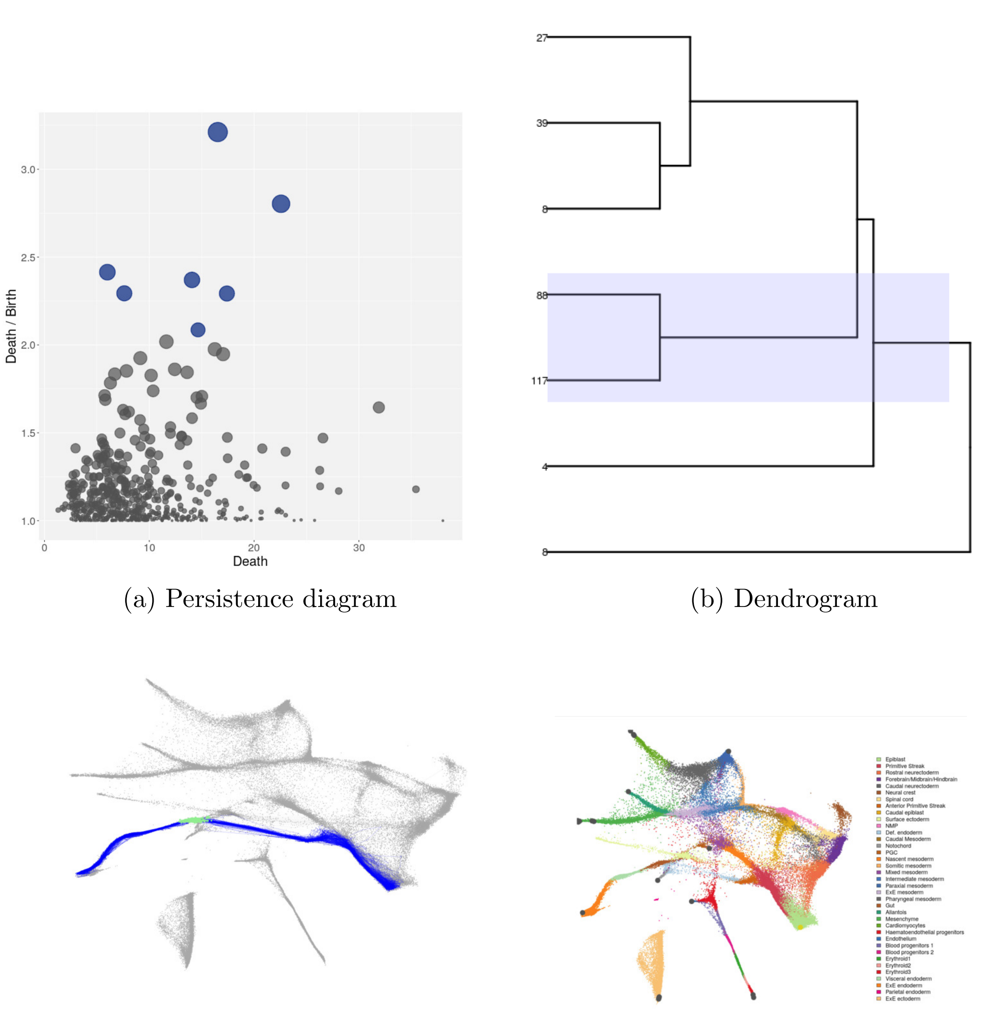

(a) Persistence diagram (b) Dendrogram

(c) Selected trajectories (d) Cell populations

Figure 51: Connection between extra-embryonic endoderm and epiblast, highlighting multiple precursors of the embryonic gut. Gut cells on the trajectory are highlighted.

dritic cells, arises when analyzing the development of cardiomyocytes. It is known that cardiomyocytes can develop through distinct pathways - one via hematoendothelial progenitors and mesoderm, and another via the neural crest (George et al. (2020)). *tviblindi* facilitates the identification and dissection of these pathways (see Figure 52). Note that we have selected a heterogeneous branch on the dendrogram (Figure 52b, red group), and

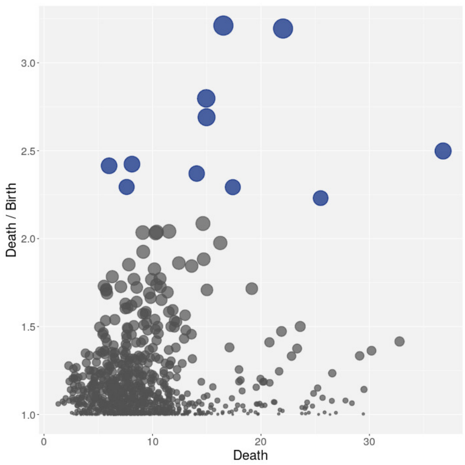

(a) Persistence diagram

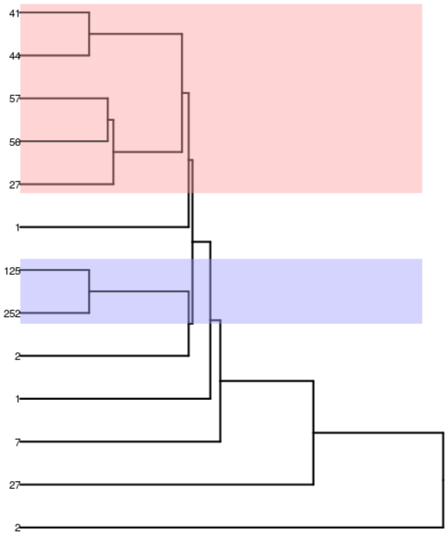

(b) Dendrogram

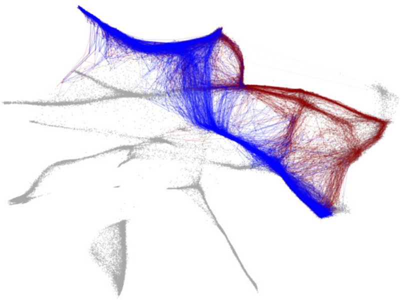

(c) Selected trajectoris

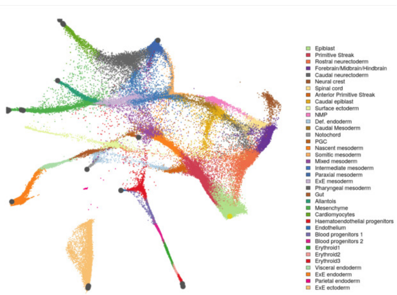

(d) Cell populations

Figure 52: The development trajectories to cardiomyocytes

finer separation could easily be achieved.

#### 11 Software acknowledgement

Apart from tools directly referenced in the text, the following tools and libraries were used in the implementation of *tviblindi*: The CGAL Project (2019); Seel (2019); Bauer et al.

(2017); Bates et al. (2023); Eddelbuettel et al. (2023b); Bates and Eddelbuettel (2013); Kulichova and Kratochvil (2023); Eddelbuettel et al. (2023a); Bailey (2022); Perrier et al. (2023); Pedersen et al. (2022); Wickham et al. (2019); de Vries and Ripley (2022); Galili (2015); Wickham and Seidel (2022); Warnes et al. (2022); Urbanek (2022); Csardi and Nepusz (2006); Eddelbuettel and Sanderson (2014); Wilke (2020); Sali and Attali (2020); Maechler et al. (2022); Meyer and Perrier (2022); Melville (2023); Ellis et al. (2023); Huber et al. (2015); Ushey et al. (2023)
